# Genetic mapping of Dutch elm disease resistance

**DOI:** 10.64898/2026.04.28.721446

**Authors:** Catherine Gudgeon, Mohammad Vatanparast, Rômulo Carleial, David Herling, Fergus Poncia, Alberto Santini, Clive Brasier, Joan F. Webber, Richard Buggs

## Abstract

Dutch elm disease (DED) has killed millions of field elms (*Ulmus minor*) in Britain and Europe since the 1970s, causing incalculable damage to landscapes and their associated biodiversity. While the species *U. minor* is highly susceptible to DED, some east Asian and Himalayan species of elm are resistant. These Asiatic species differ in their growth and form to *U. minor* and cannot fully substitute for it in the landscape. Several breeding programmes have attempted to generate trees that combine DED-resistance with the growth and form of *U. minor* via hybridisation and back-crossing. These have been partly successful, but further breeding is needed to fully realise these ambitions. Most recently in Britain, a complex resistant hybrid ‘Wingham’ (FL493) was crossed with a surviving *U. minor* tree in Tonge Mill, Kent. Sixty progeny from this cross were tested for field resistance to DED and show segregation for this trait. Here, we analyse the genome of these progeny in order to: (1) construct a linkage map of the elm genome, and (2) identify regions of their genome derived from east Asian and Himalayan species that could be associated with DED-resistance. Such knowledge could accelerate future breeding programmes and enhance our understanding of the mechanistic basis of resistance.

## Introduction

Since 1970 Dutch elm disease (DED) has killed an estimated 60-100 million elms in Britain, with the fungal pathogen *Ophiostoma novo-ulmi* spread almost exclusively by two species of *Scolytus* bark beetles which feed on healthy elms and use dying elms as breeding material (Potter et al. 2011; Gibbs and Howell 1972, 1974; West et al. 2025; Brasier and Buck 2001). Nowadays, few mature elms remain in the UK landscape although rare old survivors do occur, some of which may have a degree of natural field resistance (Shreeve and Seddon 2024). Young elms regenerate constantly in the UK and tend to survive 10-20 years before succumbing to DED (Brasier and Webber 2019; Peace 1960). The availability of regenerating elms means that the smaller vector beetle, *Scolytus multistriatus* can breed successfully but the larger *S. scolytus*, which is a more effective DED vector, requires larger elms with thicker bark to reproduce and may therefore have become more scarce (Webber 2000).

Generally, European elms have little genetic resistance to *O. novo-ulmi,* although some survive in the field because they are unpalatable to the beetles (Webber 2000). Several breeding programmes in Europe have crossed resistant Asiatic elms with European species to produce *O. novo-ulmi* resistant cultivars (Smalley and Guries 2000; Martín et al. 2019; Mittempergher and Santini 2004) although none phenotypically resemble the field elms (*Ulmus minor*) that were once a familiar sight in Britain. Therefore, the development of a DED-resistant cultivar that retains the overall phenotypic characteristics of native British elms would be of significant value for restoration initiatives and positively influence public perception (Russell and Buggs 2019).

Little is known about the genomic basis of resistance to DED in Asiatic species of elm or in complex resistant hybrids made by crossing and back-crossing them with *U. minor* and/or *U. glabra*. Most resistant cultivars were made before genomic technologies were available to analyse them. In recent years, most elm breeding programmes have ceased. One exception is late David Herling’s work in Britain, who crossed a resistant cultivar FL493 (*Ulmus* ‘Wingham’) with a surviving *U. minor* from Tonge Mill in Kent. Here, we report his inoculation trials and whole-genome resequencing to analyse 60 progeny of this cross (which we refer to as the WxTM progeny throughout the paper) in order to shed light on the genomic basis of DED resistance.

## Methods

### Biological materials

The majority of the samples were the result of a cross conducted by David Herling, yielding a full-sibling family of 60 progeny bred via controlled pollination between a putative field resistant *U. minor* specimen in Tonge Mill, Kent (Figure 1) and a clone of FL493 (*Ulmus* ‘Wingham’) obtained in Florence (Italy) at the C.N.R. - Institute for Sustainable Plant Protection) within the elm breeding program (Santini et al., 2008). The WxTM cross progeny are growing at a trial site in Wateringbury, Kent, where they were planted in early 2016. ‘Wingham’ is an elm hybrid resulting from a 4-way genetic admixture between *U. minor*, *U. glabra*, *U. pumila*, and *U. wallichiana* (Figure 1), the former two species being native to Europe with high susceptibility to DED, and the latter two being Asiatic species with high DED resistance. We also obtained 5 different representative samples for each of these ‘parental’ species, with the exception of *U. wallichiana* for which only one sample was available. We also obtained a DNA sample from Hans Heybroek’s (Heybroek 1993) clone ‘1038’ (supplied by Alberto Santini) which is the mother of FL493. Table 1 shows the details of the elm species and cultivars analysed in this study.

**Figure 1.**
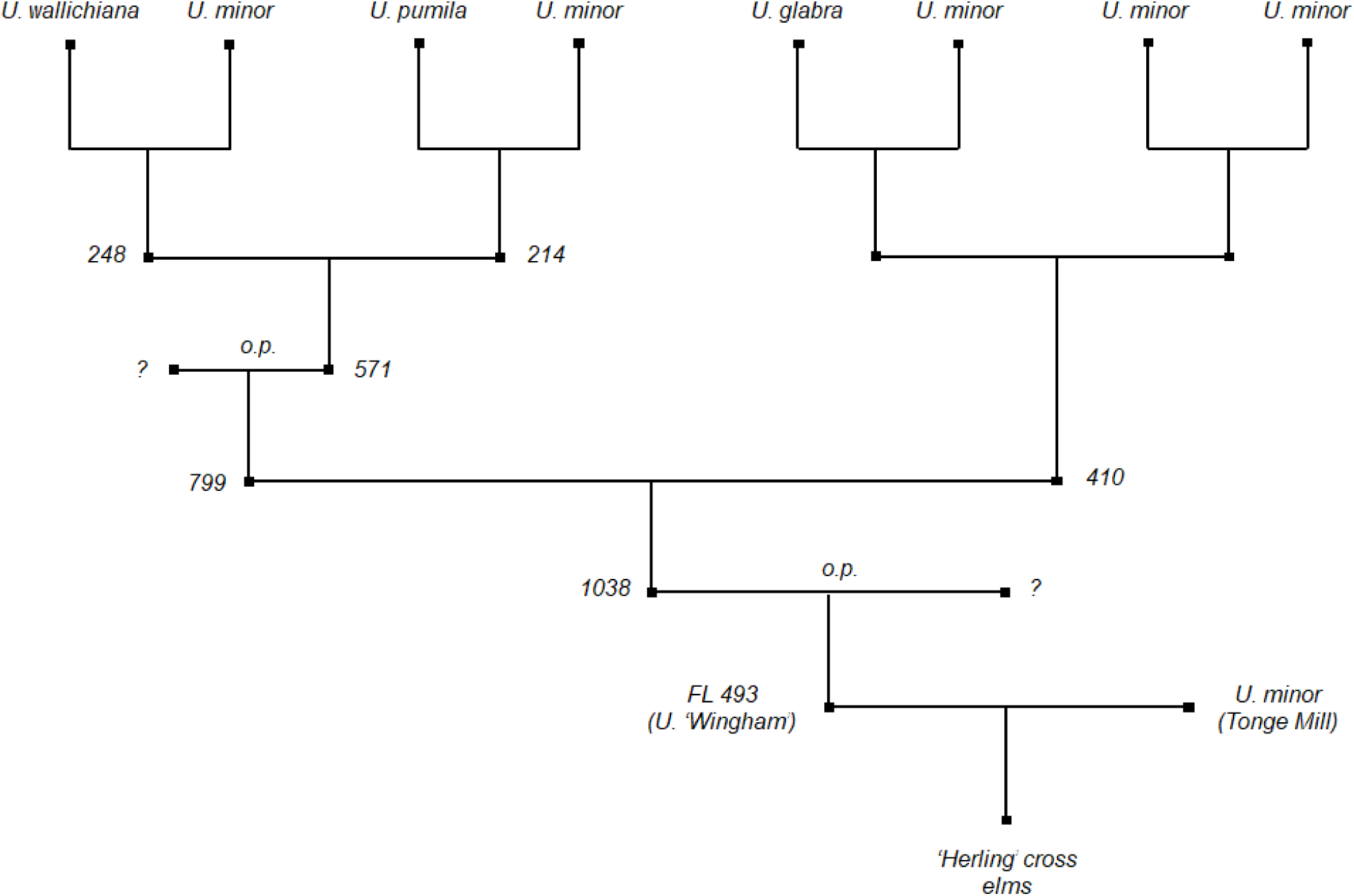
Pedigree diagram showing the breeding history of elm (*Ulmus* spp.) crosses leading to the development of the family analysed in this study. The diagram traces the parentage of FL493 through multiple hybridization events, including contributions from *U. wallichiana*, *U. minor*, *U. pumila*, and *U. glabra*. Breeding programme ID numbers for known individuals in the pedigree are provided, and open pollination (o.p.) events are indicated where applicable. The final generation includes FL493 (‘Wingham’) and *U. minor* (’Tonge Mill’), which generated the WxTM cross elms.

**Table 1.**
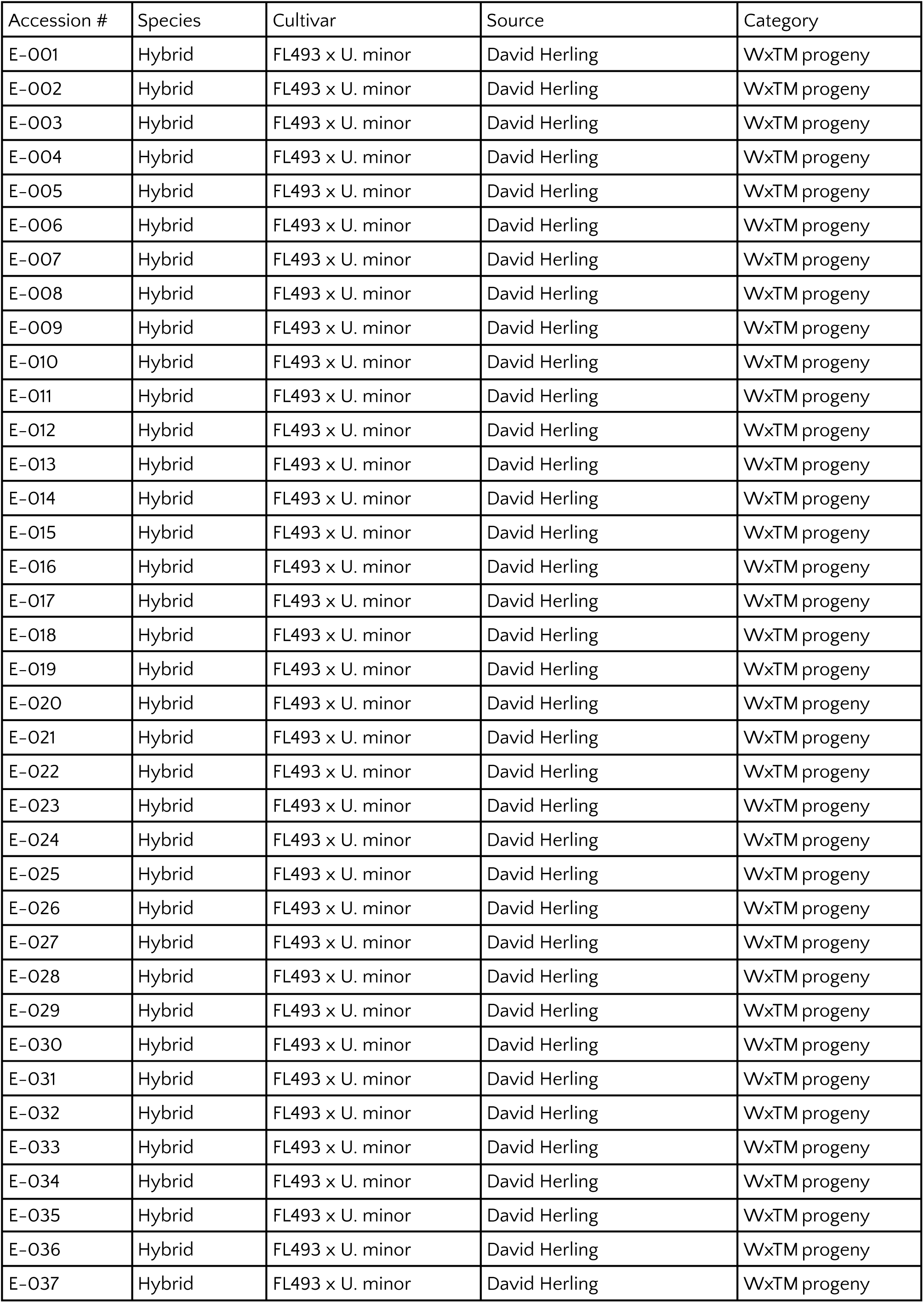
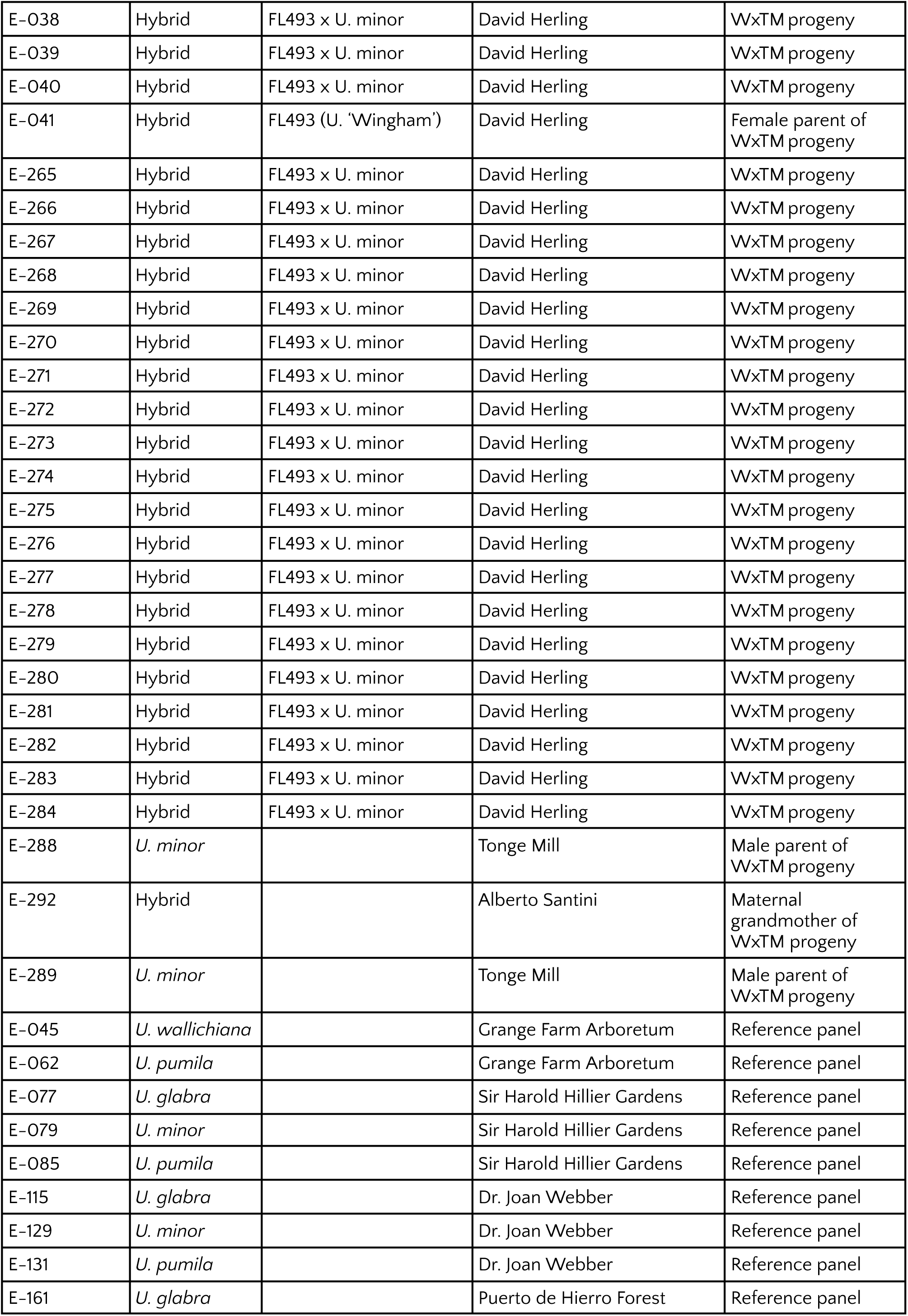
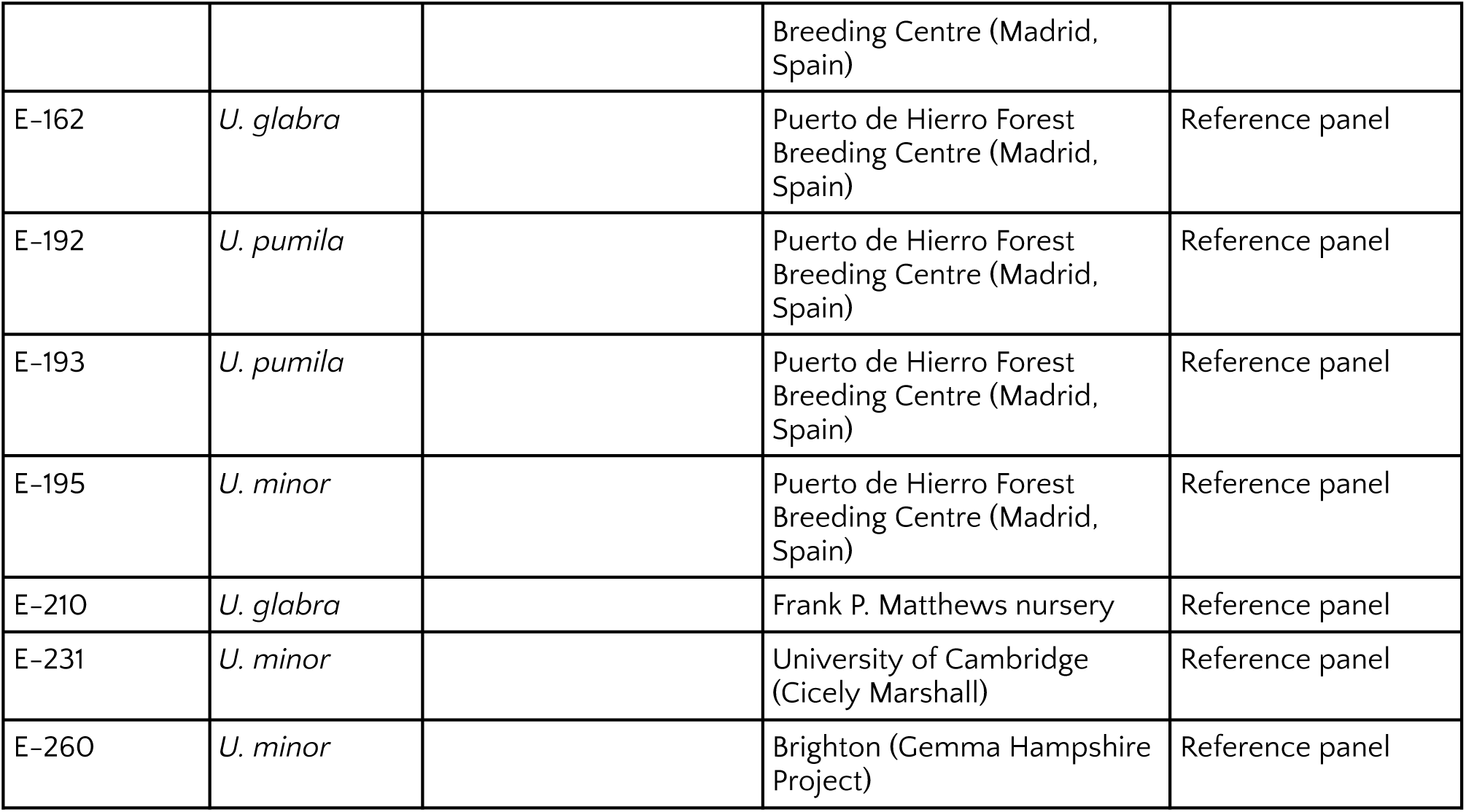
Background information for all individual elms used in this study.

### Inoculation and phenotyping

On 9th June 2019 the WxTM progeny were wound-inoculated with an aggressive strain of *Ophiostoma novo-ulmi* supplied by Clive Brasier and Joan Webber. Elm trees with a known level of susceptibility to DED (Table 3), were also inoculated as controls: four *U. minor* Atinia (syn. *U. procera*) trees in a nearby field, one Ademuz, one Dehesa de Amaniel, one Columella o.p., one FL462, one FL462 x Patriot, Accolade one *U. pumila* "aurescens", three Vada, one FL493 x Patriot, and one “Actons farm”. During the time of the inoculations weather conditions were cool and rainy and thus conducive to disease development. Higher temperatures in late July, at around 6 – 8 weeks after inoculation, at which point DED symptoms usually reach their peak, may have hindered further progression of the disease.

**Table 2.** Alterra disease index for Dutch Elm Disease (DED).

| DED disease score | Symptoms |
| --- | --- |
| 0.0 | No symptoms |
| 0.5 | Start of leaf necrosis |
| 1.0 | Yellowing and/or wilting of leaves, but no dieback of twigs |
| 1.5 | Severe leaf fall and/or dieback of the tips of the twigs or inner twigs of the crown |
| 2.0 | Partly or completely dieback of young top twigs with or without formation of “shepherd’s crooks” |
| 2.5 | Dieback to the base of this year’s shoots and/or massive occurrence of diseased twigs |
| 3.0 | Dieback of 1-year old shoots as well as partly two-year-old shoots (up to halfway) |
| 3.5 | Dieback of 1-year old shoots as well as older shoots |
| 4.0 | Severe dieback (more than half of the tree is dead) |

**Table 3.** Copying matrix (Mu). MOSAIC decouples donor reference panels from internal representations of the original mixing populations. The copying matrix reflects the relationship between the ancestral mixing groups inferred by MOSAIC and each of the 4 donor reference panels. Higher values along the diagonal mean that the admixed haplotypes are reconstructed in terms of haplotypes from the four *Ulmaceae* species.

| Species | Population 1 | Population 2 | Population 3 | Population 4 |
| --- | --- | --- | --- | --- |
| *U. minor* | 0.9 | 0.1/3 | 0.1/3 | 0.1/3 |
| *U. glabra* | 0.1/3 | 0.9 | 0.1/3 | 0.1/3 |
| *U. pumila* | 0.1/3 | 0.1/3 | 0.9 | 0.1/3 |
| *U. wallichiana* | 0.1/3 | 0.1/3 | 0.1/3 | 0.9 |

The control trees were assessed after 4 weeks using the ‘Alterra’ disease index (Buiteveld et al. 2015) (Table 2) and at 4 and 8 weeks using estimated percentage defoliation. The WxTM family were assessed at 4 and 8 weeks using the Alterra disease index and estimated percentage defoliation. The WxTM family were scored after 12 weeks using a simplified binary defoliation score (i.e. <75% or >= 75% defoliation), and after one year using the complete percentage defoliation score (i.e. 0% to 100%).

Leaf samples for DNA extraction were collected on 28th August 2019 for the trees showing the most severe symptoms. Samples from the other trees sequenced for this project were collected on 31st August 2020.

### Whole genome sequencing

Collected leaf materials were preserved in paper or plastic bags with silica gel and stored at room temperature. Genomic DNA was extracted using a modified CTAB method with a Sorbitol pre-wash step (Inglis et al. 2018). The quality and quantity of extracted genomic DNA were checked with 1% agarose gel, Quantus Fluorometer (Promega) and NanoDrop 2000 (Thermo Scientific). Due to the viscosity of some extracts, we used Agencourt AMPure XP (Beckman) beads to purify DNA before library preparation. The library preparation and sequencing was carried out by the Edinburgh Genetics Ltd (Edinburgh, UK). Library preparation was conducted using the egSEQ Enzymatic DNA protocol. Initially, DNA fragments were generated and selected to achieve an approximate insert size of 300-400 base pairs. Subsequently, end repair, A-tailing, and adaptor ligation steps were performed. Then purification and PCR amplification were carried out to enrich the library. Further purification was conducted to ensure high-quality libraries. Prior to sequencing on the MGI DNBseq-T7 platform with paired-end 150 base pair read length, circularisation steps were implemented. Data quality control and filtering were performed using the SOAPnuke software (Chen et al. 2018). Sequencing reads matching 25.0% or more of the adapter sequence (with a maximum of 2 base mismatches allowed) were trimmed. Reads with lengths less than 150 bp were discarded entirely. Reads containing 1.0% or more N content were removed from the dataset. Reads exhibiting homopolymer sequences exceeding 50 bp were eliminated. Reads containing bases with a quality value below 20, constituting 30.0% or more of the entire read, were discarded. The output read quality system was set to Phred+33 to obtain clean reads for subsequent analyses.

### Data pre-processing

To make the read data suitable for input into linkage map construction software and admixture analysis software, a pre-processing bioinformatics pipeline was implemented as follows. FastQC was used to assess the read quality of raw sequencing data, with base quality encoded using Phred+33 quality scores. Raw reads were trimmed using Trimmomatic (Bolger et al. 2014) to remove Illumina-specific adapter sequences and low-quality reads, with leading and trailing bases discarded where they fell below a quality threshold of 5. Reads were also discarded where they fell below an average quality of 15 in a 4-base sliding window, or if the total read length was less than 70 bp. FastQC was then used to confirm that reads had been trimmed correctly, and to establish the quality and number of remaining reads.

Raw reads were aligned to haplotype 1 of the *U. glabra* reference genome (Coleman et al. 2024) (GenBank accession number GCA_964106905.1) using bwa-mem2 (Vasimuddin et al. 2019), and subsequently sorted using samtools. Although the majority of samples were genetically closer to *U. minor* than *U. glabra*, no high-quality reference genome was available for *U. minor*, with *U. glabra* being the closest phylogenetic species (Whittemore et al. 2021) for which a suitable reference genome was available; and moreover, it was necessary to align all samples to the same reference genome in order to carry out downstream analyses. Picard (https://github.com/broadinstitute/picard) was then used to mark PCR duplicates in the resulting alignment files. Samtools (Danecek et al. 2021) was then used to assess coverage depth, % GC content, % of reads mapped, and mapping quality.

BCFtools v? (Danecek et al. 2021) was used to call variants in all samples, with stringent filtering implemented throughout. Where alignments had a Phred-scaled mapping quality score < 30, or base calls had a base quality score < 30, these were discarded prior to variant calling using BCFtools ‘mpileup’. Moreover, alignments were included in variant calling if they were marked as proper pairs by bwa-mem2, and discarded if they were marked as secondary alignments, duplicates, or QC fails. BCFtools ‘call’ was used to generate genomic VCF (gVCF) files containing blocks of homozygous reference calls with a minimum coverage depth of 10 reads, in addition to non-reference genotype calls. Genotype calls within 5 base pairs of indels were discarded, as they are more prone to be genotyping errors. VCF files were then sorted, normalised, and further filtered to remove multiallelic SNPs as well as indels. Finally, files were merged across all samples to produce one VCF file for each chromosome, and filtered to only include variants with Phred-scaled variant quality > 30, 0% missingness across all samples, and >0.01% minor allele frequency. Variant calls were only included if they were supported by a minimum of 5 reads, and a maximum of 40 reads in each individual sample. BCFtools ‘stats’ was then used to generate statistics on the number of variants called, and quality metrics such as transition/transversion (Ts/Tv) ratios. Post-filtering SNP density along each chromosome was plotted using CMplot (LiLin-Yin 2015).

### Construction of linkage map

Lep-MAP3 was used to construct a linkage map based on variant calls from the WxTM cross elms (Rastas 2017), which were generated in the previous step. A pedigree file was also generated to convey the familial relationships between E041 (‘Wingham’, the female parent), E289 (*U. minor* at Tonge Mill, the male parent), E292 (the maternal grandmother), and 60 WxTM progeny. The ‘ParentCall2’ module was used to call parental genotypes, based on genotype information included in the VCF input, with non-informative markers discarded at this step. Markers were then clustered into linkage groups using the ‘SeparateChromosomes2’ module, which determines linkage groups based on ‘logarithm of the odds’ (LOD) scores between pairs of markers. The LOD score limit was set to 5, with marker pairs penalised if they belonged to different contigs (parameter ‘usePhysical=1 0.01’). A LOD score of 3 between two markers is generally considered to be evidence of linkage. Although grandparental data was available, this was not used to aid phasing as the number of markers able to be phased in this way was too small (∼2,000 markers across all 14 linkage groups). The ‘distortionLod=1’ parameter was also used to generate segregation distortion-aware LOD scores (check discussion). The minimum size limit for linkage groups was set to 50 markers.

The ‘OrderMarkers2’ module was subsequently used to carry out marker ordering within each linkage group, once a set of 14 linkage groups had been identified which mapped 1:1 to all 14 chromosomes. The number of merge iterations was set to 10, and all other parameters were given default settings. This computational step first involves using a phasing algorithm to identify maternal and paternal haplotypes. The phased data is then used to identify where recombinations have taken place within each linkage group and calculate genetic distances between markers based on the rates of recombination along each chromosome. The resulting genetic maps were fitted to step-wise functions using the ‘FitStepFunction’ module to remove noise and ensure that maps were strictly monotonically increasing. Genetic distance was plotted against physical distance using RStudio (RStudio Team, 2022, http://www.rstudio.com/), with a maternal and paternal genetic map generated for each chromosome.

### Haplotype phasing

SHAPEIT5 was used to estimate haplotypes for all samples (Hofmeister et al. 2023), with pedigree information used where available to aid phasing. The genetic maps generated in the previous step were also used to improve phasing accuracy. Default phasing parameters were used, and only the ‘phase_common’ module was used.

### Admixture analysis

MOSAIC was used to carry out admixture analysis and estimate local ancestries along each chromosome (Salter-Townshend and Myers 2019). Phased haplotypes generated in the previous step were used to create reference panels, with one reference panel for each of the four species contributing to the genetic admixture in the WxTM cross elms. Five samples were included for each ‘parental’ species (*U. minor, U. pumila, U. glabra, U. wallichiana*) in 4 donor panels, with the exception of *U. wallichiana* for which only one sample was available. The genetic maps generated in the previous steps were used to model recombination rates along each chromosome. A four-way admixture model was implemented, and the ‘singlePI’ flag was used. Since the sources of genetic admixture in the WxTM cross elms are known, the value of ‘Mu’ was updated to a diagonally dominant copying matrix (Table 3), to ensure that MOSAIC’s internal representation of ancestral populations mapped 1:1 to the donor panels. The admixture analysis was run on all 14 chromosomes simultaneously to infer the parameters theta, rho, alpha, and Pi, with 20 rounds of EM inference carried out; only the parameter Mu was not updated during EM inference.

We qualitatively inspected whether resistant and susceptible individuals differed in the proportion of the genome ascribed to a particular parental species at specific locations in the chromosome. We calculated the mean local ancestry ascribed to each of the four parental species for susceptible and resistant individuals separately. Proportions by chromosome were then plotted using R. Chromosomal regions showing clear differences between the two groups are promising candidates for containing genes conferring resistance to DED.

Finally, we explored whether variation in the proportion of the genome ascribed to any of the parental species had an effect on the susceptibility to DED among individuals in the WxTM cross progeny. Susceptibility here was defined by the percentage of defoliation after four different periods (i.e. four weeks, eight weeks, 12 weeks, and one year) following inoculation (Table 4). If DED-resistance were very highly polygenic, we would expect individuals with a higher proportion of the genome inherited from Asiatic species to have undergone less defoliation than those individuals with higher proportion of *U. minor* genetic material. To test this, we regressed the percentage of the genome ascribed to one of the parental species (estimated with Admixture analysis) on the percentage of defoliation at the four different time points. For year one and weeks four and eight, we used a beta regression since we were dealing with proportion data. For week 12 we used a logistic regression since defoliation was originally registered as a binary variable (susceptible >= 70% defoliation, resistant < 70% defoliation). Significance tests were performed with likelihood ratio tests using the lmtest v0.9-40 package in R (Zeileis and Hothorn 2002).

**Table 4:**
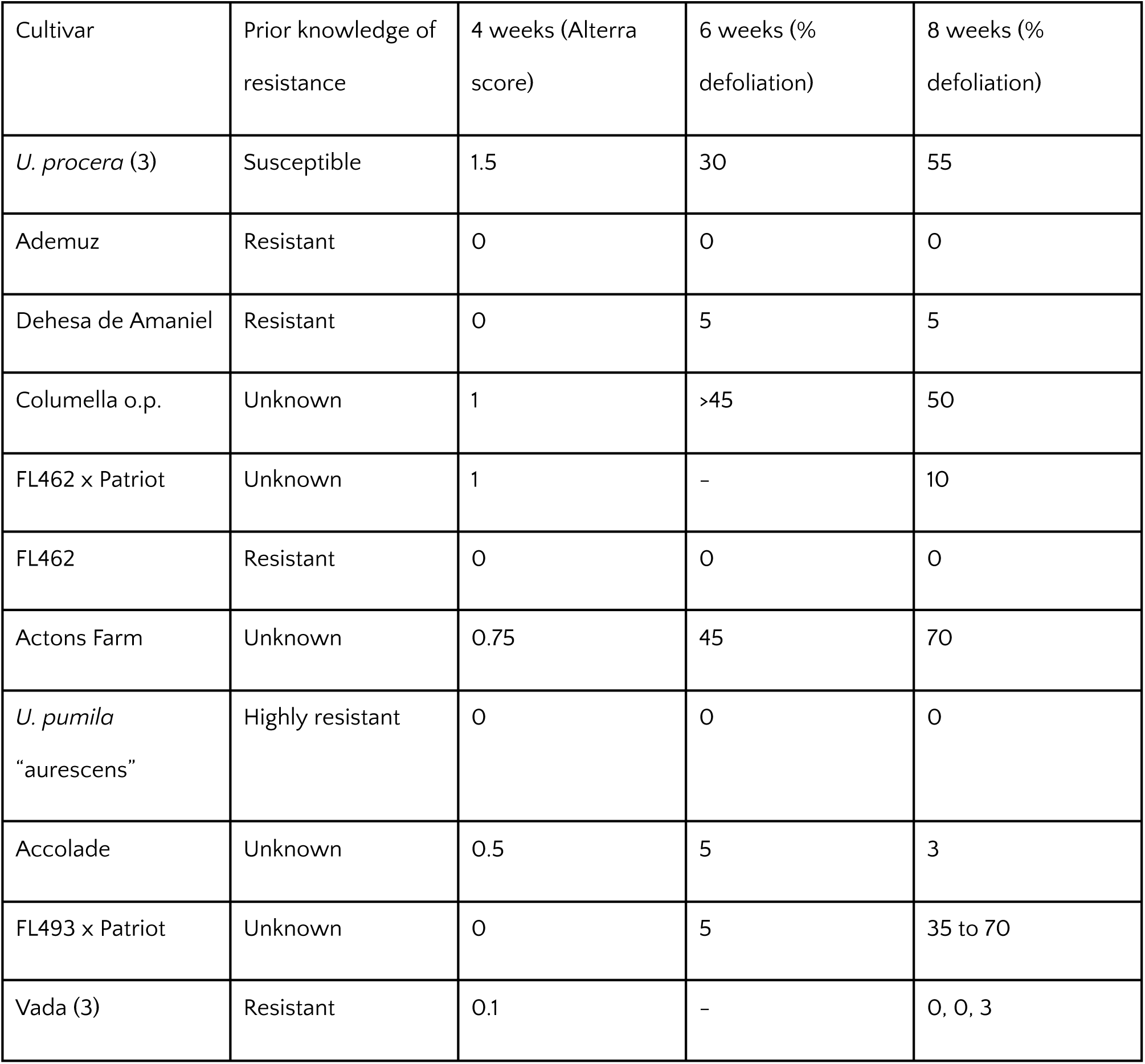
Phenotype scores in control trees. Numbers in parenthesis refer to the sample size.

## Results

### Phenotyping

Damage to the control trees ranged from zero to heavy damage and showed a pattern expected from the known levels of DED resistance (Table 4).

On 7th July 2019, four weeks post-inoculation, in WxTM progeny, 12 plants were symptomless: (4.3, 4.7, 4.16, 5.13, 5.22, 5.27, 5.4, 6.13, 6.2, 6.21, 6.23, 6.9). On 4th August, 8 weeks post-inoculation, 6.21 remained asymptomatic and 6.13 had foliar damage of 1%. At the other extreme, 6.7 had 95% foliar loss or damage, and was on an Alterra disease index of 4 equating to "severe dieback; more than half the tree is dead". Among the progeny, 19 plants showed some signs of regrowth, five plants had foliar damage of 10% or less, two had regrowth (or continued extension growth) despite the death of lower leaves and twigs. Full defoliation results for 4, 8, and 12 weeks, and one year post inoculation, are shown in Table 5.

**Table 5.** Defoliation rates and Alterra scores of sequenced WxTM progeny at multiple time points following inoculation procedure on 28th August 2019.

| Row and tree number | DNA Sample identifier | Alterra score at 4 weeks | Defoliation at 4 weeks | Alterra score at 8 weeks | Defoliation at 8 weeks | Defoliation at 12 weeks | Defoliation at one year |
| --- | --- | --- | --- | --- | --- | --- | --- |
| 5.1 | E-001 | - | - | - | - | <70% | 1% |
| 5.2 | E-002 | 0.5 | 5% | 1.5 | 45% | <70% | 0% |
| 5.3 | E-003 | 0.5 | 5% | 1.5 | 35% | <70% | 5% |
| 5.6 | E-004 | 1.5 | 15% | 2 | 65% | <70% | 0% |
| 5.8 | E-005 | 1 | 5% | 1.5 | 30% | <70% | 15% |
| 5.12 | E-006 | - | - | - | - | <70% | 0% |
| 5.13 | E-007 | 0 | 0% | 2 | 50% | <70% | 45% |
| 5.14 | E-008 | 0.5 | 5% | 2 | 50% | <70% | 40% |
| 5.16 | E-009 | 2.5 | 50% | 2.5 | 55% | <70% | 5% |
| 5.17 | E-010 | 1.5 | 10% | 2 | 65% | <70% | 15% |
| 5.18 | E-011 | 2 | 20% | 2 | 50% | <70% | 40% |
| 5.19 | E-012 | 1.5 | 20% | 2 | 55% | <70% | 35% |
| 5.2 | E-013 | 0.5 | 5% | 1.5 | 40% | <70% | 75% |
| 5.21 | E-014 | 0.25 | 2% | 1.5 | 35% | <70% | 2% |
| 5.22 | E-015 | 0 | 0% | 0.25 | 2% | <70% | 0% |
| 5.23 | E-016 | 2.5 | 35% | 3.5 | 90% | <70% | 2% |
| 5.24 | E-017 | 1.5 | 15% | 1 | 10% | <70% | 2% |
| 5.25 | E-018 | 1.5 | 15% | 2 | 30% | <70% | 2% |
| 5.26 | E-019 | - | - | - | - | <70% | 0% |
| 5.27 | E-020 | 0.25 | 0% | 1.5 | 20% | <70% | 0% |
| 6.1 | E-021 | - | - | - | - | <70% | 30% |
| 6.2 | E-022 | 1.75 | 15% | 2 | 40% | <70% | 30% |
| 6.3 | E-023 | 1.25 | 5% | 2 | 50% | <70% | 10% |
| 6.4 | E-024 | 2.5 | 50% | 2.5 | 65% | <70% | 75% |
| 6.5 | E-025 | 1 | 20% | 2.5 | 65% | <70% | 20% |
| 6.6 | E-026 | - | - | - | - | <70% | 95% |
| 6.8 | E-027 | 1.25 | 20% | 2.5 | 70% | <70% | 5% |
| 6.9 | E-028 | 0.25 | 0% | 1.5 | 35% | <70% | 45% |
| 5.1 | E-029 | 0.75 | 35% | 1.5 | 55% | <70% | 95% |
| 6.11 | E-030 | 0.5 | 5% | 1.5 | 55% | <70% | 0% |
| 6.12 | E-031 | 0.75 | 10% | 2.5 | 55% | <70% | 25% |
| 6.13 | E-032 | 0 | 0% | 0.5 | 1% | <70% | 2% |
| 6.14 | E-033 | 1.25 | 15% | 2.5 | 70% | <70% | 5% |
| 6.15 | E-034 | - | - | - | - | <70% | 0% |
| 6.18 | E-035 | - | - | - | - | <70% | 0% |
| 6.2 | E-036 | 0.25 | 0% | 1.5 | 20% | <70% | 40% |
| 6.21 | E-037 | 0 | 0% | 0 | 0% | <70% | 0% |
| 6.23 | E-038 | 0 | 0% | 0.25 | 2% | <70% | 30% |
| 6.24 | E-039 | 0.5 | 5% | 2.5 | 75% | <70% | 0% |
| 6.25 | E-040 | - | - | - | - | <70% | 0? |
| 4.3 | E-265 | 0.75 | 0% | 2.5 | 75% | >70% | 10% |
| 4.7 | E-266 | 1 | 0% | 2 | 70% | >70% | 45% |
| 4.12 | E-267 | 1.5 | 5% | 2.5 | 80% | >70% | 20% |
| 4.16 | E-268 | 0.25 | 0% | 2 | 45% | >70% | 15% |
| 4.18 | E-269 | 0.5 | - | 2.5 | 70% | >70% | 40% |
| 4.2 | E-270 | 2 | 20% | 3.5 | 90% | >70% | 95% |
| 4.21 | E-271 | 2 | 20% | 3.5 | 90% | >70% | 95% |
| 5.4 | E-272 | 0.25 | 0% | 1.5 | 35% | >70% | 75% |
| 5.5 | E-273 | 1.5 | 15% | 2.5 | 70% | >70% | 65% |
| 5.7 | E-274 | 1.5 | 15% | 2 | 75% | >70% | 10% |
| 5.9 | E-275 | 2 | 30% | 2.5 | 80% | >70% | 10% |
| 5.1 | E-276 | 2 | 30% | 2.5 | 80% | >70% | 10% |
| 5.11 | E-277 | 2 | 25% | 2.5 | 70% | >70% | 2% |
| 5.15 | E-278 | 2.5 | 30% | 3 | 90% | >70% | 75% |
| 5.22 | E-279 | 2.5 | 30% | 2.5 | 70% | >70% | 50% |
| 6.7 | E-280 | 2.5 | 50% | 4 | 95% | >70% | dead |
| 6.16 | E-281 | 1.75 | 30% | 3 | 80% | >70% | 75% |
| 6.17 | E-282 | 1.5 | 35% | 3 | 75% | >70% | 75% |
| 6.19 | E-283 | 1.5 | 35% | 3 | 80% | >70% | 75% |
| 6.26 | E-284 | - | - | - | - | >70% | Dead? |

### Sequence data and quality control

Prior to trimming, the number of raw reads ranged from 126.0 M – 200.6 M across all samples. Following adapter trimming and the removal of low-quality/unpaired reads, the number of reads ranged from 106.0 M – 180.6 M. % GC content ranged from 31.0% – 38.0%. Across all samples, 98.5% of trimmed reads mapped to the *U. glabra* reference genome, with 80.6% of these reads mapping as proper pairs (i.e. both ends of a read mapped within a reasonable distance of one another). Duplicate reads ranged from 0.4% - 10.4%, with the majority of samples reporting duplicate rates of < 3%. After SNP calling and stringent filtering, 5,497,580 high quality SNPs remained for downstream analysis (Table 6). Their density along each chromosome is provided in Figure 3.

**Figure 2.**
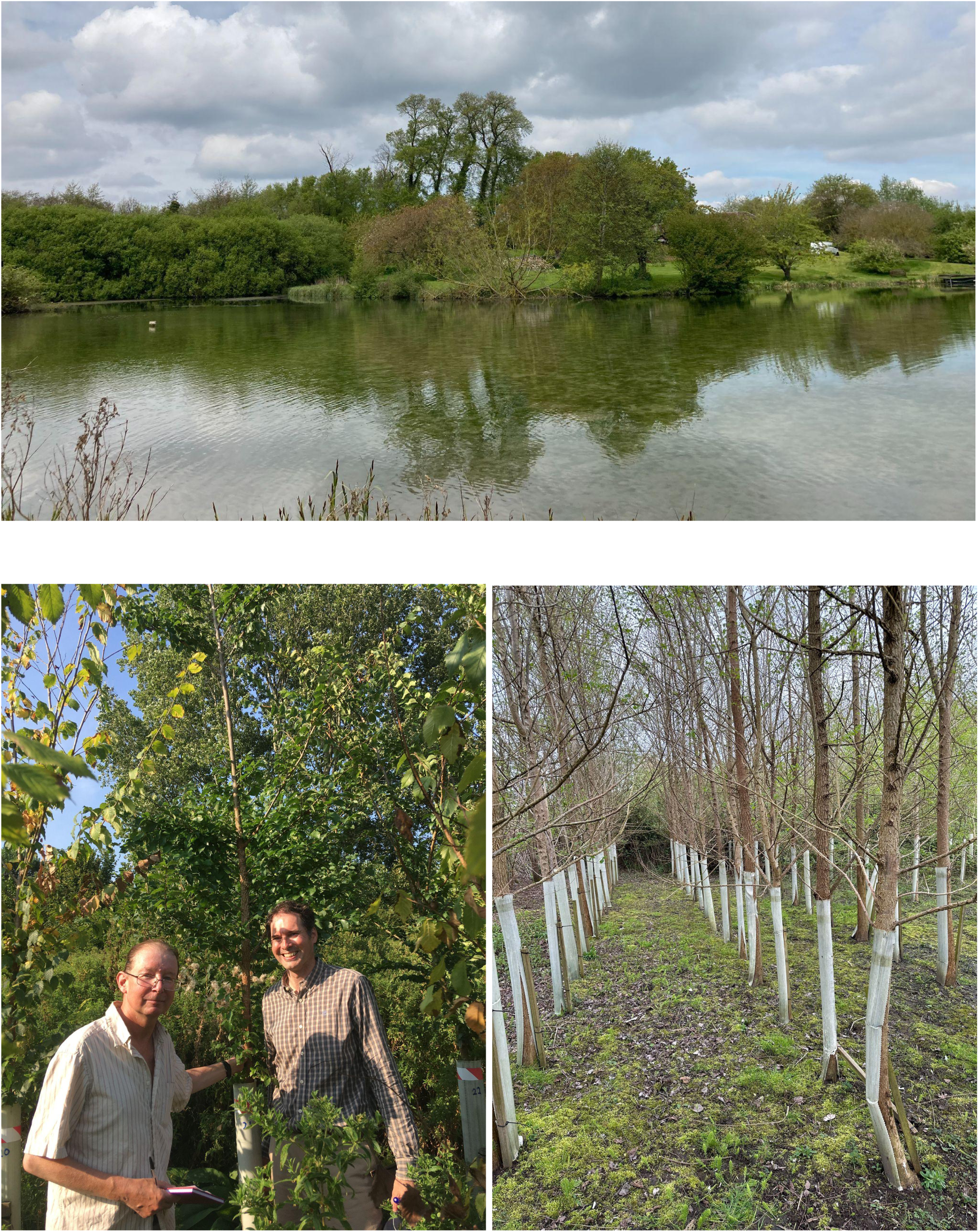
Top: The Tonge Mill elms in 2024 (stand in centre; Photo: Richard Buggs). Left: David Herling and Fergus Poncia by tree 6.21 in 2019 (Photo: Richard Buggs). Right: the WxTM progeny in 2024 (Photo: Fergus Poncia).

**Figure 3.**
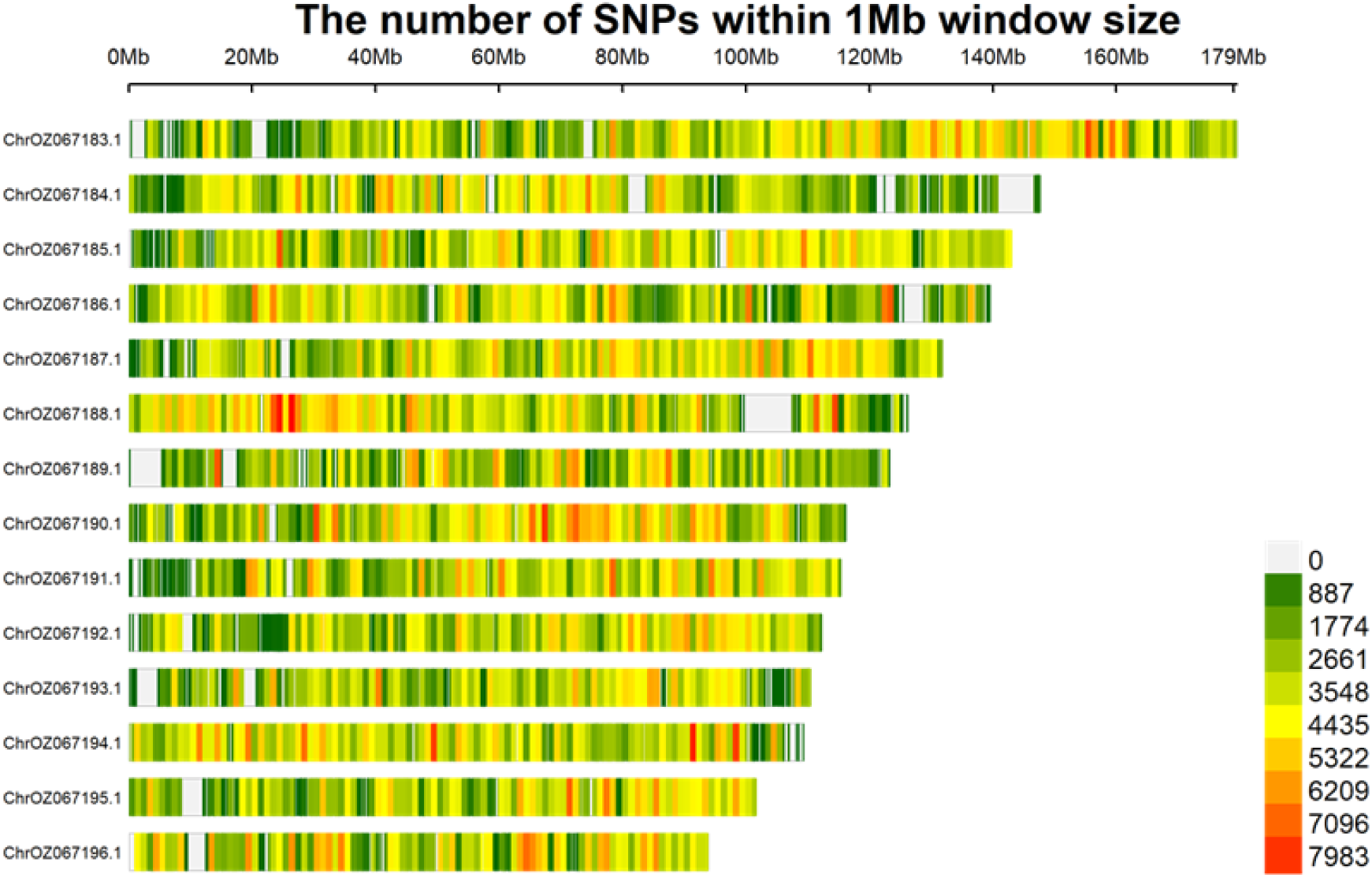
Plot of SNP density along each chromosome. Generated using CMplot.

**Table 6.** Number of SNPs remaining post-filtering, with reported transition/transversion (Ts/Tv) ratios for each chromosome.

| Chromosome | No. of SNPs | Ts/Tv ratio |
| --- | --- | --- |
| 1 | 550,038 | 2.41 |
| 2 | 402,259 | 2.38 |
| 3 | 469,301 | 2.42 |
| 4 | 414,778 | 2.40 |
| 5 | 440,350 | 2.42 |
| 6 | 443,381 | 2.36 |
| 7 | 328,927 | 2.36 |
| 8 | 414,953 | 2.42 |
| 9 | 356,277 | 2.37 |
| 10 | 350,587 | 2.37 |
| 11 | 314,566 | 2.38 |
| 12 | 383,090 | 2.39 |
| 13 | 316,365 | 2.37 |
| 14 | 312,708 | 2.38 |
| **Total/average:** | 5,497,580 | 2.39 |

### Linkage map construction

The vast majority of software tools designed to conduct admixture analysis require as input genetic maps (also sometimes referred to as ‘linkage maps’), which model the rates of recombination along each chromosome. Since to the best of our knowledge no linkage maps have been developed for any *Ulmaeceae* species, we exploited the family structure of the WxTM cross elms to develop the first *de novo* genetic maps for *Ulmaeceae*.

**Table 7.** Physical lengths and genetic distances for each chromosome, alongside the estimated number of recombinations in the WxTM progeny.

| Chromosome | # markers in linkage group | # recombinations | Maximum genetic distance (male, cM) | Maximum genetic distance (female, cM) | Maximum genetic distance (sex-averaged, cM) | Physical length (Mb) |
| --- | --- | --- | --- | --- | --- | --- |
| 1 | 30,660 | 117 | 121.2 | 73.3 | 97.5 | 179.5 |
| 2 | 22,510 | 103 | 86.7 | 68.3 | 77.5 | 147.7 |
| 3 | 21,838 | 116 | 110.0 | 68.3 | 89.2 | 143.0 |
| 4 | 20,856 | 88 | 76.7 | 66.7 | 71.7 | 139.5 |
| 5 | 20,499 | 113 | 93.3 | 73.3 | 83.3 | 131.8 |
| 6 | 15,827 | 105 | 81.7 | 50.0 | 65.8 | 126.2 |
| 7 | 14,384 | 94 | 86.7 | 68.3 | 77.5 | 123.2 |
| 8 | 14,138 | 102 | 85.0 | 50.0 | 67.5 | 116.1 |
| 9 | 14,070 | 87 | 81.7 | 63.3 | 72.5 | 115.4 |
| 10 | 13,597 | 95 | 78.3 | 65.8 | 65.8 | 112.3 |
| 11 | 12,905 | 105 | 76.7 | 60.0 | 68.3 | 110.4 |
| 12 | 11,752 | 112 | 90.0 | 68.3 | 79.2 | 109.3 |
| 13 | 10,964 | 95 | 65.0 | 66.7 | 65.8 | 101.7 |
| 14 | 10,822 | 96 | 60.0 | 50.0 | 55.0 | 93.9 |
| **Total** | 234,822 | 1428 | 1193.0 | 892.3 | 1036.6 | 1750.0 |
| **Average** | 16,773 | 102 | 85.2 | 63.7 | 74.0 | 125.0 |

We estimated the number of recombination events in the WxTM cross progeny (excluding E292, the mother of ‘Wingham’) to range from 88 to 117, depending on the chromosome (Table 6). Genetic distances were calculated separately for the maternal and paternal parents, i.e. E041 (‘Wingham’) and E292, *U. minor* at Tonge Mill respectively. Note that the physical lengths are based off of the reference genome assembly for *U. glabra* and are therefore not completely accurate.

We calculated the genetic distance, measured in centiMorgans, between each pair of markers, with 1 centiMorgan reflecting a 1% likelihood of two markers becoming separated due to recombination. We found that the total genetic length of the male maps (1193.0 cM) is significantly higher than that of the female maps (892.3 cM), with a discrepancy of 300.7 cM reflecting higher rates of recombination in males (Figures 4a,b). We note that certain regions demonstrate the absence of any recorded recombinations (e.g. the region at 100 - 120 Mb in chromosome 6), which may simply be due to the small size of the WxTM family (n = 62, excluding E292).

**Figure 4a.**
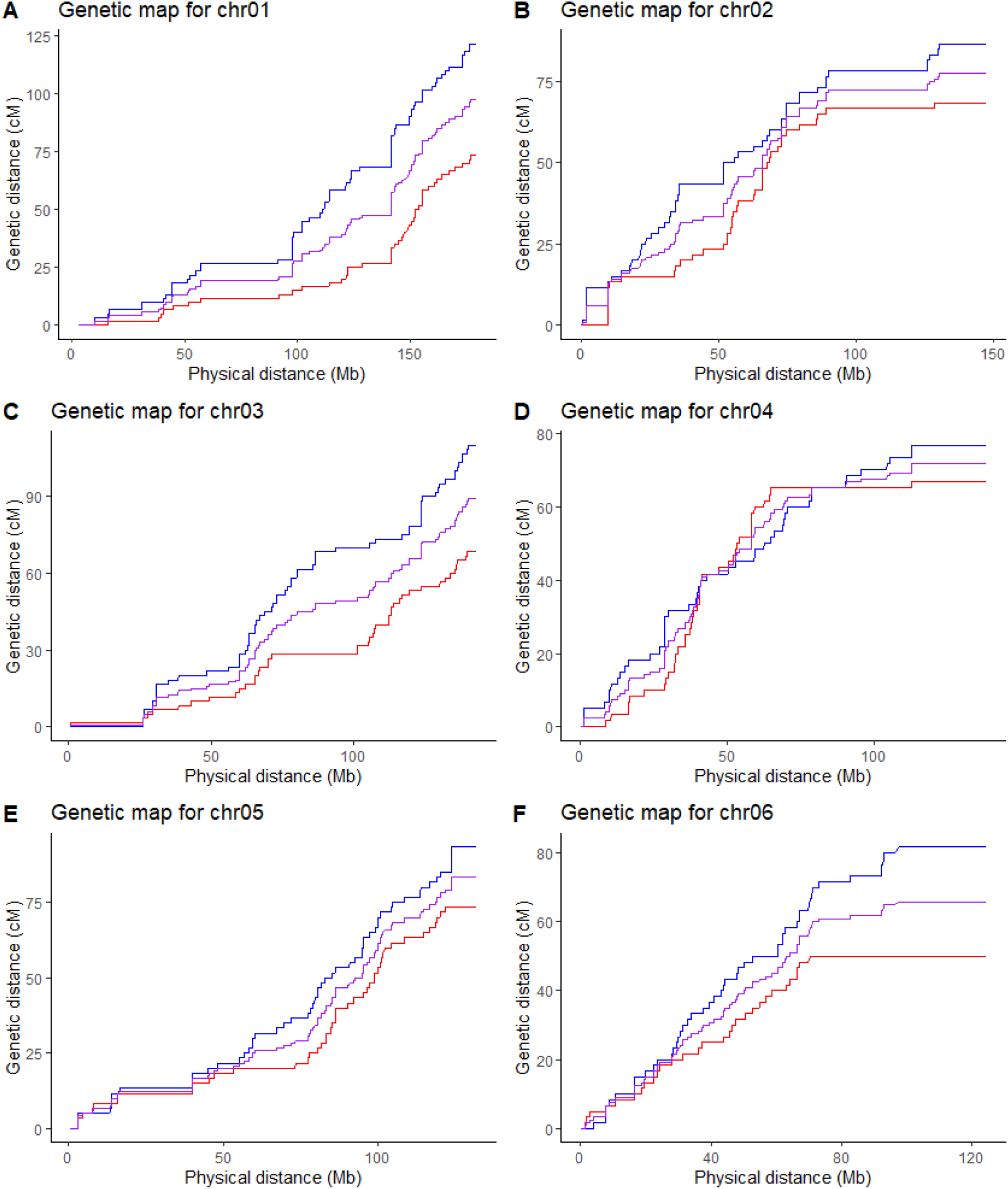
Genetic maps for chromosomes 1-6. Physical distance (Mb) is plotted against cumulative genetic distance (cM). The male map is plotted in blue; the female map is plotted in red; the sex-averaged map is plotted in purple. Steeper gradients reflect higher rates of recombination.

**Figure 4b.**
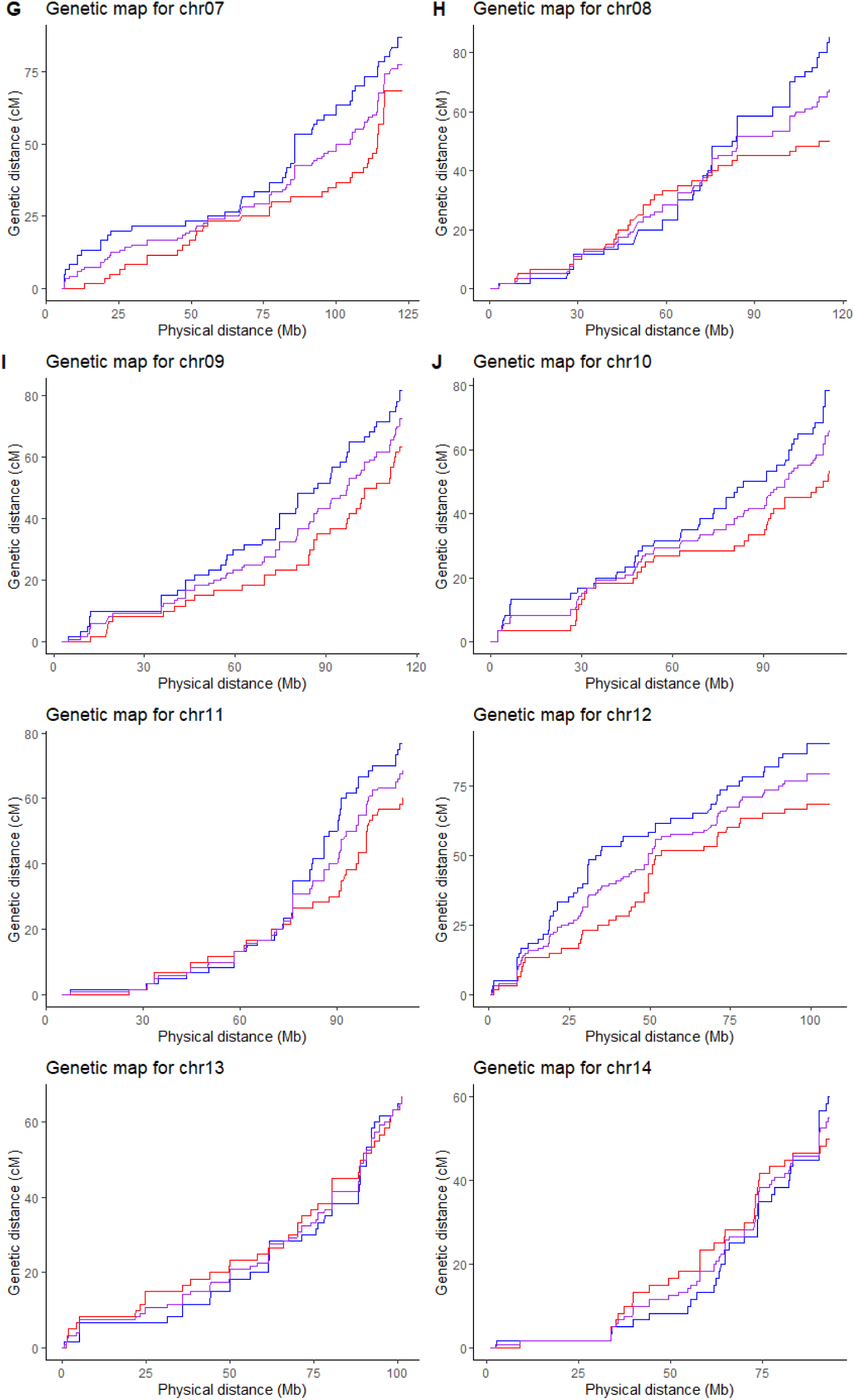
Genetic maps for chromosomes 7-14. Physical distance (Mb) is plotted against cumulative genetic distance (cM). The male map is plotted in blue; the female map is plotted in red; the sex-averaged map is plotted in purple. Steeper gradients reflect higher rates of recombination.

### Admixture analysis

Local ancestry along each chromosome was estimated for the WxTM progeny and their parents using MOSAIC. We define local ancestry as the number of alleles (0, 1, or 2) from each of the 4 parental species contributing to the genetic admixture. Note that although the values for local ancestry should theoretically be confined to 0, 1, or 2, the y-values on the karyograms can vary between these values due to uncertainty in the inferred probabilities.

Figure 5 shows karyograms for the parents of the WxTM cross and one grandparent (1038), and Figures S1a-k show karyograms for every individual. The admixture analysis identified *U. minor* at Tonge Mill (Figure 5a) as *U. minor*, but with numerous small regions containing ancestry from *U. glabra* and a very small amount of *U. wallichiana*. These results differ among the two Tonge Mill elms that we sequenced, however, with Individual 62 showing less evidence of *U. wallichiana* (Figure S1k). While it is possible that these elms contain some introgression from other species, this signal could also be a misassignment of local ancestry by MOSAIC due to the limited size of the reference panels for each parental population. For *U. minor* and *U. glabra* the relatedness between these two species (Whittemore et al. 2021) may have made inaccurate assignments more likely.

**Figure 5.**
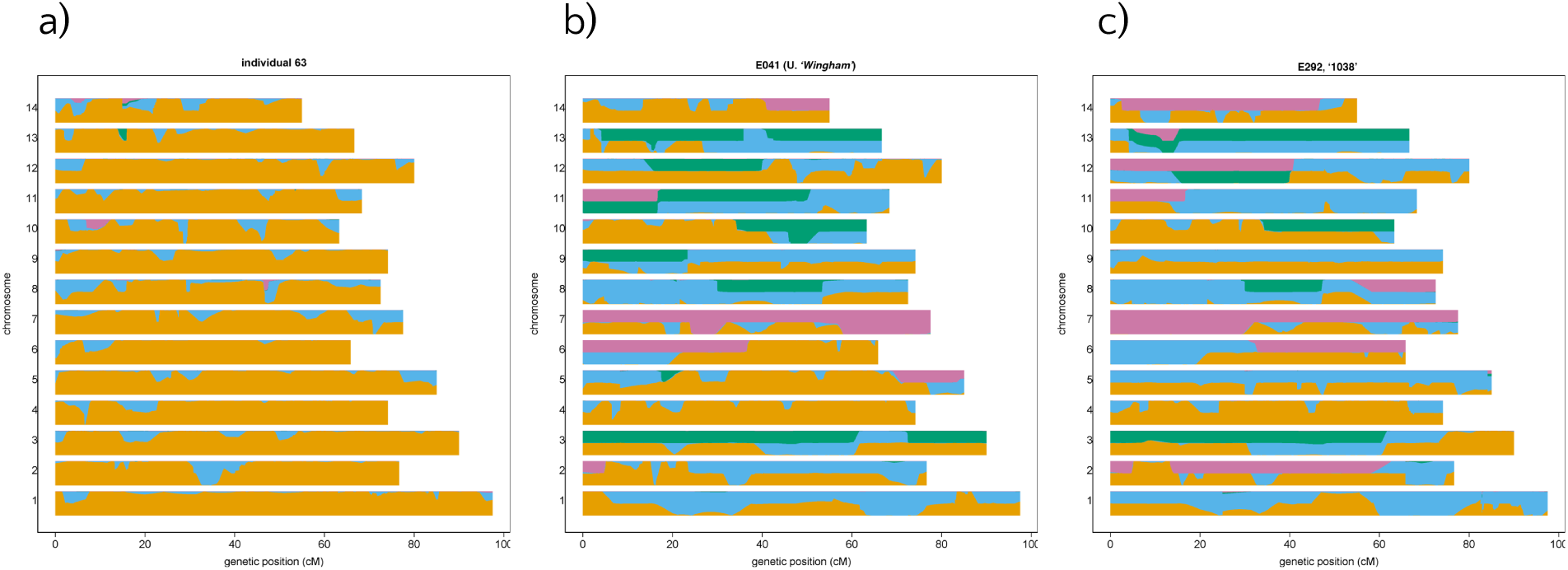
Karyogram for (a) Tonge Mill (individual 63), (b) FL483 ‘Wingham’ (individual 41), and (c) 1038 (Individual 64, E292) the maternal parent of FL483. Y-axis values for each chromosome indicate the estimated number of alleles from each ancestral species; genetic distance (cM) is plotted along the x-axis. Local ancestry assignments are indicated by colour; orange = *U. minor*, blue = *U. glabra*, green = *U. pumila*, pink = *U. wallichiana*.

As expected, ‘Wingham’ (Figure 1b) and 1038 (Figure 1c) show a complex mixture of material from the parental species shown in Figure 1, albeit mainly composed of *U. minor* and *U. glabra*. The differences between 1038 and ‘Wingham’ shed some light on the open pollination that gave rise to ‘Wingham’. The fact that ‘Wingham’ appears to have admixture from *U. pumila* and *U. wallichiana* in some chromosomal regions where 1039 does not, suggests that its father was also a complex hybrid. An alternative hypothesis is that in these regions we have insufficiently sampled the variation present in each species, and cannot fully distinguish them. This could cause the analysis to misassign some regions.

Global ancestry of each sample, i.e. the genome-wide proportions of ancestry from each of the 4 parental species, was also estimated by summing over local ancestry estimates (Figure 6).

**Figure 6.**
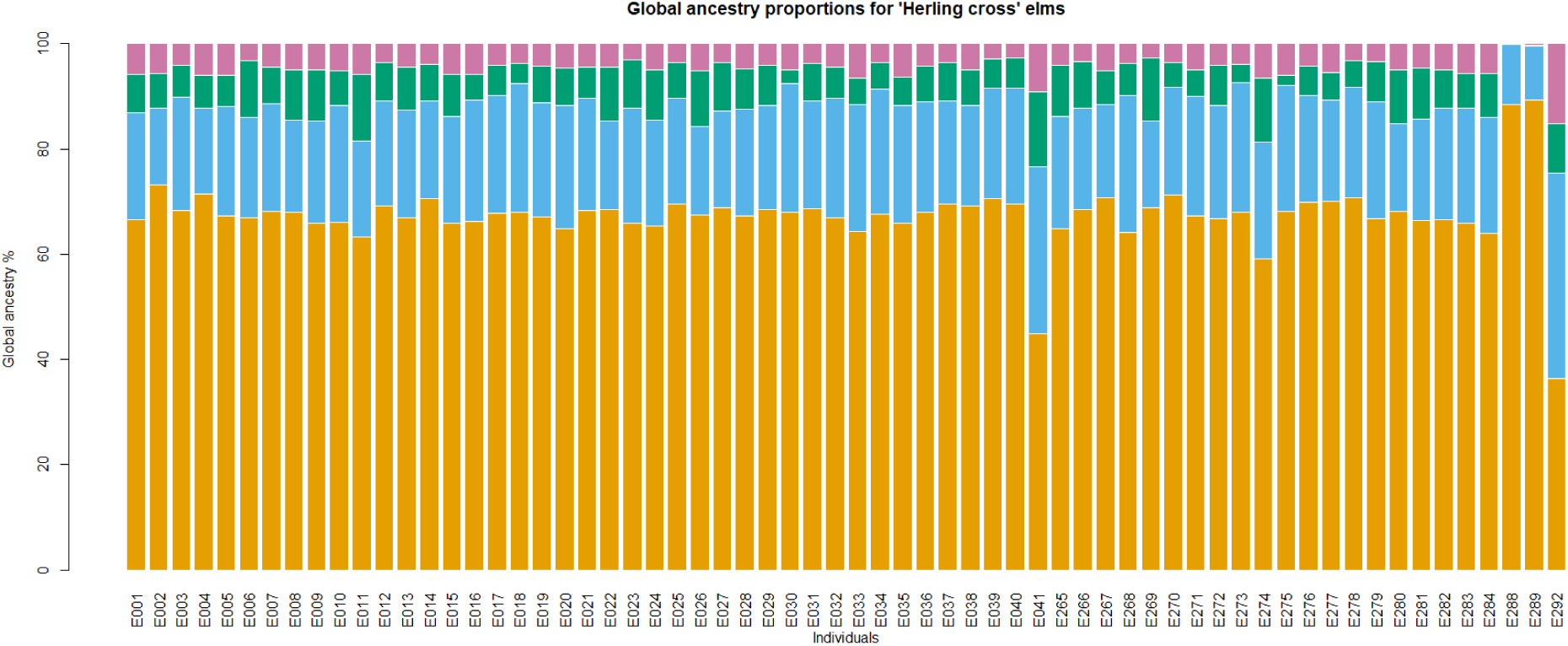
Estimated global ancestry for the WxTM cross elms, with sample labels (x-axis) and genome-wide proportions from each parental species (y-axis). Ancestry assignments are indicated by colour; orange = *U. minor*, blue = *U. glabra*, green = *U. pumila*, pink = *U. wallichiana*. The final three columns show two elms from Tonge Mill and also ‘Wingham’ (FL493).

The average compositions of the chromosomes in the WxTM progeny as percentages of ancestral species are shown in Figure 7.

**Figure 7.**
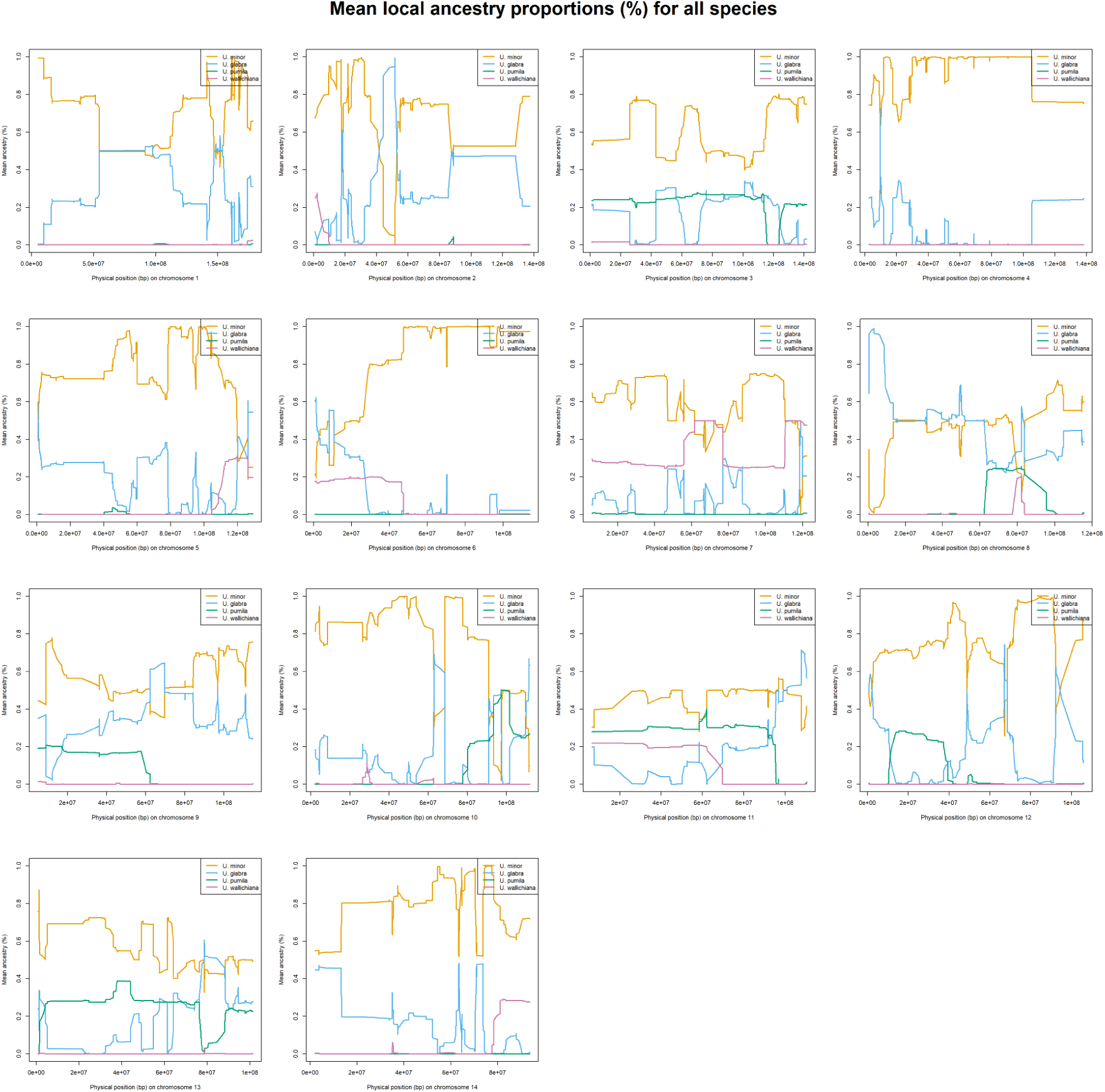
Mean local ancestry percentages for the WxTM progeny from all four ancestral species, for all 14 chromosomes. Ancestry assignments are indicated by colour; orange = *U. minor*, blue = *U. glabra*, green = *U. pumila*, pink = *U. wallichiana*.

Within the admixed elms, we were able to identify haplotypes from all four parental species owing to extensive admixture linkage disequilibrium. The genetic composition of the WxTM progeny, on average, comprised 67.58% ancestry from *U. minor*, 20.67% ancestry from *U. glabra*, 7.16% ancestry from *U. pumila*, and 4.59% ancestry from *U. wallichiana* (Figures S1a-k). We identified haplotypes inherited from *U. pumila* in chromosomes 3, 8, 9, 10, 11, 12, and 13; and haplotypes inherited from *U. wallichiana* in chromosomes 2, 5, 6, 7, 11, and 14.

Comparisons between the mean ancestry proportions along the genome in resistant and susceptible individuals based on a threshold of ≤25% defoliation, 1 year post-inoculation are shown in Figures 8-11 (similar comparisons for other phenotypic assessments are shown in Supplementary Figures S2 to S25). In many regions of the genome there is little difference between the resistant and susceptible individuals. However, under the assumption that resistance is most likely to be conferred by *U. wallichiana* or *U. pumila*, and least likely to be conferred by *U. minor*, there are several regions that are interesting in terms of the differences they show between susceptible and resistant WxTM progeny. There is a region in chromosome 7 where resistant individuals had a consistently higher proportion of the genome ascribed to *U. wallichiana* (Figure 11) than susceptible individuals and a lower proportion of *U. minor* (Figure 8). In chromosome 14, there is a region with higher *U. glabra* (Figure 9) and lower *U. minor* (Figure 8) in resistant individuals. In chromosome 12 there is a region with higher *U. pumila* (Figure 10) and lower *U. minor* (Figure 8) in resistant individuals. In chromosome 3 there is a region with lower *U. glabra* (Figure 9) and higher *U. pumila* (Figure 10) in resistant individuals. In chromosome 11 there is a region where *U. wallichiana* (Figure 11) is higher and *U. pumila* (Figure 10) is lower in resistant individuals. All these regions deserve further investigation for involvement in resistance.

**Figure 8.**
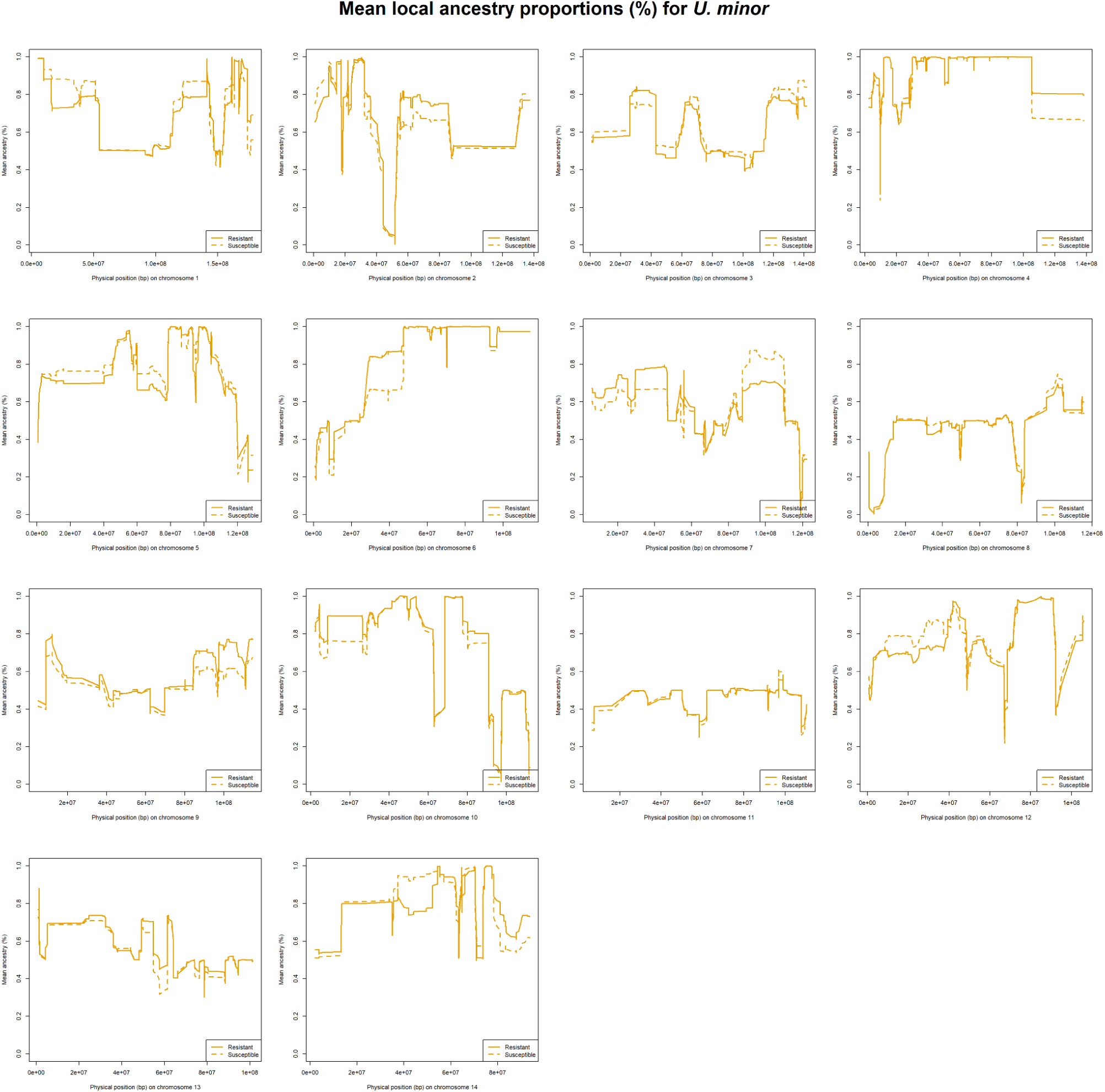
Mean local ancestry proportions (%) for *U. minor*, comparing resistant WxTM progeny (solid line) with susceptible WxTM progeny (dashed line), based on a threshold of ≤25% defoliation, 1 year post-inoculation.

**Figure 9.**
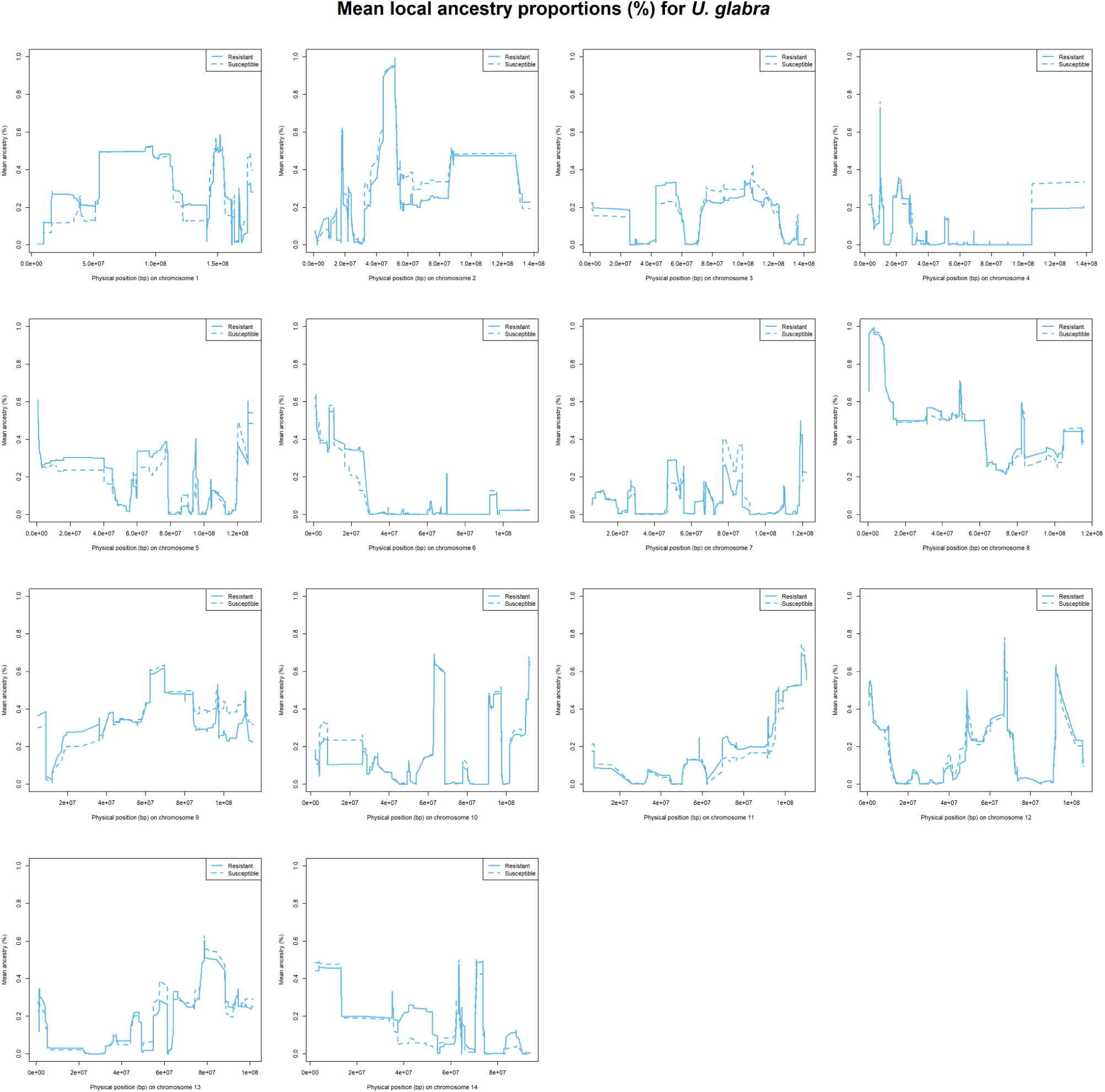
Mean local ancestry proportions (%) for *U. glabra*, comparing resistant WxTM progeny (solid line) with susceptible WxTM progeny (dashed line), based on a threshold of ≤25% defoliation, 1 year post-inoculation.

**Figure 10.**
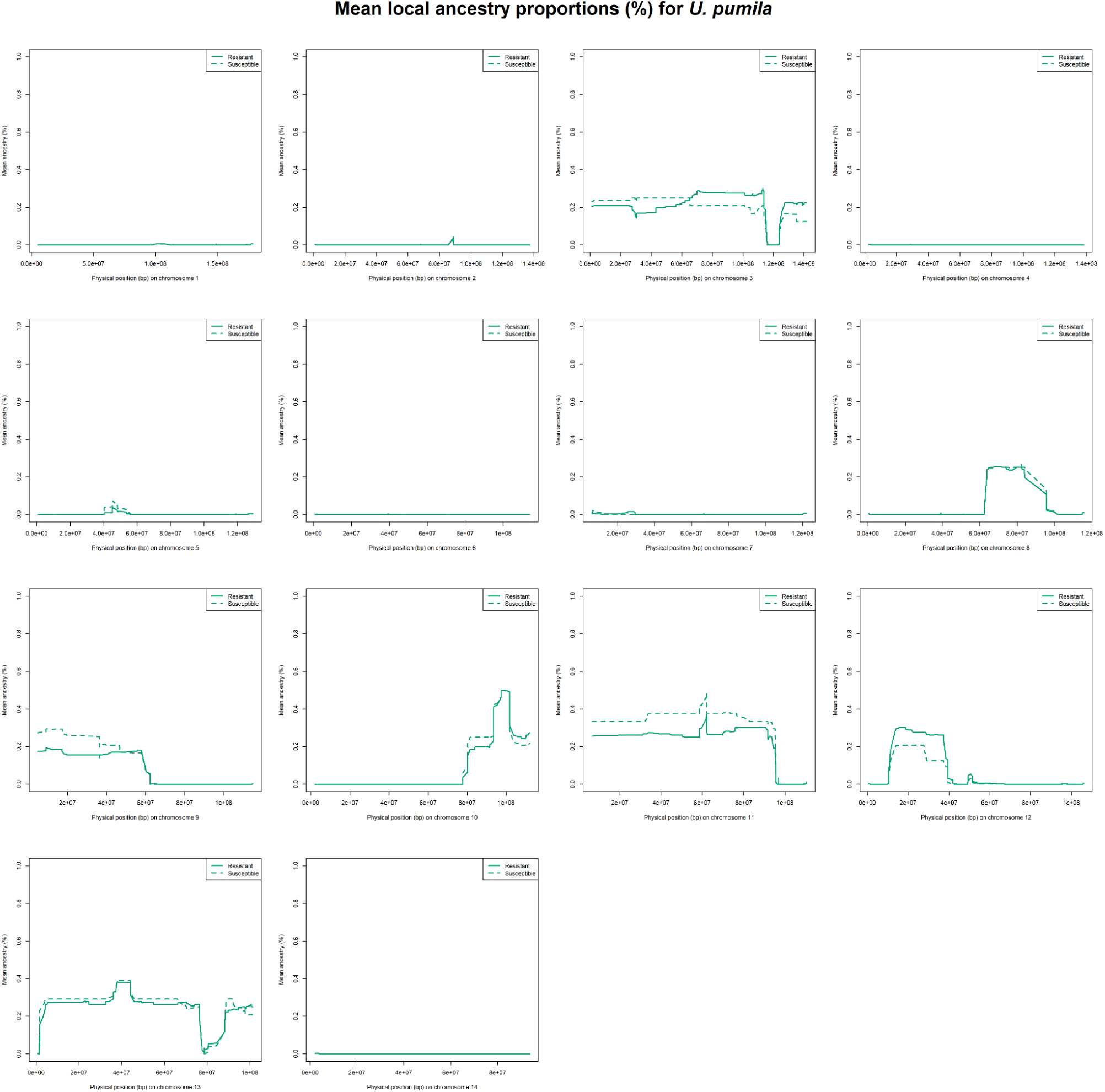
Mean local ancestry proportions (%) for *U. pumila*, comparing resistant WxTM progeny (solid line) with susceptible WxTM progeny (dashed line), based on a threshold of ≤25% defoliation, 1 year post-inoculation.

**Figure 11.**
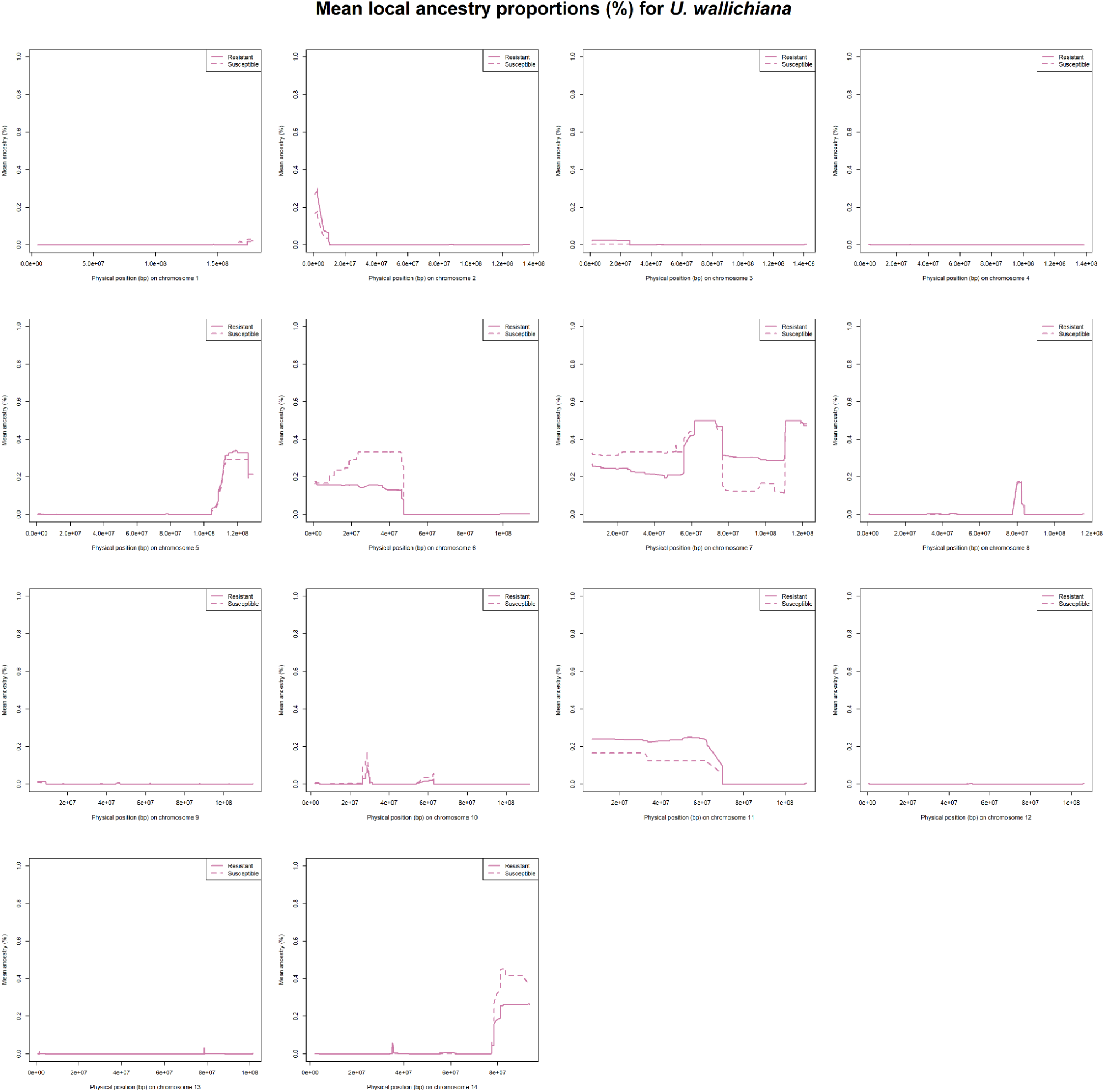
Mean local ancestry proportions (%) for *U. wallichiana*, comparing resistant WxTM progeny (solid line) with susceptible WxTM progeny (dashed line), based on a threshold of ≤25% defoliation, 1 year post-inoculation.

We did not find a significant association between the percentage of defoliation at 4, 8, and 12 weeks and one year after inoculation (Table 4, Table 7) and the percentage of global ancestry proportion of each parental species, namely *U. minor*, *U. glabra*, *U. pumila*, and *U. wallichiana* (Figures S26-S28).

**Table 7.**
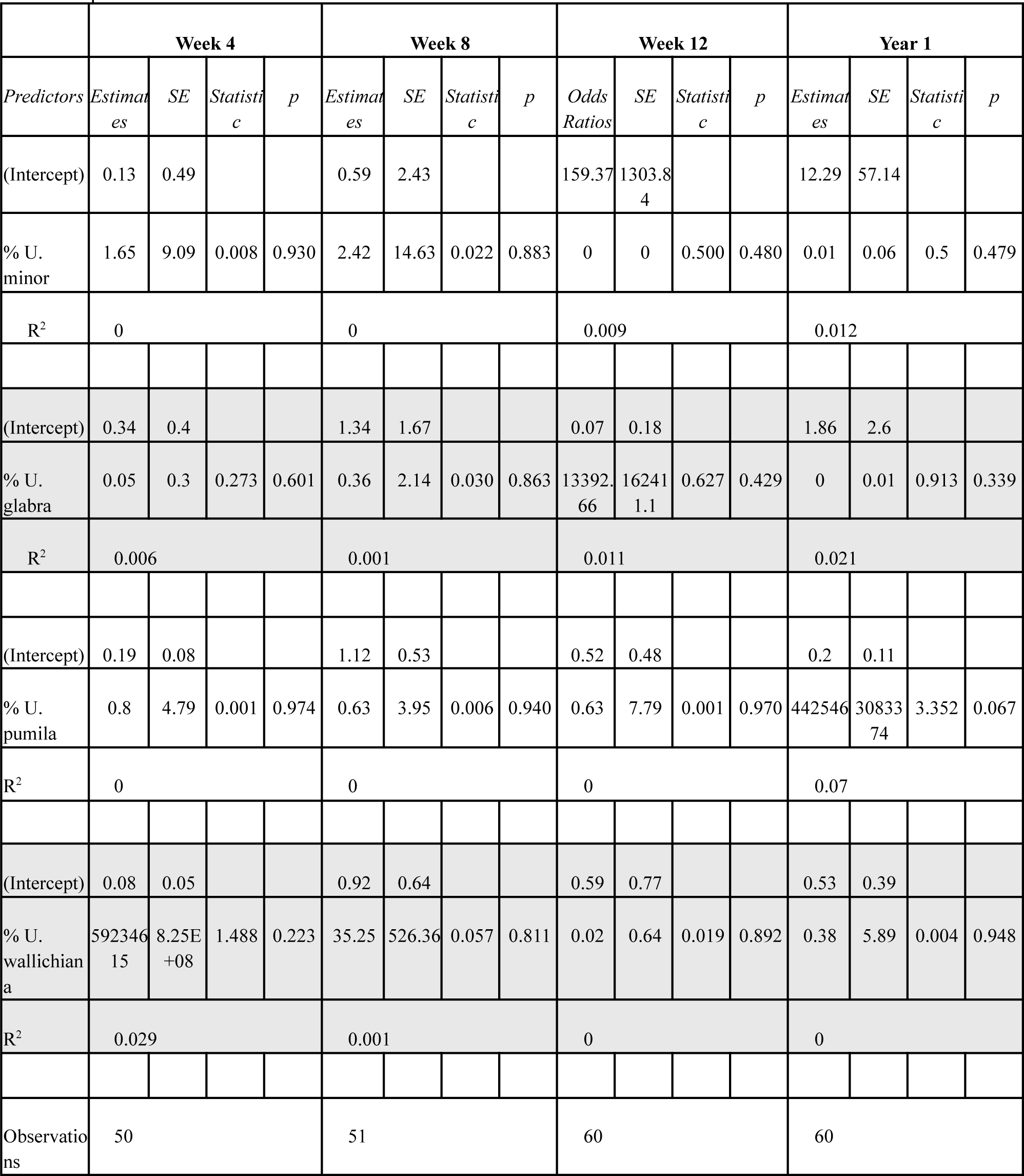
Summary statistics for model results investigating the relationship between defoliation scores and the percentage of the genome ascribed to one of the four parental *Ulmus* species.

|  | Week 4 |  |  |  | Week 8 |  |  |  | Week 12 |  |  |  | Year 1 |  |  |  |
| --- | --- | --- | --- | --- | --- | --- | --- | --- | --- | --- | --- | --- | --- | --- | --- | --- |
| Predictors | Estimates | SE | Statistic | p | Estimates | SE | Statistic | p | Odds Ratios | SE | Statistic | p | Estimates | SE | Statistic | p |
| (Intercept) | 0.13 | 0.49 |  |  | 0.59 | 2.43 |  |  | 159.37 | 1303.84 |  |  | 12.29 | 57.14 |  |  |
| % *U. minor* | 1.65 | 9.09 | 0.008 | 0.930 | 2.42 | 14.63 | 0.022 | 0.883 | 0 | 0 | 0.500 | 0.480 | 0.01 | 0.06 | 0.5 | 0.479 |
| R<sup>2</sup> | 0 |  |  |  | 0 |  |  |  | 0.009 |  |  |  | 0.012 |  |  |  |
| (Intercept) | 0.34 | 0.4 |  |  | 1.34 | 1.67 |  |  | 0.07 | 0.18 |  |  | 1.86 | 2.6 |  |  |
| % *U. glabra* | 0.05 | 0.3 | 0.273 | 0.601 | 0.36 | 2.14 | 0.030 | 0.863 | 13392.66 | 16241.1 | 0.627 | 0.429 | 0 | 0.01 | 0.913 | 0.339 |
| R<sup>2</sup> | 0.006 |  |  |  | 0.001 |  |  |  | 0.011 |  |  |  | 0.021 |  |  |  |
| (Intercept) | 0.19 | 0.08 |  |  | 1.12 | 0.53 |  |  | 0.52 | 0.48 |  |  | 0.2 | 0.11 |  |  |
| % *U. pumila* | 0.8 | 4.79 | 0.001 | 0.974 | 0.63 | 3.95 | 0.006 | 0.940 | 0.63 | 7.79 | 0.001 | 0.970 | 442546 | 3083374 | 3.352 | 0.067 |
| R<sup>2</sup> | 0 |  |  |  | 0 |  |  |  | 0 |  |  |  | 0.07 |  |  |  |
| (Intercept) | 0.08 | 0.05 |  |  | 0.92 | 0.64 |  |  | 0.59 | 0.77 |  |  | 0.53 | 0.39 |  |  |
| % *U. wallichiana* | 59234615 | 8.25E+08 | 1.488 | 0.223 | 35.25 | 526.36 | 0.057 | 0.811 | 0.02 | 0.64 | 0.019 | 0.892 | 0.38 | 5.89 | 0.004 | 0.948 |
| R<sup>2</sup> | 0.029 |  |  |  | 0.001 |  |  |  | 0 |  |  |  | 0 |  |  |  |
| Observations | 50 |  |  |  | 51 |  |  |  | 60 |  |  |  | 60 |  |  |  |

## Discussion

Here, we use the family structure of the ‘Wingham’ x Tonge Mill cross elms to develop the first linkage maps we are aware of for *Ulmaceae*. This confirms the accuracy of the scaffolding of the *U. glabra* genome assembly, showing no major rearrangements between that assembly and our data. Our linkage map was a prerequisite for the accuracy of our downstream analyses (Salter-Townshend and Myers 2019). Being able to model local recombination rates along each chromosome is essential for carrying out admixture analysis, a procedure which seeks to identify ancestry blocks that occur as a result of homologous recombination.

Our linkage maps represent the first step towards building accurate, high-density linkage maps. With only 60 WxTM progeny sequenced, the limited number of recombinations we were able to identify (<90 in some chromosomes) will have impaired the accuracy of the predicted recombination rates. Generating additional WxTM progeny from the same cross would significantly improve the linkage maps, and consequently the procedures which rely on them.

Using five samples for each of *U. minor*, *U. glabra*, and *U. pumila*, as well as one sample for *U. wallichiana* we were able to reconstruct the complete set of genomes for the WxTM cross elms and their parents in terms of haplotypes from each of the four parental species with relatively high confidence. The structure of the genome of FL493 (‘Wingham’) is notable because it is the product of several generations of breeding and careful screening for DED resistance. Although much of its parentage is known from crossing records, there are two open pollinations within this ancestry, so its full parentage is unknown. The extent to which chromosomal material from its parental species has been passed down was also previously unknown. Here, we estimate its genomic composition for the first time, showing that its father was likely also a complex hybrid. Wingham contains much *U. pumila* material on chromosomes 3, 8, 9, 10, 11, 12 and 13 and much *U. wallichana* material on chromosomes 6, 7, 11 and 14 with smaller fragments on chromosomes 2 and 5.

Plotting mean local ancestry for *U. pumila* and *U. wallichiana* separately for both resistant and susceptible elms (defined as ≤25% defoliation) allowed us to identify chromosomal regions potentially harbouring loci associated with resistance. Notably, a region roughly spanning 8 – 11 Mb on chromosome 7 was found to contain ∼15% more ancestry from *U. wallichiana* in resistant elms relative to susceptible elms, indicating that one or more genes in this region may contribute to DED resistance. Additional regions on chromosome 8 and chromosome 11 also displayed disparities in local ancestry between resistant and susceptible elms, with resistant elms demonstrating enhanced ancestry from *U. wallichiana* and *U. pumila* relative to susceptible elms in these regions.

If resistance to DED were a very highly polygenic trait, we might expect resistance to simply correlate with the percentage of the genome that is derived from resistant species (Westbrook et al. 2026). Plotting defoliation percentages against genome-wide proportions of ancestry from *U. minor*, *U. pumila*, and *U. wallichiana* revealed no strong correlation, suggesting that DED resistance is not a highly polygenic trait. Though no correlations were significant, the trends seen were the opposite of what we might have expected from a highly polygenic trait. Based on these initial results, we would anticipate that genes conferring DED resistance are limited in number, likely located in regions introgressed from *U. wallichiana* and *U. pumila*.

Our analyses are limited and should be regarded as preliminary. As mentioned above, our linkage maps likely contain inaccuracies due to the low number of progeny that we had to work with (many similar studies use hundreds of progeny), which will have also affected the ability of MOSAIC to identify ancestry switches. The low number of progeny analysed also limits the resolution with which we can identify chromosomal regions of interest. If we had hundreds of progeny, we would have far more recombination events breaking up regions of progenitor genomes, allowing more precise knowledge of the location of loci associated with resistance.

We also had only small sample sizes of the parental species for the admixture analysis, especially of *U. pumila* and *U. wallichiana*. This means we were not able to capture all the alleles from these species that could be present in FL493 (as we do not have the exact genotypes of the parental species used in its ancestry). By sequencing additional elms from these species, our reference panels could better capture the full genetic diversity of parental species, and thus increase the ability of admixture analysis to distinguish between haplotypes from each species. Alternatively, if we were able to sequence the exact progenitor genotypes, we could have even higher confidence in our results. The level of genomic distance between species also affects our ability to differentiate them in admixture analysis. As *U. minor* is closely related to *U. glabra* (Whittemore et al. 2021) these will be particularly hard to distinguish. These limitations may be why the Tonge Mill elms appeared to have some *U. glabra* and a very small amount of *U. wallichiana* within them.

In our analysis we relied upon mapping all sequence reads to the reference genome for *U. glabra* as this is the best reference assembly available for the genus. It is likely that the other parental species contain segments of sequence that are entirely absent from *U. glabra*. This is particularly likely for *U. pumila* and *U. wallichiana* which are distant relatives. Genes conferring resistance to DED may have introgressed into the WxTM progeny that are missing from the *U. glabra* genome. Sequence reads from such genes would not have mapped to the *U. glabra* reference, and would have been discarded. Our ability to find such genes would be greatly enhanced by the assembly and annotation of chromosome level genomes for *U. pumila* and *U. wallichiana*. If we had these, we could inspect the regions of interest in chromosomes 7, 8 and 11 detected in the present study, seeking genes in these regions that are present in *U. pumila* or *U. wallichiana* but not in *U. glabra* and *U. minor*.

Our analysis greatly benefited from the long history of breeding for DED-resistance (Martín et al. 2019; Mittempergher and Santini 2004; Holmes and Heybroek 1990), meaning that a family containing five generations of crossing was available for analysis. As recombination happens once per generation, the higher the number of generations leads to more recombination events, and smaller blocks of ancestry within the genomes. Had it not been for this, the areas of interest identified in our admixture mapping would have been larger. If more generations were available, the ancestry blocks would become even smaller, and the results more precise. Thus, it is highly desirable to make F2 progeny by crossing WxTM trees, and use these to make large mapping populations.

Moreover, though it is conventional to employ admixture analysis to identify chromosomal segments harbouring genes of interest, another direction for future analysis would be to conduct a local ancestry-aware GWAS using tools such as MIXSCORE or TRACTOR (Atkinson et al. 2021). Admixture is often cited as a source of false positives in GWAS, but incorporating information about local ancestry can substantially improve the power of GWAS (Pasaniuc et al. 2011). This would enable us to identify SNPs which potentially contribute to DED resistance, allowing us to exploit local ancestry inference while circumventing the issue of extensive admixture linkage disequilibrium.

Our findings demonstrate that admixture mapping has the potential to significantly enhance disease mapping approaches in the context of DED. Moreover, our results support the idea that resistance to DED is conferred by a small number of loci, with no clear correlation between defoliation and ancestry from *U. minor* or any other elm species analysed herein. We were able to identify genomic regions of interest – particularly one region on chromosome 7 – which demonstrated enhanced ancestry from *U. wallichiana* and *U. pumila* in more resistant trees, that could potentially harbour genes conferring heightened resistance to DED. However, the accuracy and statistical power of our analysis is limited by a number of factors, largely related to small sample sizes.

## Data availability

The raw DNA sequences are available from GenBank under project number PRJNA982084.

## Acknowledgements

This project was funded by Defra under the Center for Forest Protection (Ref. 2967). We are grateful to the Department for Environment, Food & Rural Affairs staff for their support. We are also grateful for the knowledge sharing of David Macaya Sanz. We used Queen Mary’s Apocrita computing cluster (http://doi.org/10.5281/zenodo.438045) for data analysis, and their Research-IT team’s support is much appreciated.

## Author contributions

Catherine Gudgeon carried out the genomic analyses from read mapping to data analysis. Mohammad Vatanparast oversaw DNA extractions and read mapping to SNP calling. Rômulo Carleial contributed statistical analyses. David Herling conceived the cross and carried it out, and conducted inoculations and phenotype scoring. Fergus Poncia planted the progeny and conducted inoculations and phenotype scoring. Alberto Santini supplied biological materials and information on ancestry. Clive Brasier and Joan F. Webber supplied inoculum and designed the inoculation protocol. Richard Buggs conceived the analysis, collected materials for DNA extraction and oversaw the project. All living authors wrote the manuscript.

## Supplementary information

### Supplementary Tables

### Supplementary Figures

**Figure S1a.**
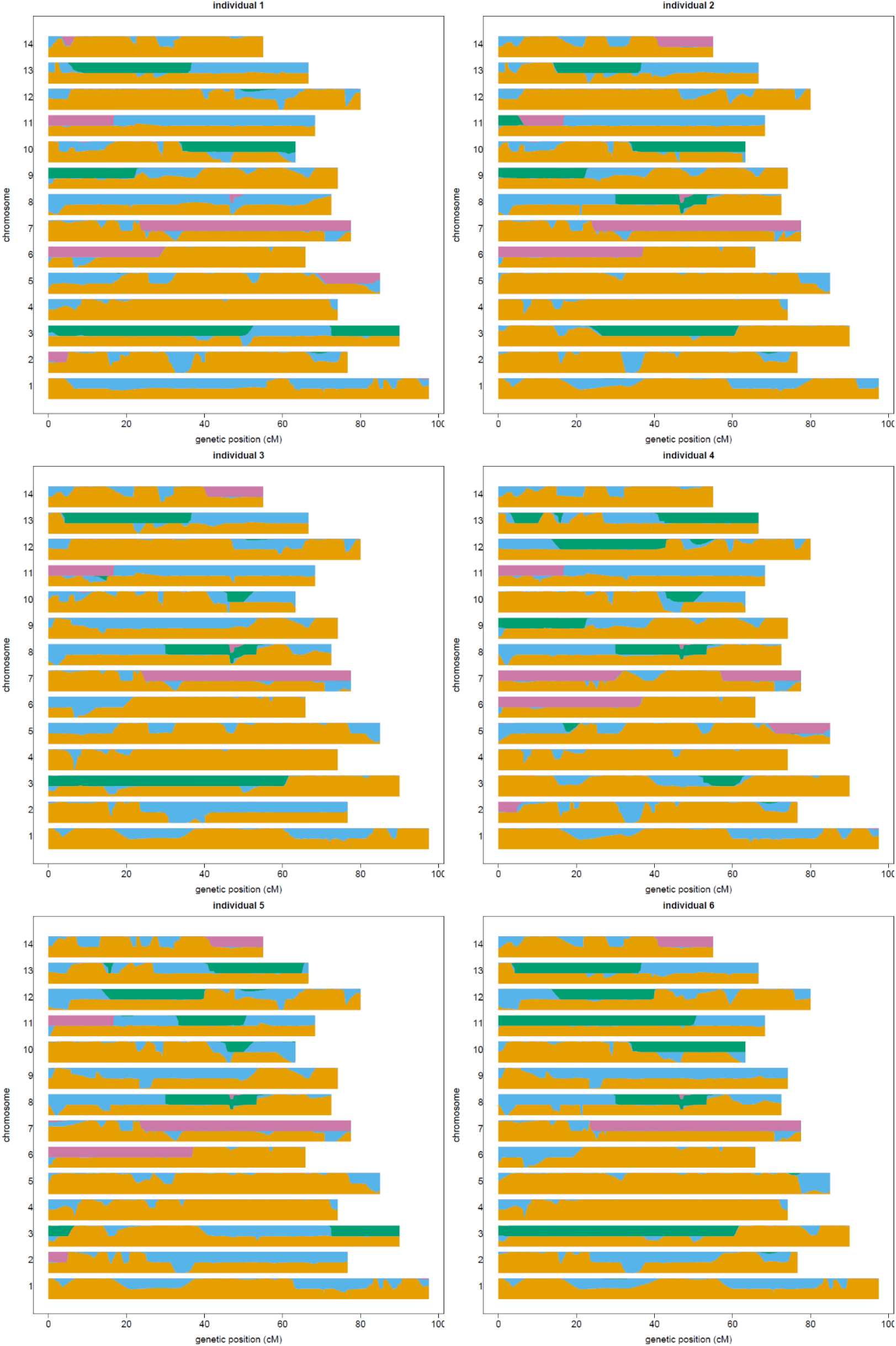
Karyograms for individuals 1-6, i.e. E001, E002, E003, E004, E005, and E006 respectively. Y-axis values for each chromosome indicate the estimated number of alleles from each ancestral species; genetic distance (cM) is plotted along the x-axis. These individuals are all admixed WxTM progeny of FL493 × *U. minor* (Tonge Mill). Local ancestry assignments are indicated by colour; orange = *U. minor*, blue = *U. glabra*, green = *U. pumila*, pink = *U. wallichiana*.

**Figure S1b.**
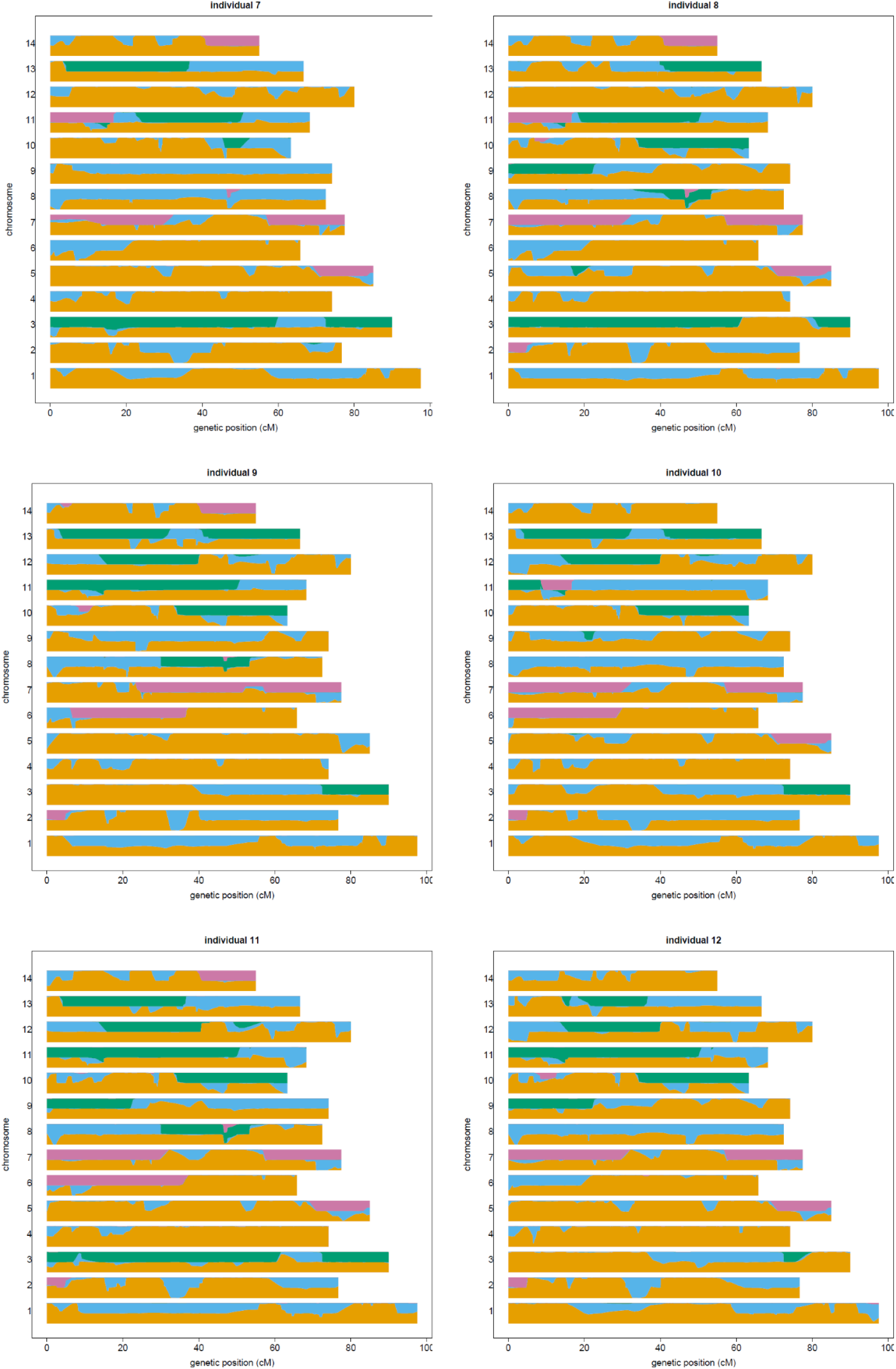
Karyograms for E007, E008, E009, E010, E011, and E012 respectively. Y-axis values for each chromosome indicate the estimated number of alleles from each ancestral species; genetic distance (cM) is plotted along the x-axis. These individuals are all admixed WxTM progeny of FL493 × *U. minor* (Tonge Mill). Local ancestry assignments are indicated by colour; orange = *U. minor*, blue = *U. glabra*, green = *U. pumila*, pink = *U. wallichiana*.

**Figure S1c.**
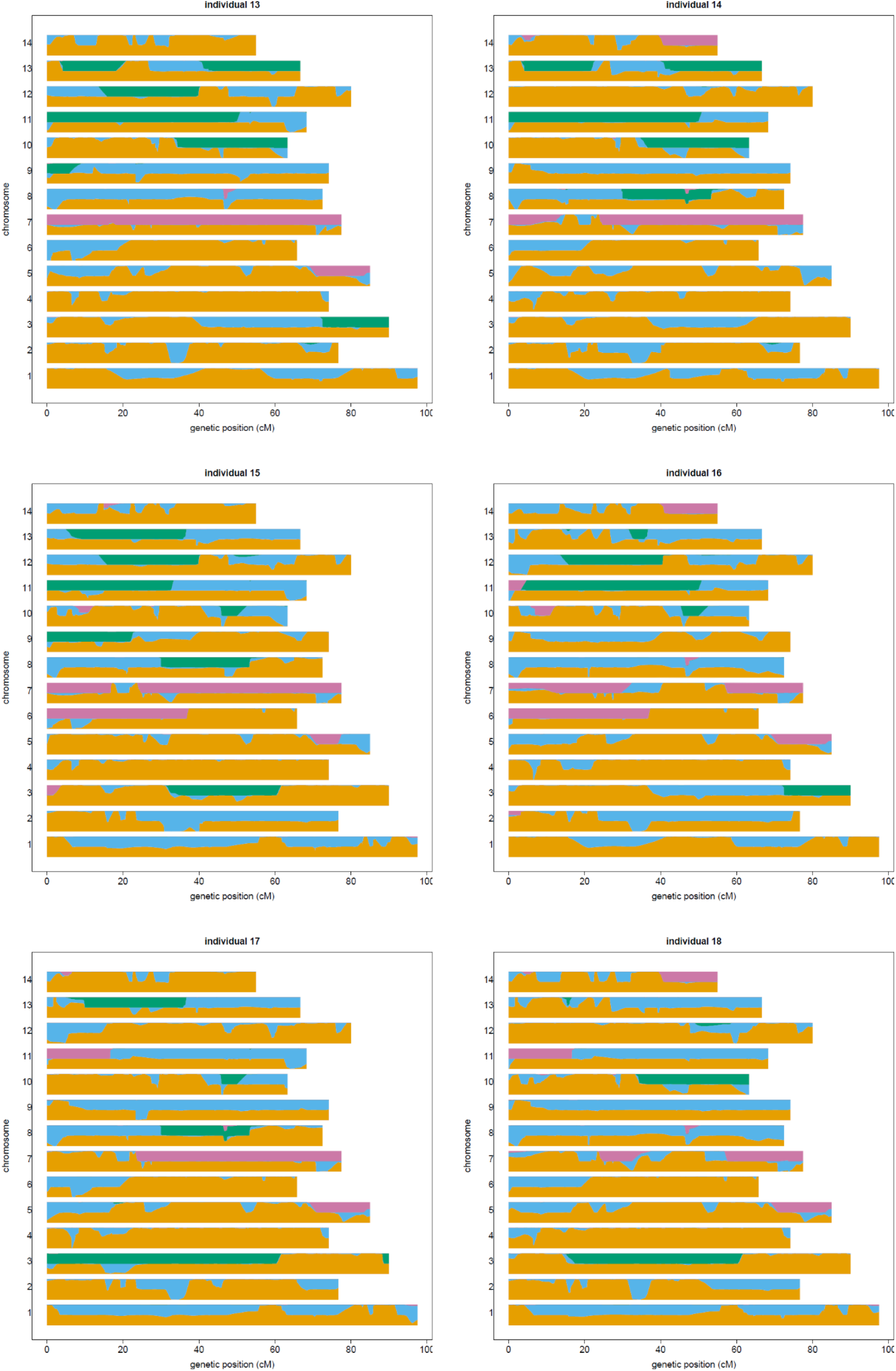
Karyograms for E013, E014, E015, E016, E017, and E018 respectively. Y-axis values for each chromosome indicate the estimated number of alleles from each ancestral species; genetic distance (cM) is plotted along the x-axis. These individuals are all admixed WxTM progeny of FL493 × *U. minor* (Tonge Mill). Local ancestry assignments are indicated by colour; orange = *U. minor*, blue = *U. glabra*, green = *U. pumila*, pink = *U. wallichiana*.

**Figure S1d.**
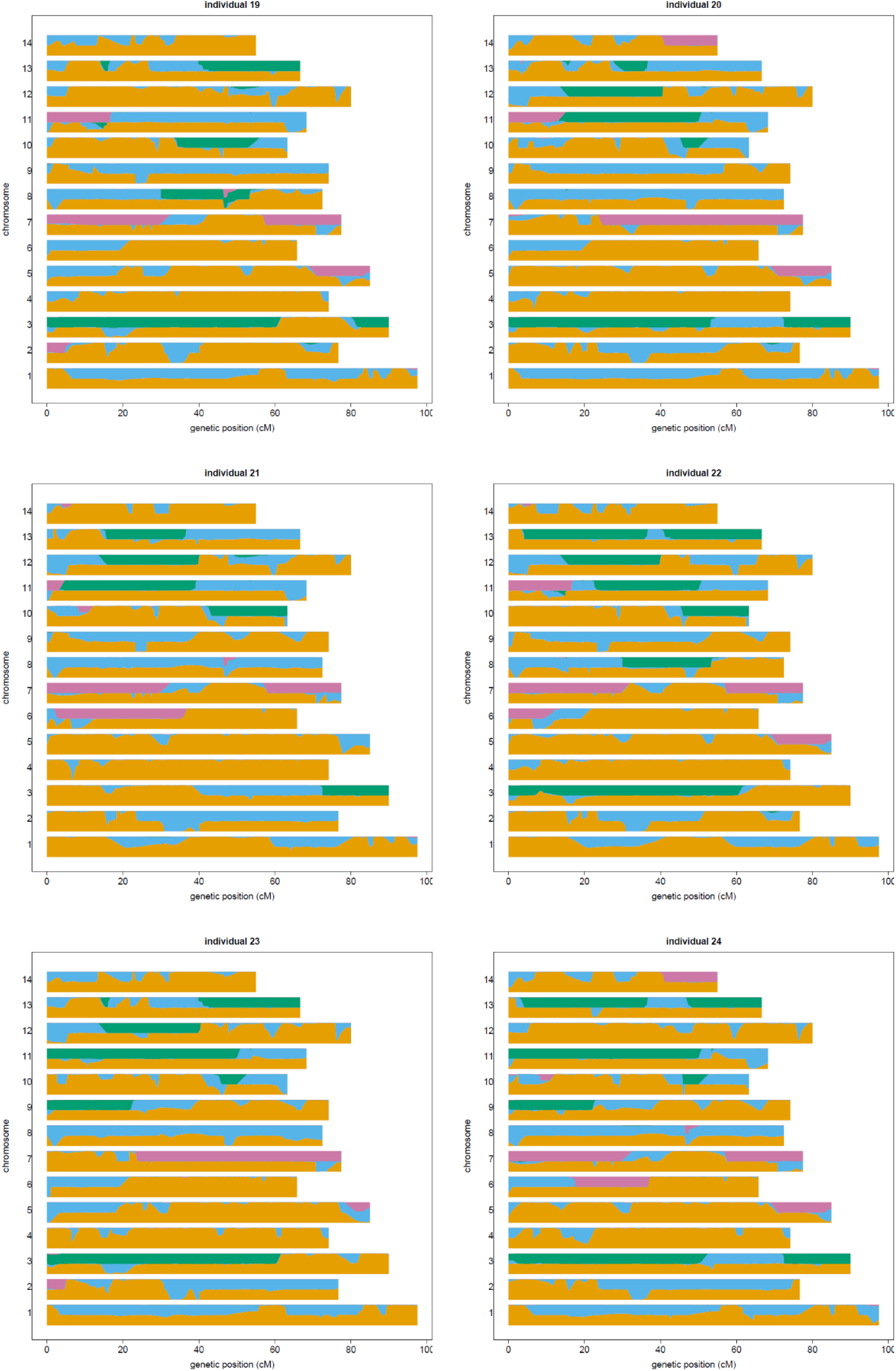
Karyograms for E019, E020, E021, E022, E023, and E024 respectively. Y-axis values for each chromosome indicate the estimated number of alleles from each ancestral species; genetic distance (cM) is plotted along the x-axis. These individuals are all admixed WxTM progeny of FL493 × *U. minor* (Tonge Mill). Local ancestry assignments are indicated by colour; orange = *U. minor*, blue = *U. glabra*, green = *U. pumila*, pink = *U. wallichiana*.

**Figure S1e.**
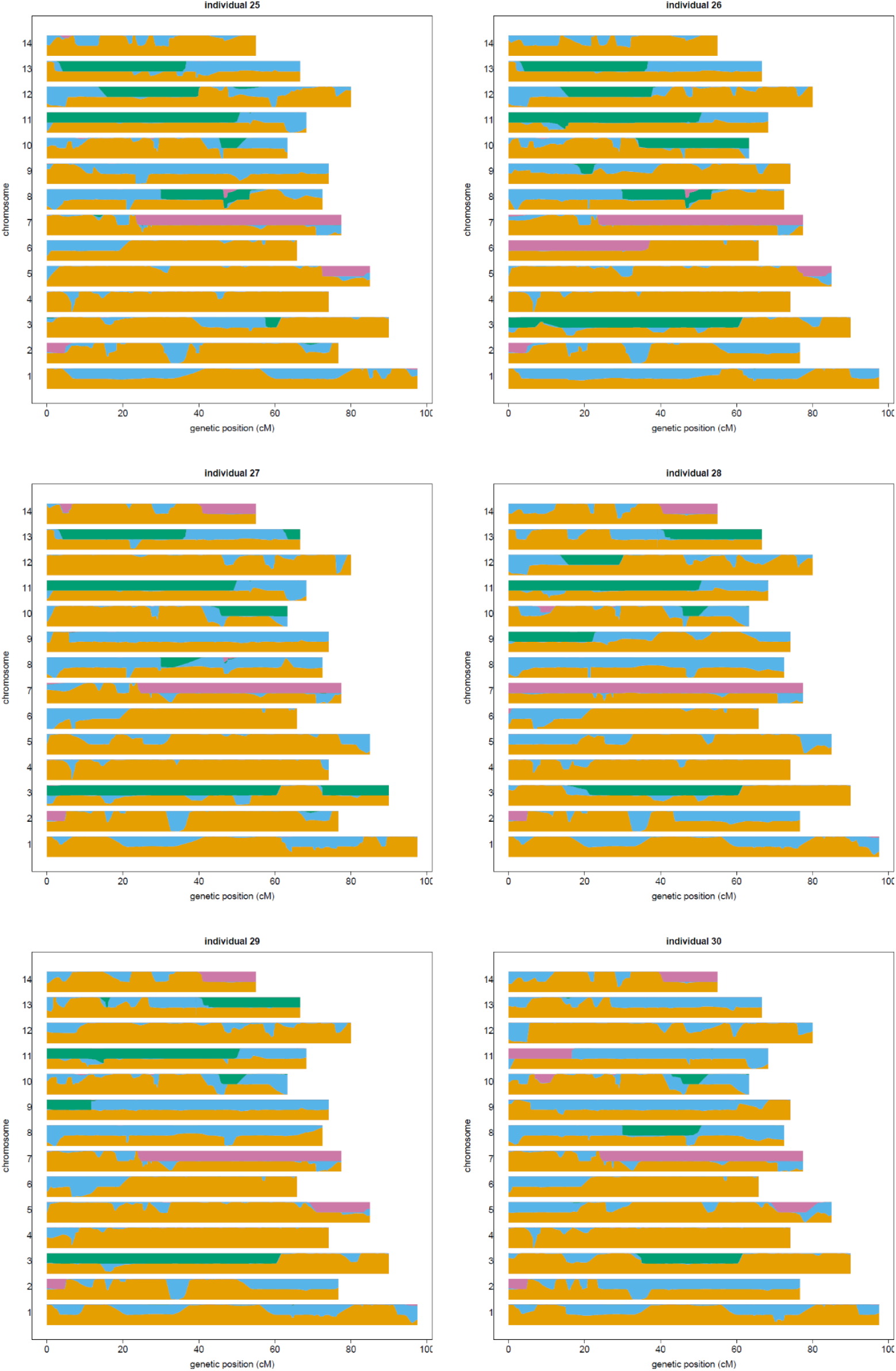
Karyograms for E025, E026, E027, E028, E029, and E030 respectively. Y-axis values for each chromosome indicate the estimated number of alleles from each ancestral species; genetic distance (cM) is plotted along the x-axis. These individuals are all admixed WxTM progeny of FL493 × *U. minor* (Tonge Mill). Local ancestry assignments are indicated by colour; orange = *U. minor*, blue = *U. glabra*, green = *U. pumila*, pink = *U. wallichiana*.

**Figure S1f.**
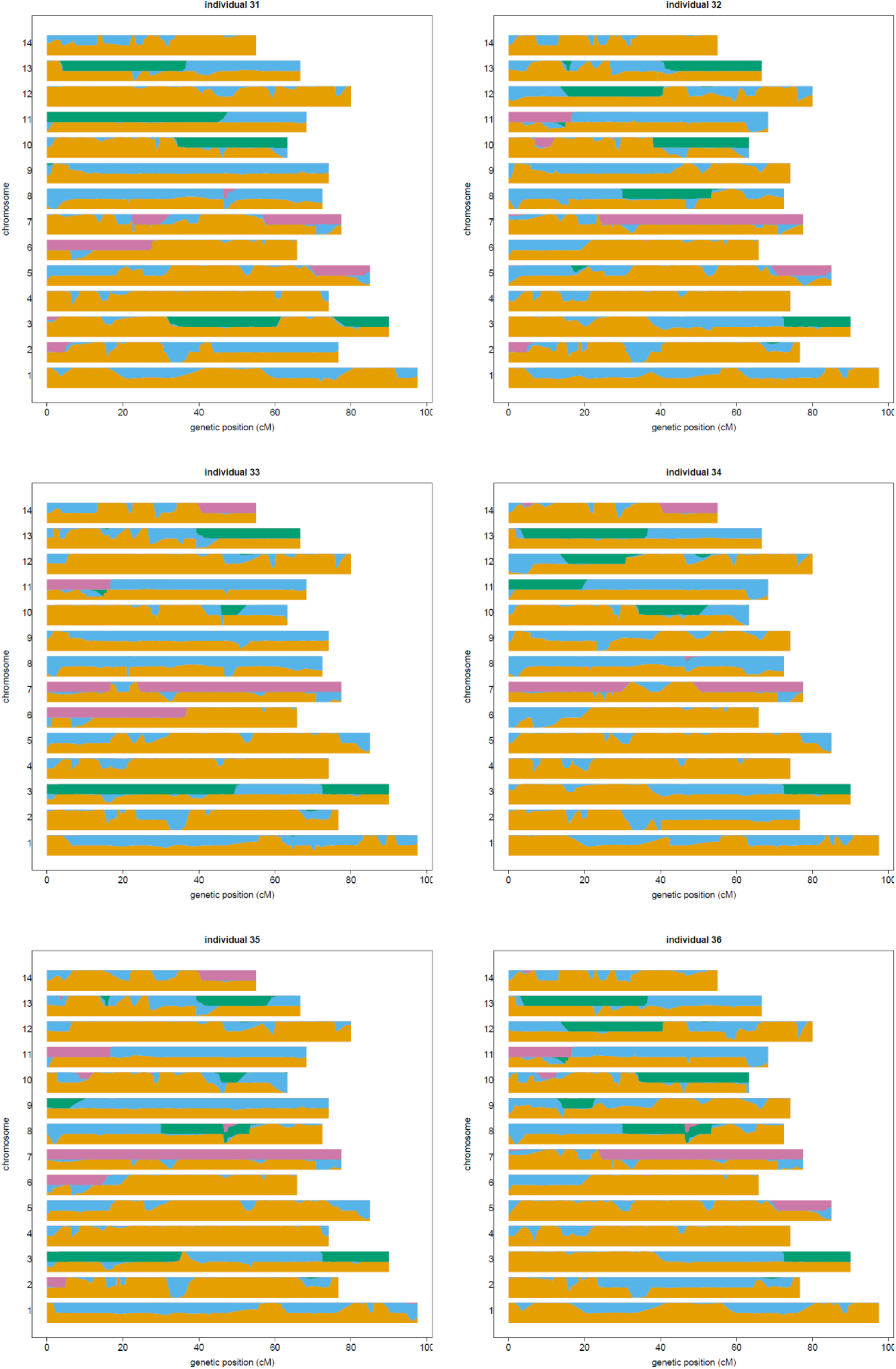
Karyograms for E031, E032, E033, E034, E035, and E036 respectively. Y-axis values for each chromosome indicate the estimated number of alleles from each ancestral species; genetic distance (cM) is plotted along the x-axis. These individuals are all admixed WxTM progeny of FL493 × *U. minor* (Tonge Mill). Local ancestry assignments are indicated by colour; orange = *U. minor*, blue = *U. glabra*, green = *U. pumila*, pink = *U. wallichiana*.

**Figure S1g.**
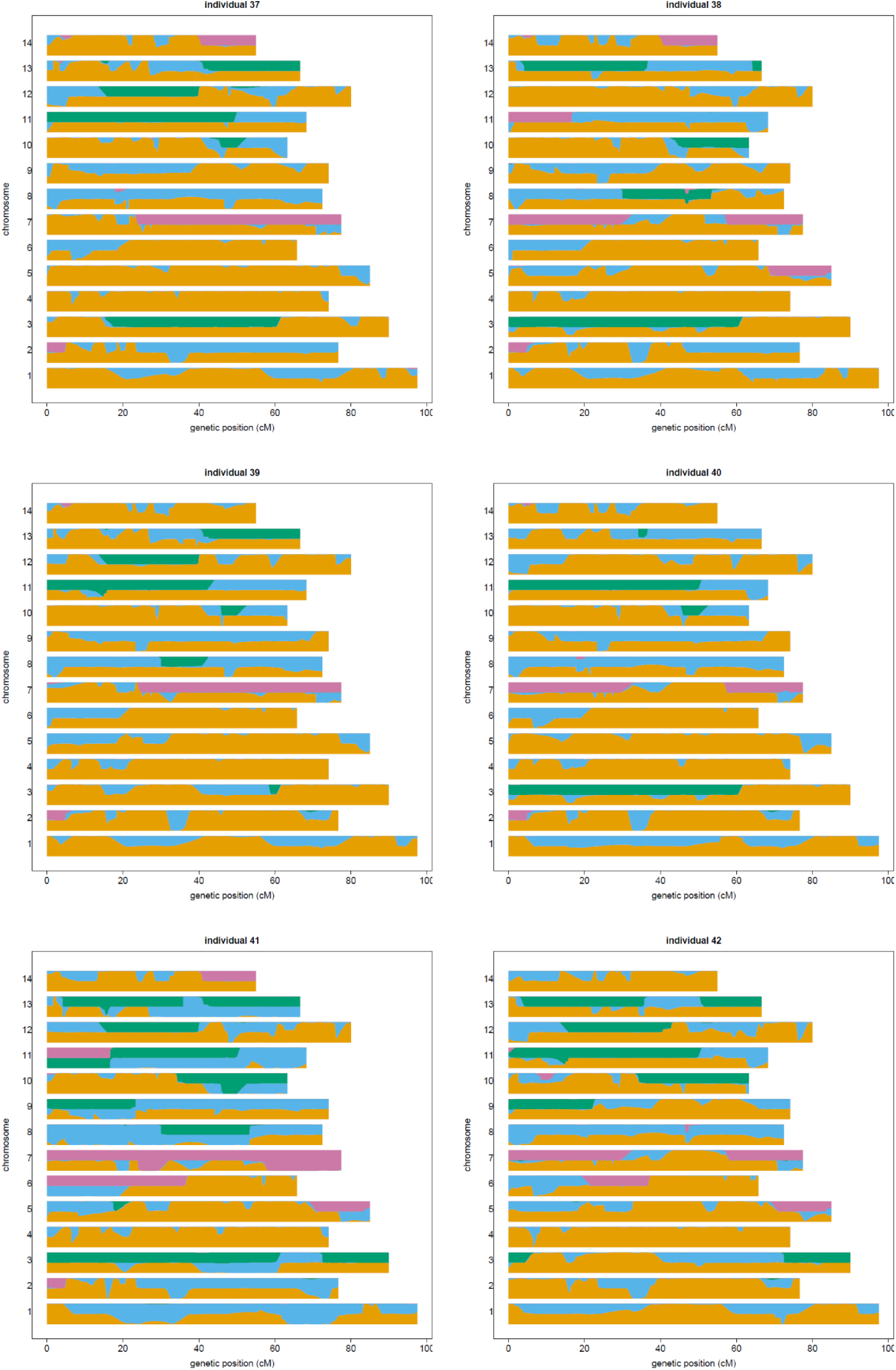
Karyograms for E037, E038, E039, E040, E041, and E265 respectively. Y-axis values for each chromosome indicate the estimated number of alleles from each ancestral species; genetic distance (cM) is plotted along the x-axis. With the exception of individual 41 (E041, ‘Wingham’) these individuals are all admixed WxTM progeny of FL493 × *U. minor* (Tonge Mill). Local ancestry assignments are indicated by colour; orange = *U. minor*, blue = *U. glabra*, green = *U. pumila*, pink = *U. wallichiana*.

**Figure S1h.**
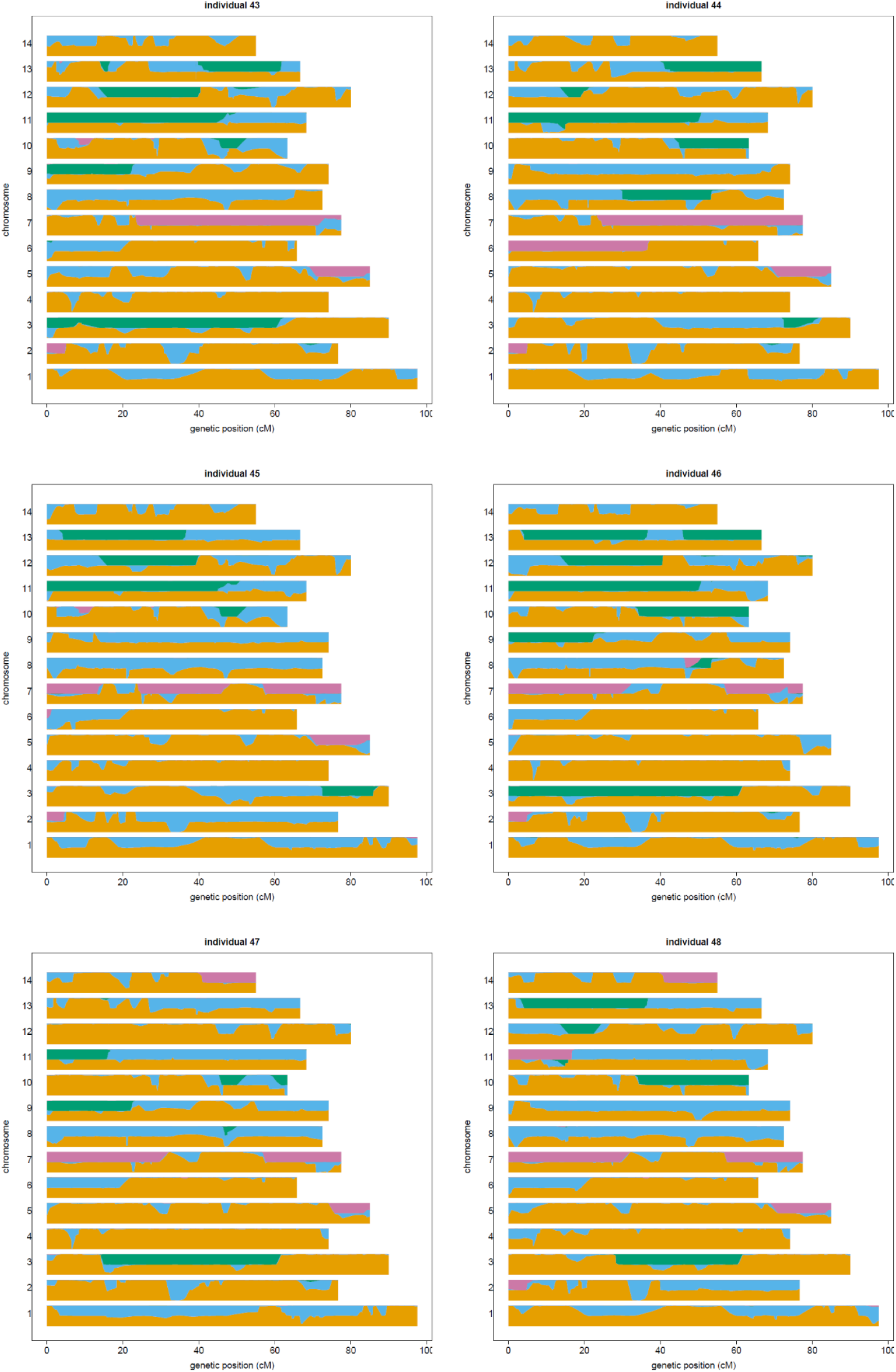
Karyograms for E266, E267, E268, E269, E270, and E271 respectively. Y-axis values for each chromosome indicate the estimated number of alleles from each ancestral species; genetic distance (cM) is plotted along the x-axis. These individuals are all admixed WxTM progeny of FL493 × *U. minor* (Tonge Mill). Local ancestry assignments are indicated by colour; orange = *U. minor*, blue = *U. glabra*, green = *U. pumila*, pink = *U. wallichiana*.

**Figure S1i.**
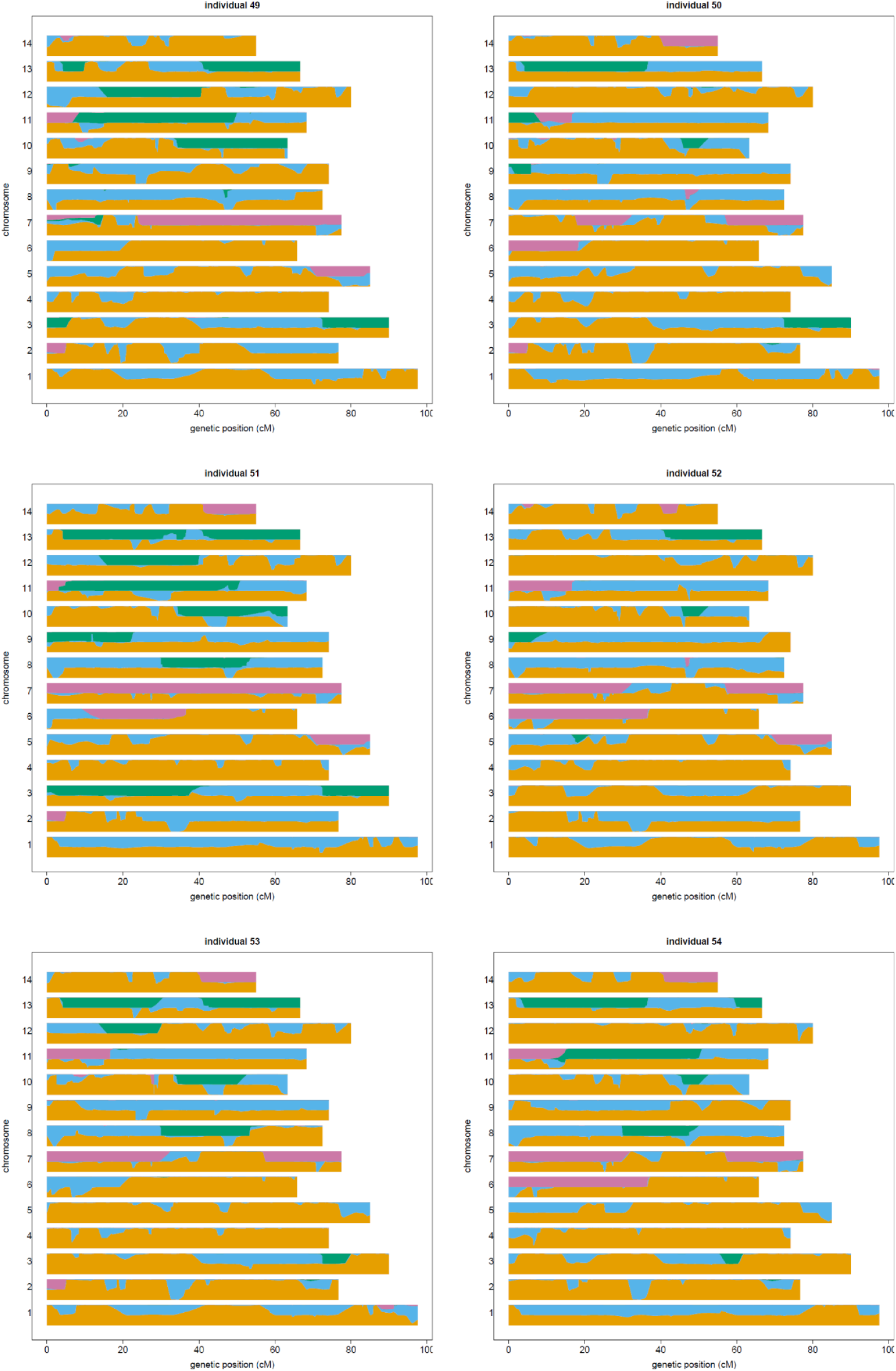
Karyograms for E272, E273, E274, E275, E276, and E277 respectively. Y-axis values for each chromosome indicate the estimated number of alleles from each ancestral species; genetic distance (cM) is plotted along the x-axis. These individuals are all admixed WxTM progeny of FL493 × *U. minor* (Tonge Mill). Local ancestry assignments are indicated by colour; orange = *U. minor*, blue = *U. glabra*, green = *U. pumila*, pink = *U. wallichiana*.

**Figure S1j.**
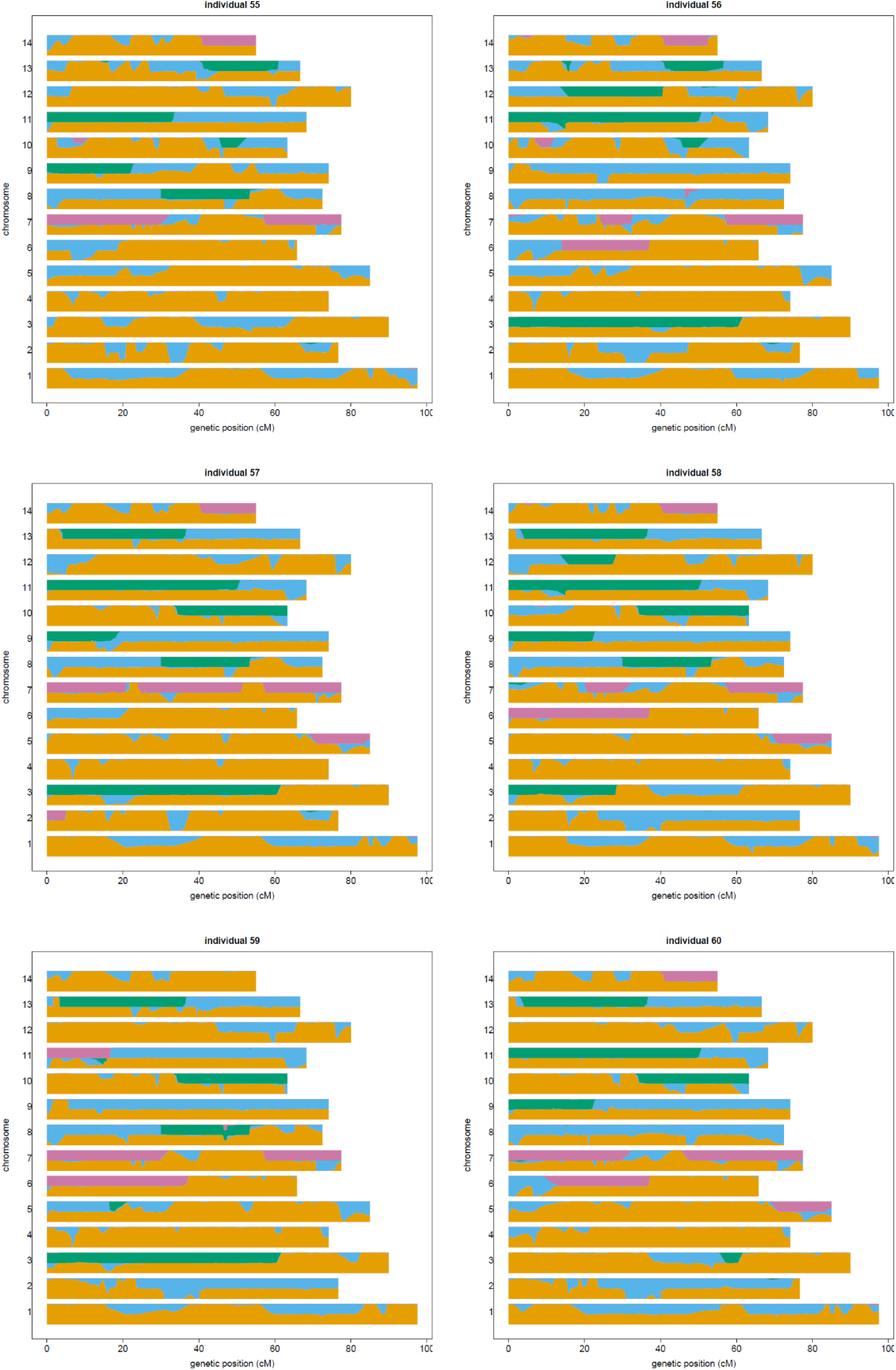
Karyograms for E278, E279, E280, E281, E282, and E283 respectively. Y-axis values for each chromosome indicate the estimated number of alleles from each ancestral species; genetic distance (cM) is plotted along the x-axis. These individuals are all admixed WxTM progeny of FL493 × *U. minor* (Tonge Mill). Local ancestry assignments are indicated by colour; orange = *U. minor*, blue = *U. glabra*, green = *U. pumila*, pink = *U. wallichiana*.

**Figure S1k.**
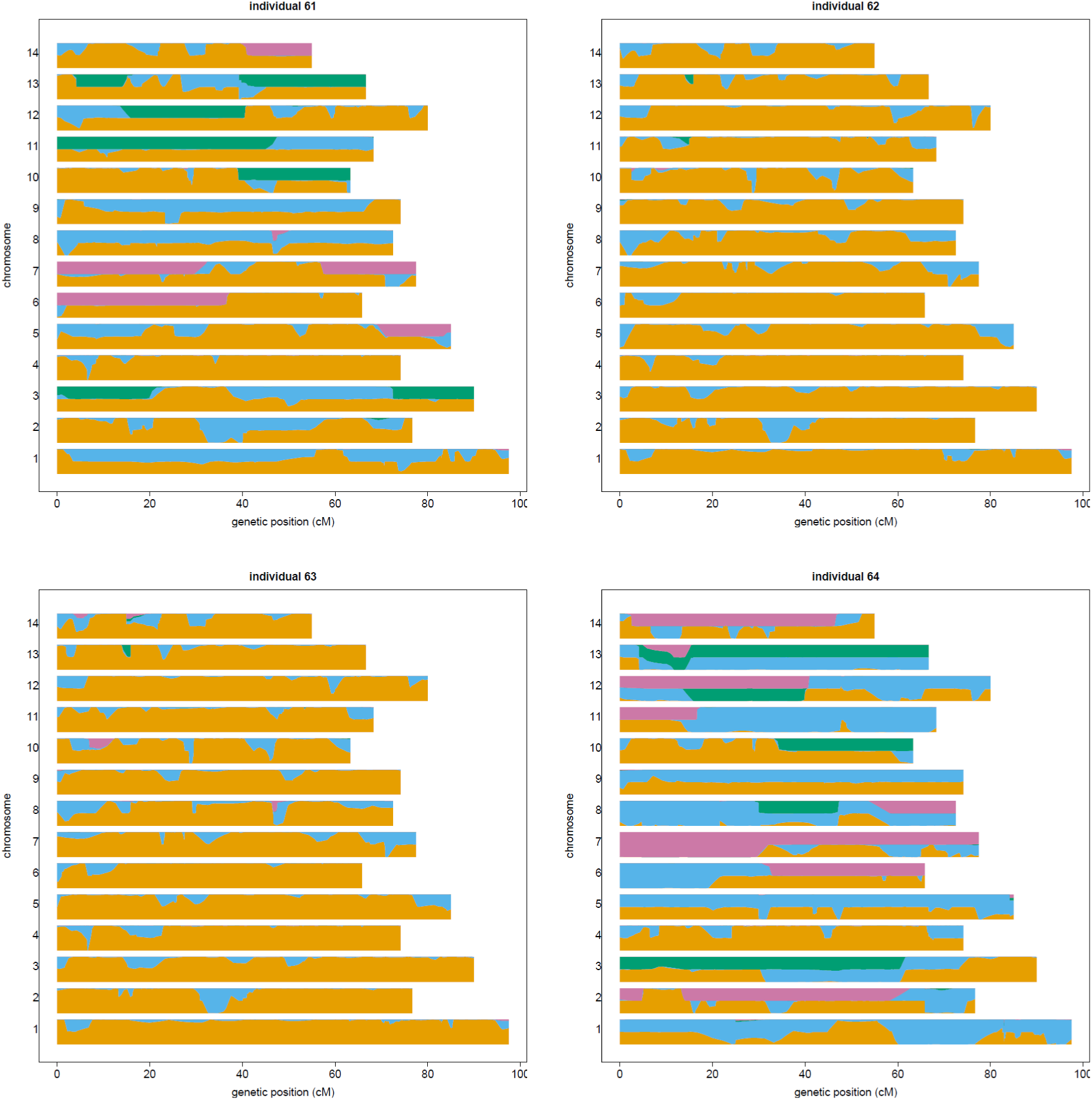
Karyograms for E284, E288, E289 and E291 respectively. E284 is WxTM progeny of the FL493 × *U. minor* (Tonge Mill). E288 and E289 are clones of *U. minor* at Tonge Mill. E291 is the mother of ‘Wingham’. Y-axis values for each chromosome indicate the estimated number of alleles from each ancestral species; genetic distance (cM) is plotted along the x-axis. Local ancestry assignments are indicated by colour; orange = *U. minor*, blue = *U. glabra*, green = *U. pumila*, pink = *U. wallichiana*.

**Figure S2.**
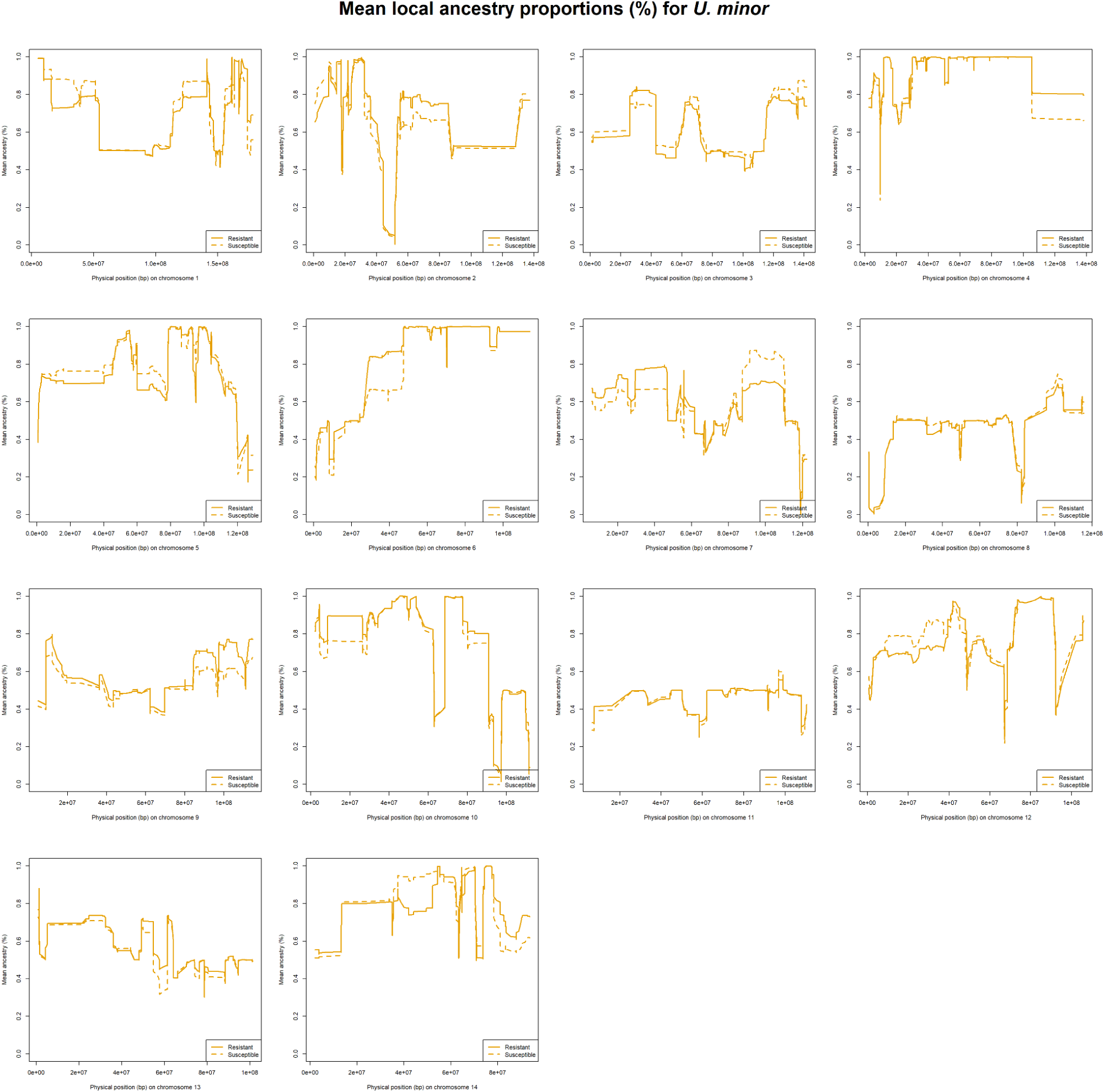
Mean local ancestry proportions (%) for *U. minor*, comparing resistant WxTM progeny (solid line) with susceptible WxTM progeny (dashed line), based on a threshold of ≤25% defoliation, 4 weeks post-inoculation. Defoliation measurements were only available for 50/60 of the WxTM progeny during this timeframe.

**Figure S3.**
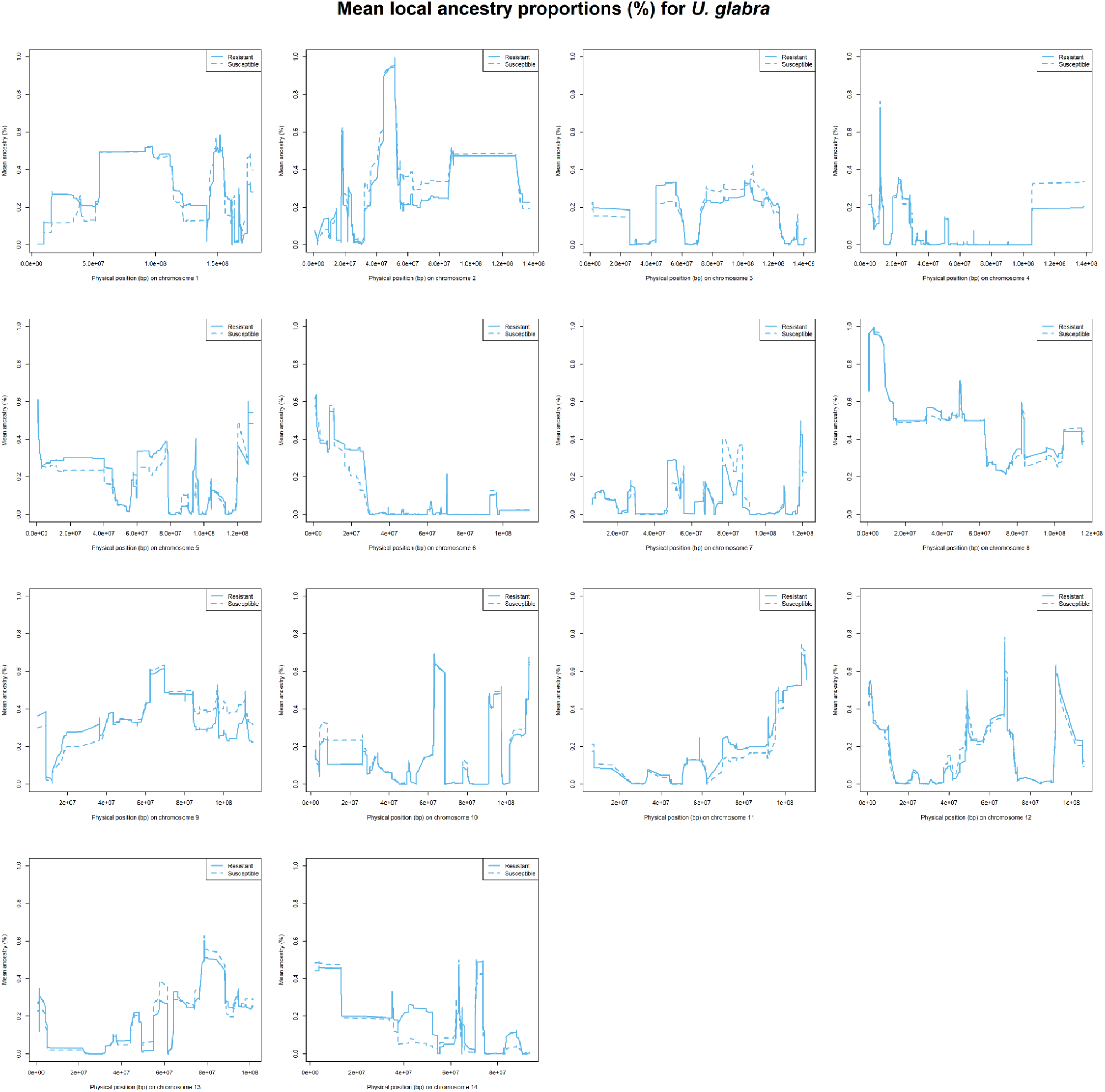
Mean local ancestry proportions (%) for *U. glabra*, comparing resistant WxTM progeny (solid line) with susceptible WxTM progeny (dashed line), based on a threshold of ≤25% defoliation, 4 weeks post-inoculation. Defoliation measurements were only available for 50/60 of the WxTM progeny during this timeframe.

**Figure S4.**
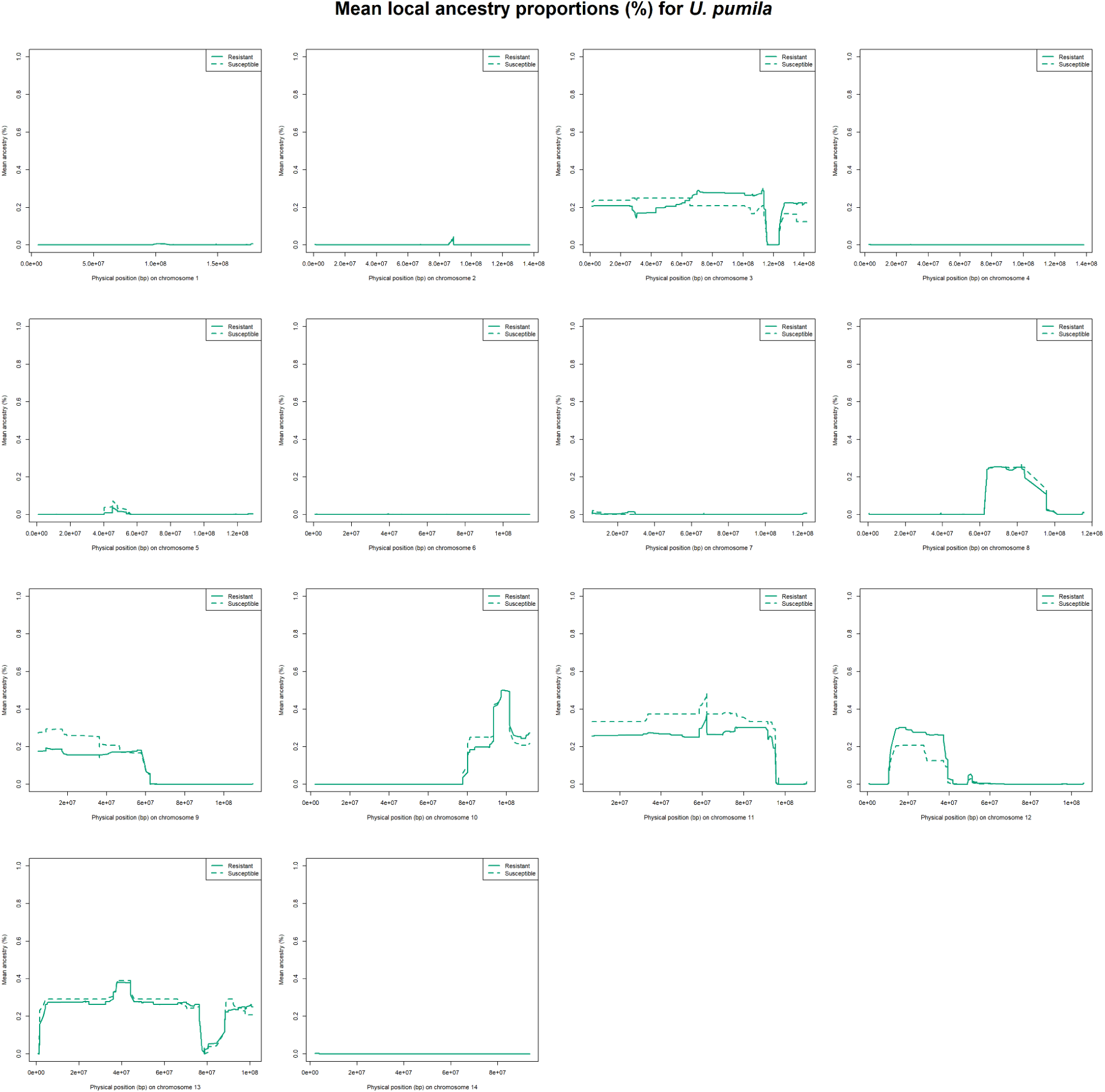
Mean local ancestry proportions (%) for *U. pumila*, comparing resistant WxTM progeny (solid line) with susceptible WxTM progeny (dashed line), based on a threshold of ≤25% defoliation, 4 weeks post-inoculation. Defoliation measurements were only available for 50/60 of the WxTM progeny during this timeframe.

**Figure S5.**
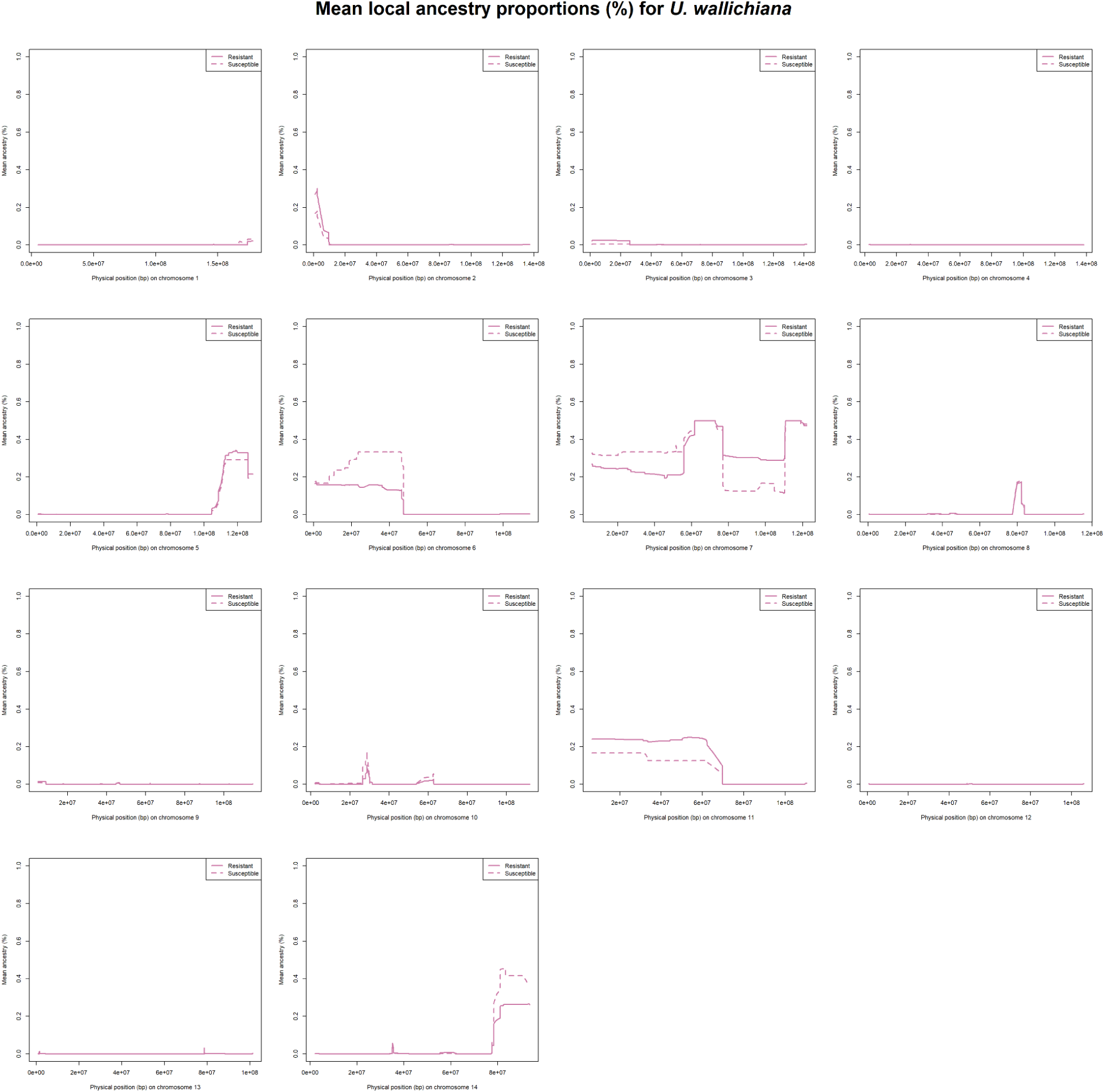
Mean local ancestry proportions (%) for *U. wallichiana*, comparing resistant WxTM progeny (solid line) with susceptible WxTM progeny (dashed line), based on a threshold of ≤25% defoliation, 4 weeks post-inoculation. Defoliation measurements were only available for 50/60 of the WxTM progeny during this timeframe.

**Figure S6.**
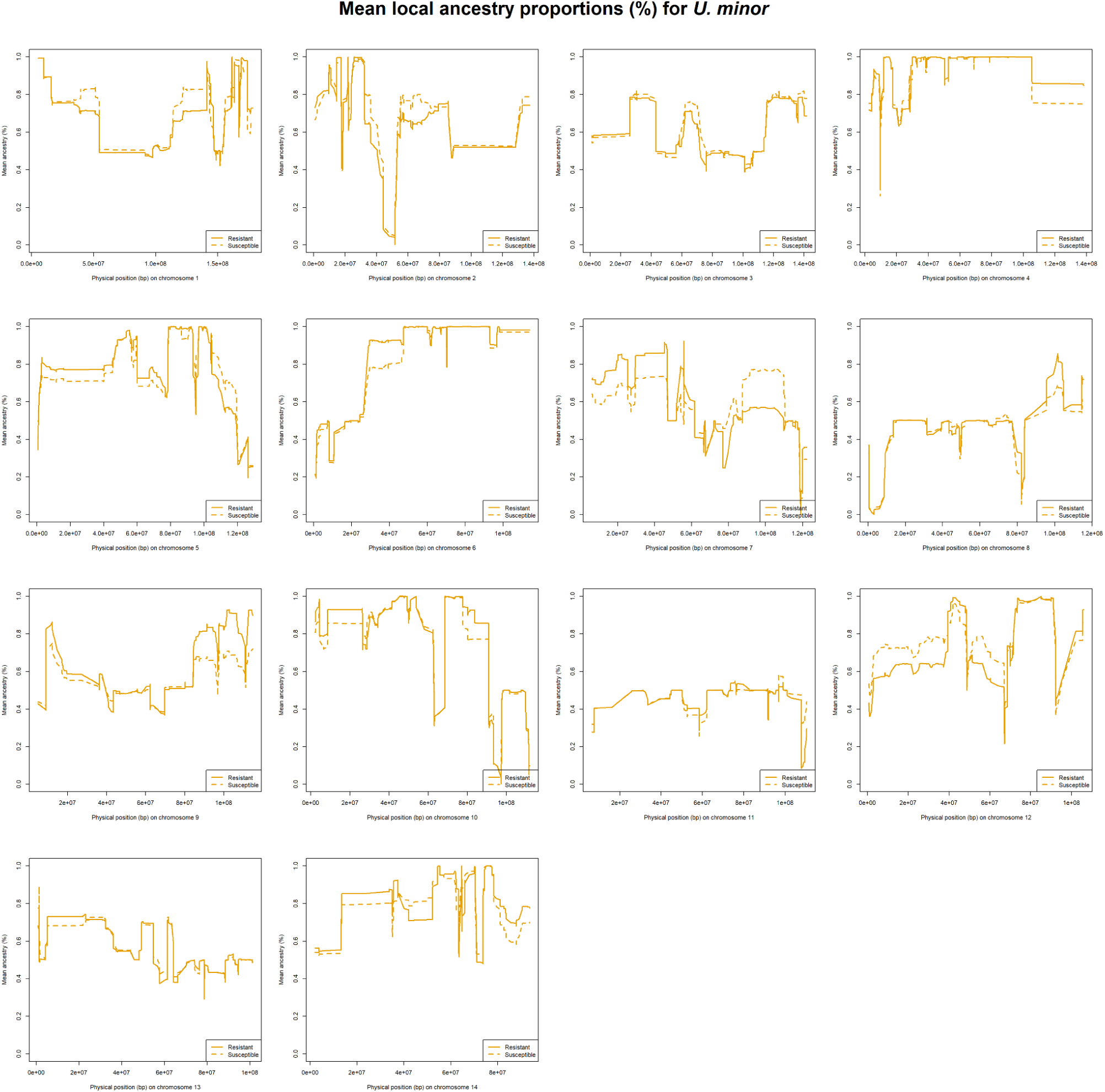
Mean local ancestry proportions (%) for *U. minor*, comparing resistant WxTM progeny (solid line) with susceptible WxTM progeny (dashed line), based on a threshold of ≤25% defoliation, 8 weeks post-inoculation. Defoliation measurements were only available for 51/60 of the WxTM progeny during this timeframe.

**Figure S7.**
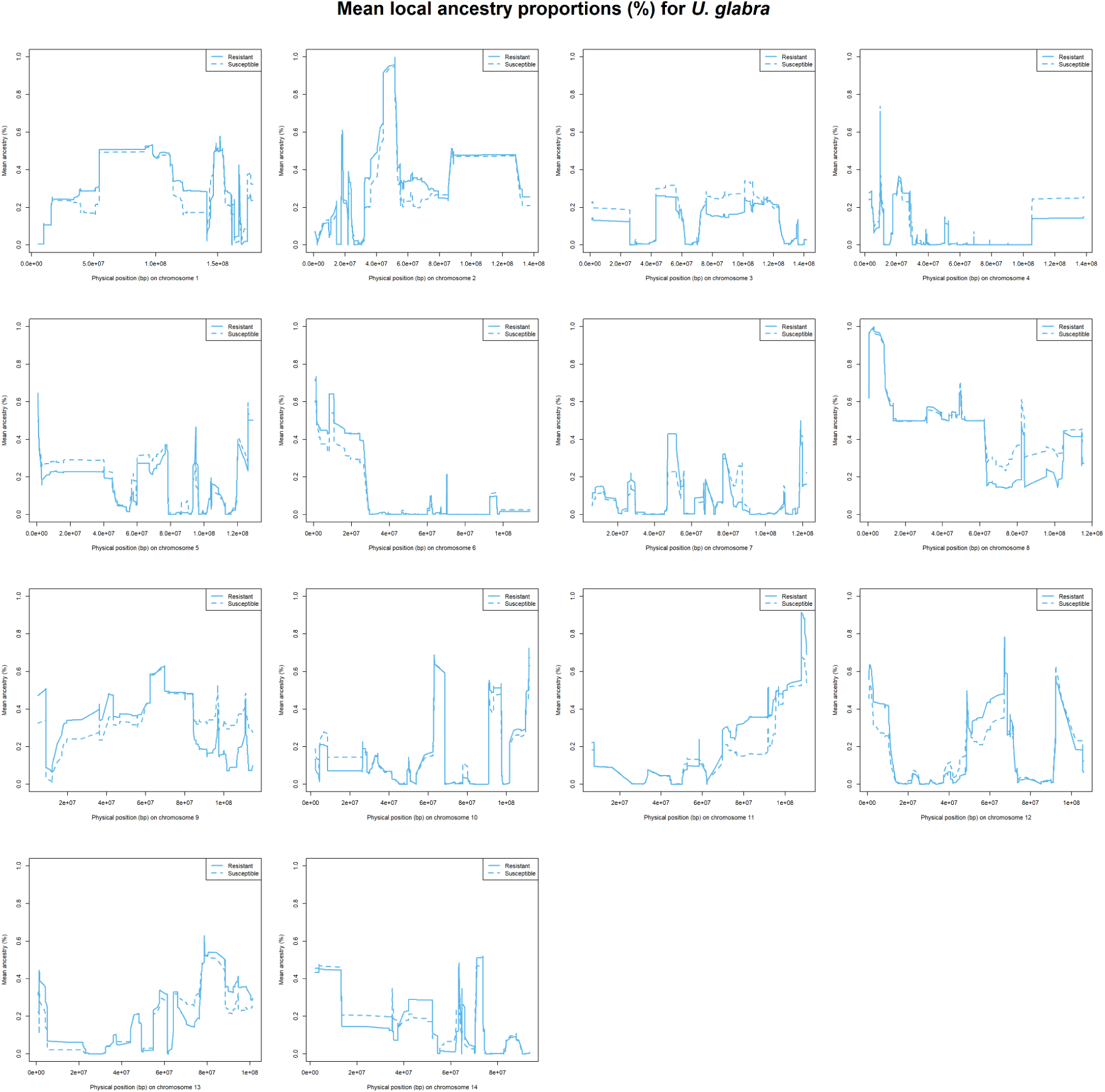
Mean local ancestry proportions (%) for *U. glabra*, comparing resistant WxTM progeny (solid line) with susceptible WxTM progeny (dashed line), based on a threshold of ≤25% defoliation, 8 weeks post-inoculation. Defoliation measurements were only available for 51/60 of the WxTM progeny during this timeframe.

**Figure S8.**
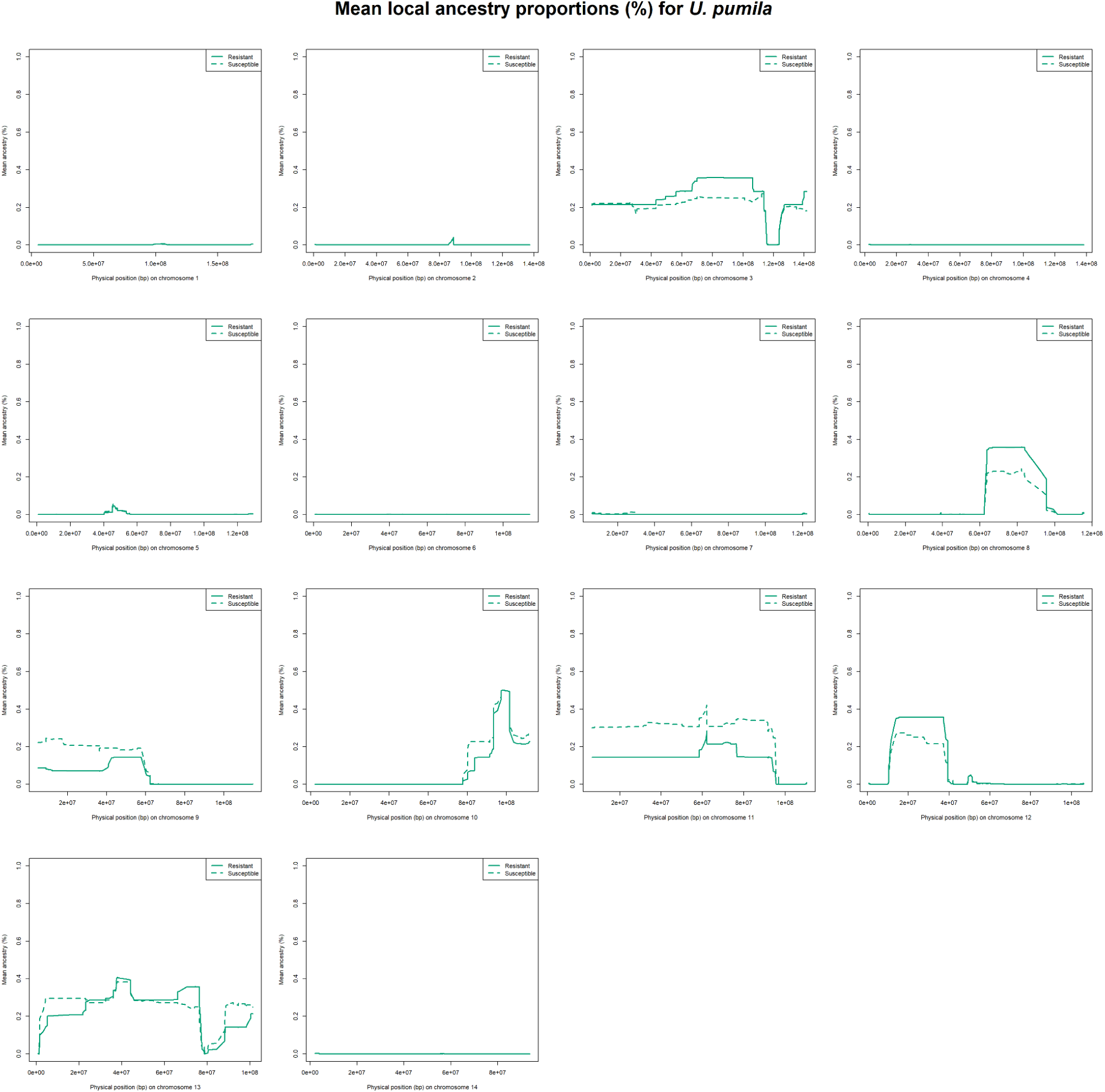
Mean local ancestry proportions (%) for *U. pumila*, comparing resistant WxTM progeny (solid line) with susceptible WxTM progeny (dashed line), based on a threshold of ≤25% defoliation, 8 weeks post-inoculation. Defoliation measurements were only available for 51/60 of the WxTM progeny during this timeframe.

**Figure S9.**
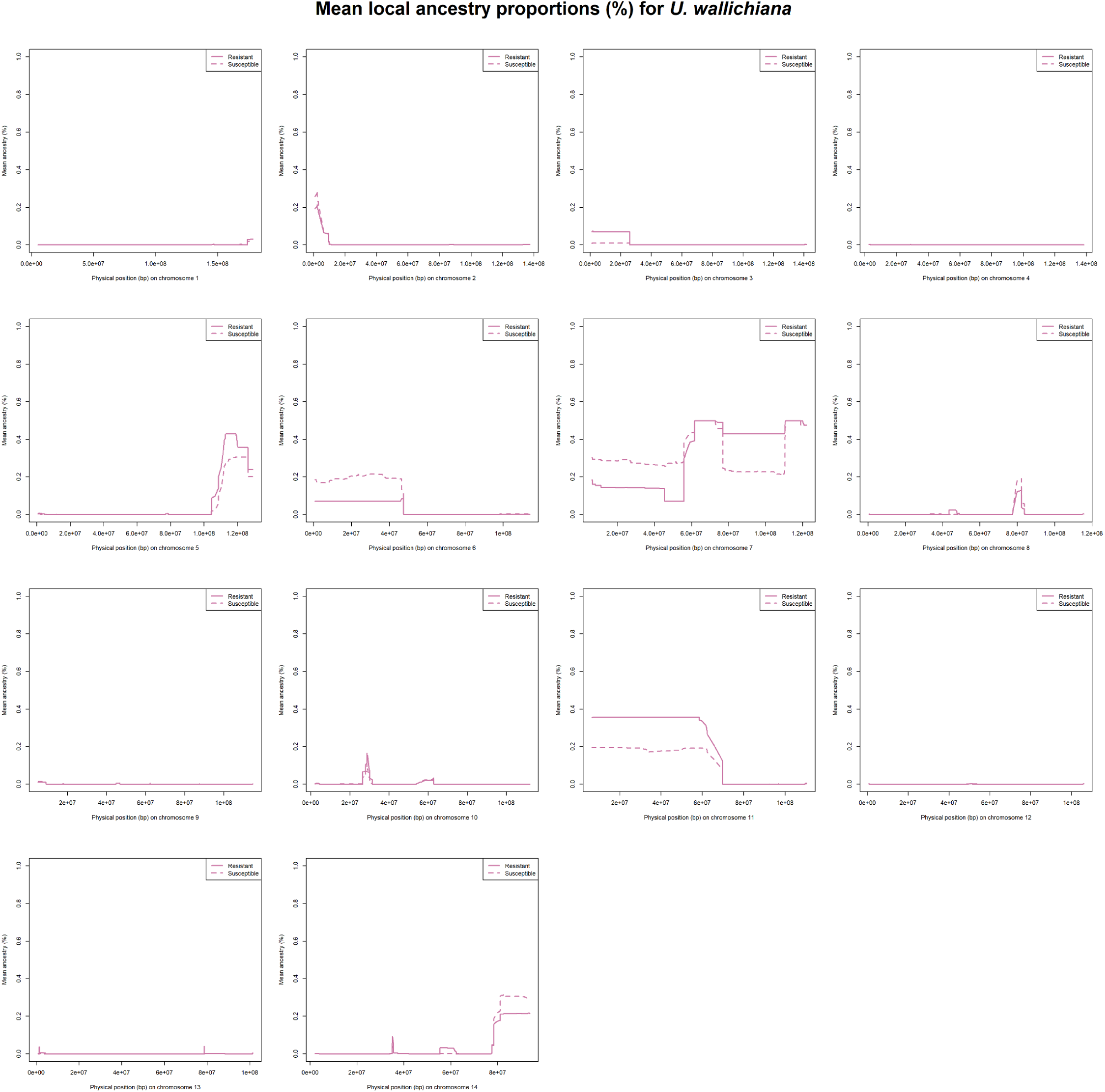
Mean local ancestry proportions (%) for *U. wallichiana*, comparing resistant WxTM progeny (solid line) with susceptible WxTM progeny (dashed line), based on a threshold of ≤25% defoliation, 8 weeks post-inoculation. Defoliation measurements were only available for 51/60 of the WxTM progeny during this timeframe.

**Figure S10.**
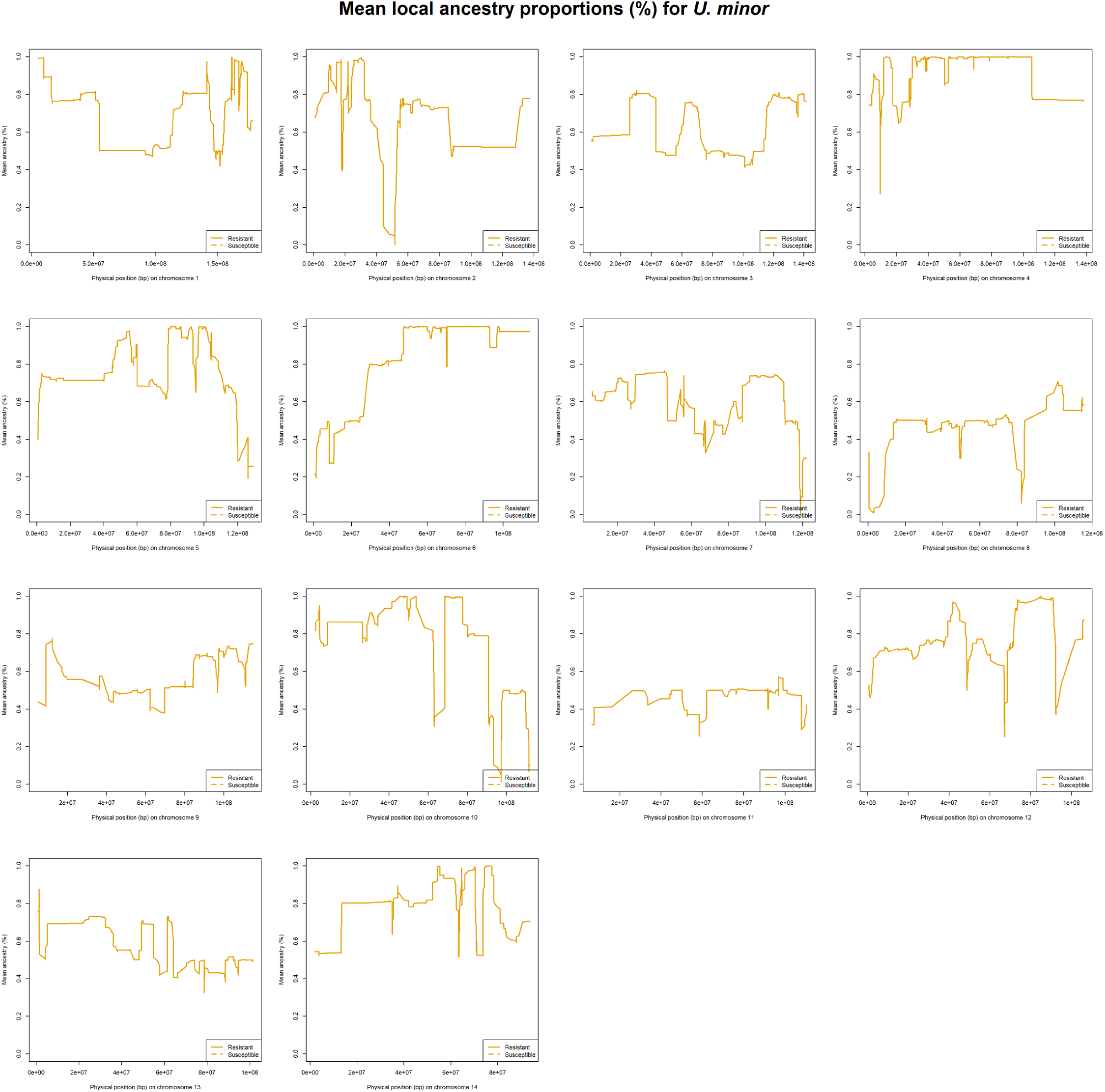
Mean local ancestry proportions (%) for *U. minor*, comparing resistant WxTM progeny (solid line) with susceptible WxTM progeny (dashed line), based on a threshold of <70% defoliation, 4 weeks post-inoculation. Defoliation measurements were only available for 50/60 of the WxTM progeny during this timeframe.

**Figure S11.**
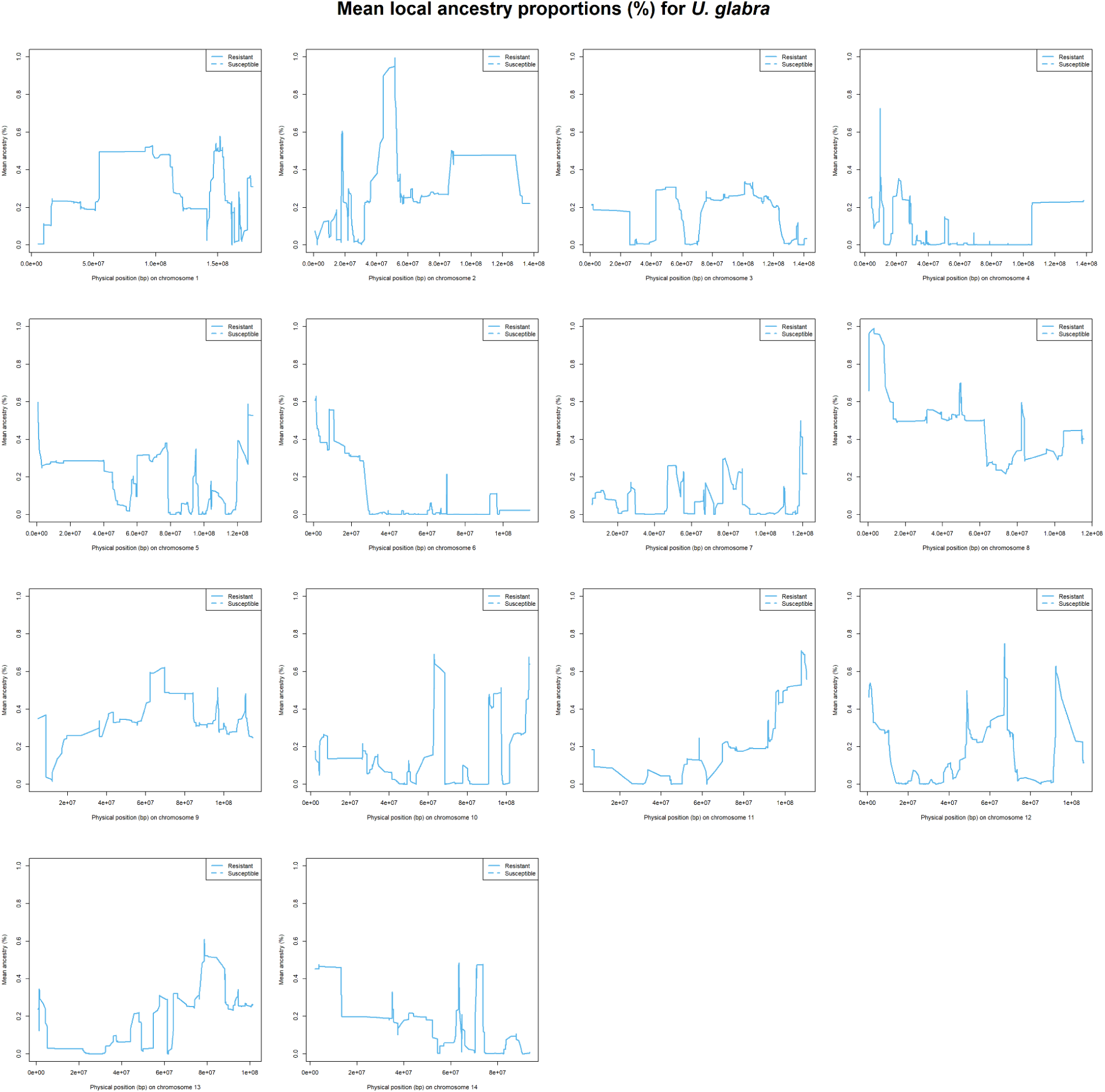
Mean local ancestry proportions (%) for *U. glabra*, comparing resistant WxTM progeny (solid line) with susceptible WxTM progeny (dashed line), based on a threshold of <70% defoliation, 4 weeks post-inoculation. Defoliation measurements were only available for 50/60 of the WxTM progeny during this timeframe.

**Figure S12.**
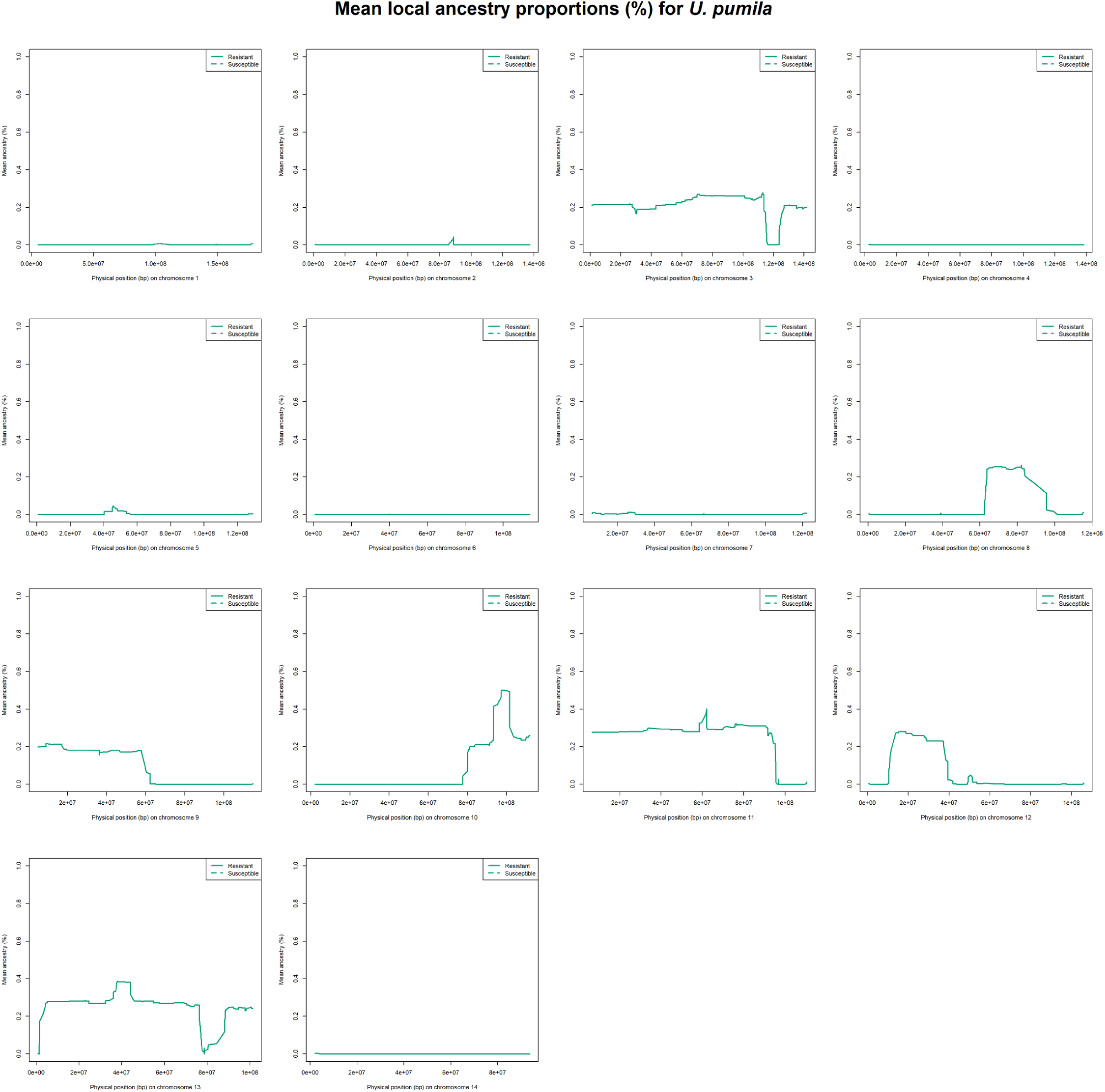
Mean local ancestry proportions (%) for *U. pumila*, comparing resistant WxTM progeny (solid line) with susceptible WxTM progeny (dashed line), based on a threshold of <70% defoliation, 4 weeks post-inoculation. Defoliation measurements were only available for 50/60 of the WxTM progeny during this timeframe.

**Figure S13.**
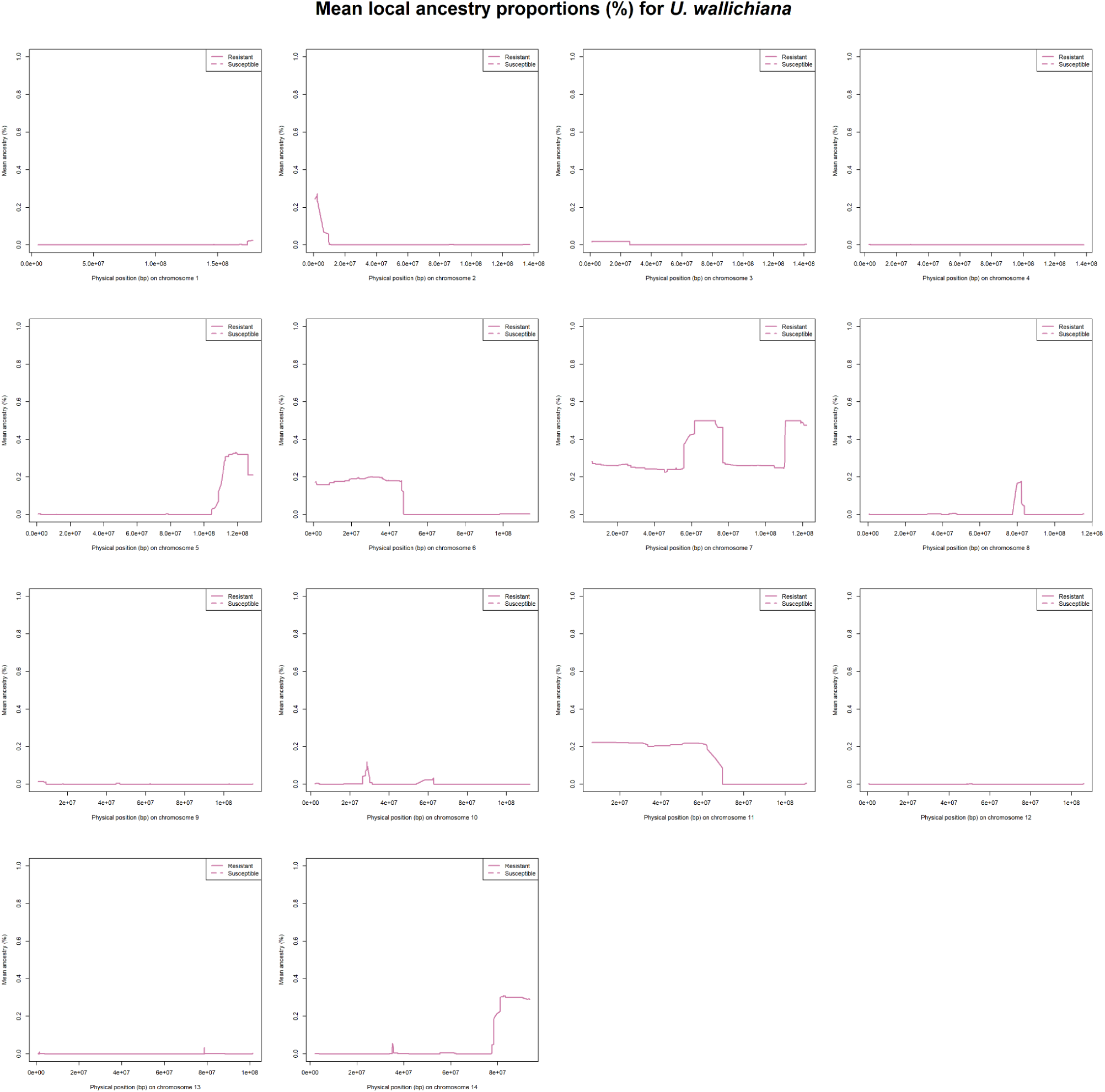
Mean local ancestry proportions (%) for *U. wallichiana*, comparing resistant WxTM progeny (solid line) with susceptible WxTM progeny (dashed line), based on a threshold of <70% defoliation, 4 weeks post-inoculation. Defoliation measurements were only available for 50/60 of the WxTM progeny during this timeframe.

**Figure S14.**
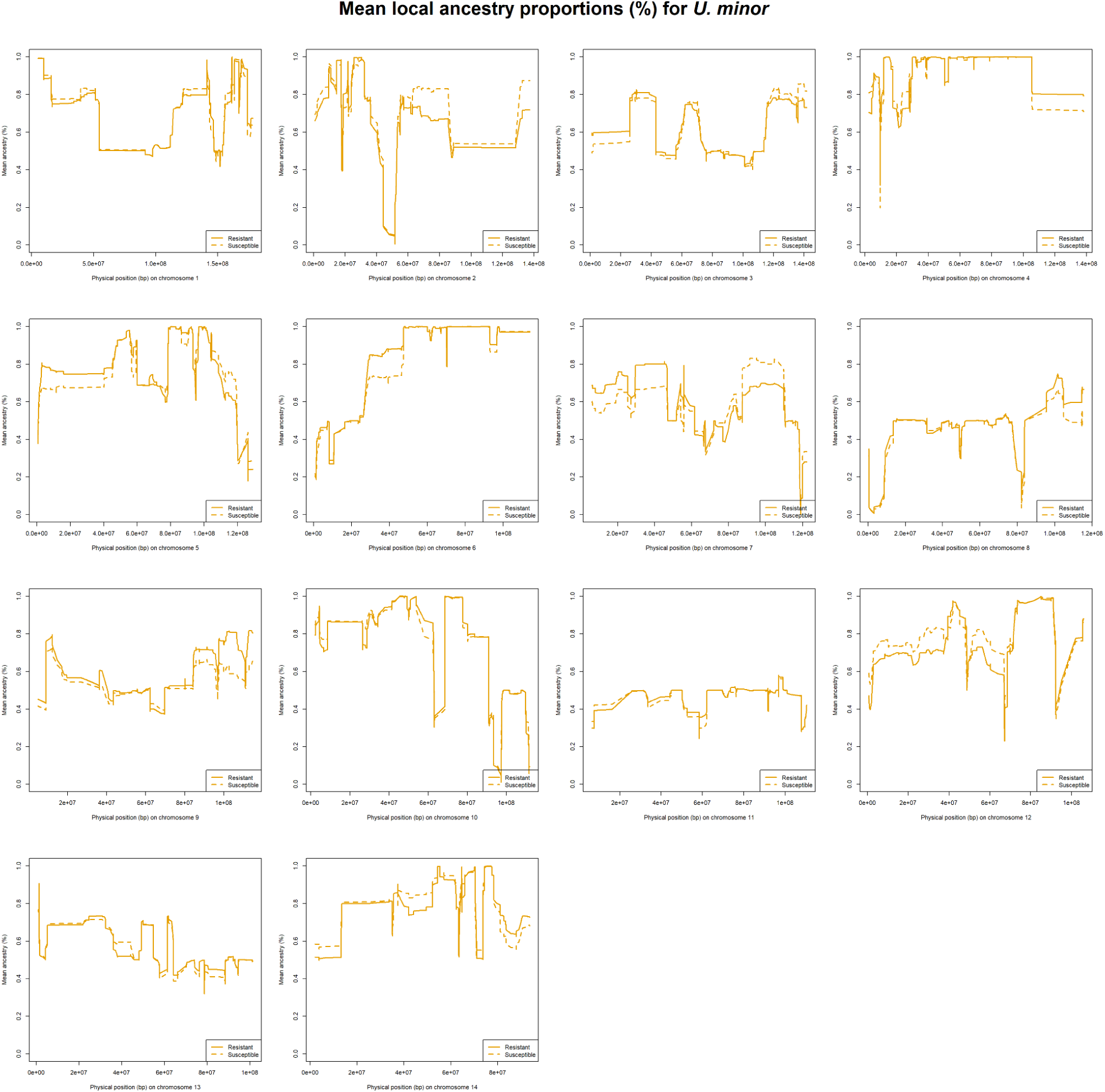
Mean local ancestry proportions (%) for *U. minor*, comparing resistant WxTM progeny (solid line) with susceptible WxTM progeny (dashed line), based on a threshold of <70% defoliation, 8 weeks post-inoculation. Defoliation measurements were only available for 51/60 of the WxTM progeny during this timeframe.

**Figure S15.**
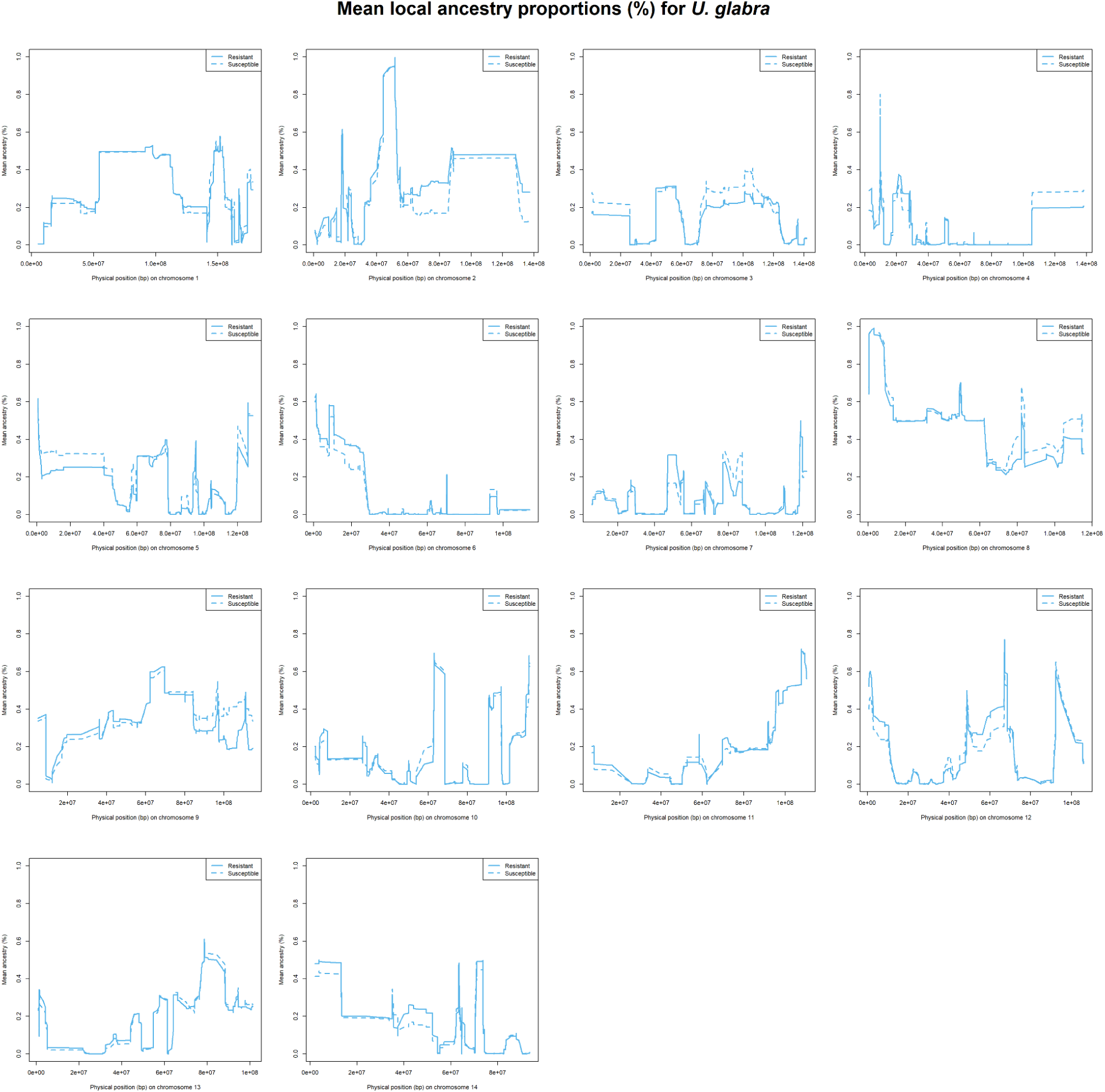
Mean local ancestry proportions (%) for *U. glabra*, comparing resistant WxTM progeny (solid line) with susceptible WxTM progeny (dashed line), based on a threshold of <70% defoliation, 8 weeks post-inoculation. Defoliation measurements were only available for 51/60 of the WxTM progeny during this timeframe.

**Figure S16.**
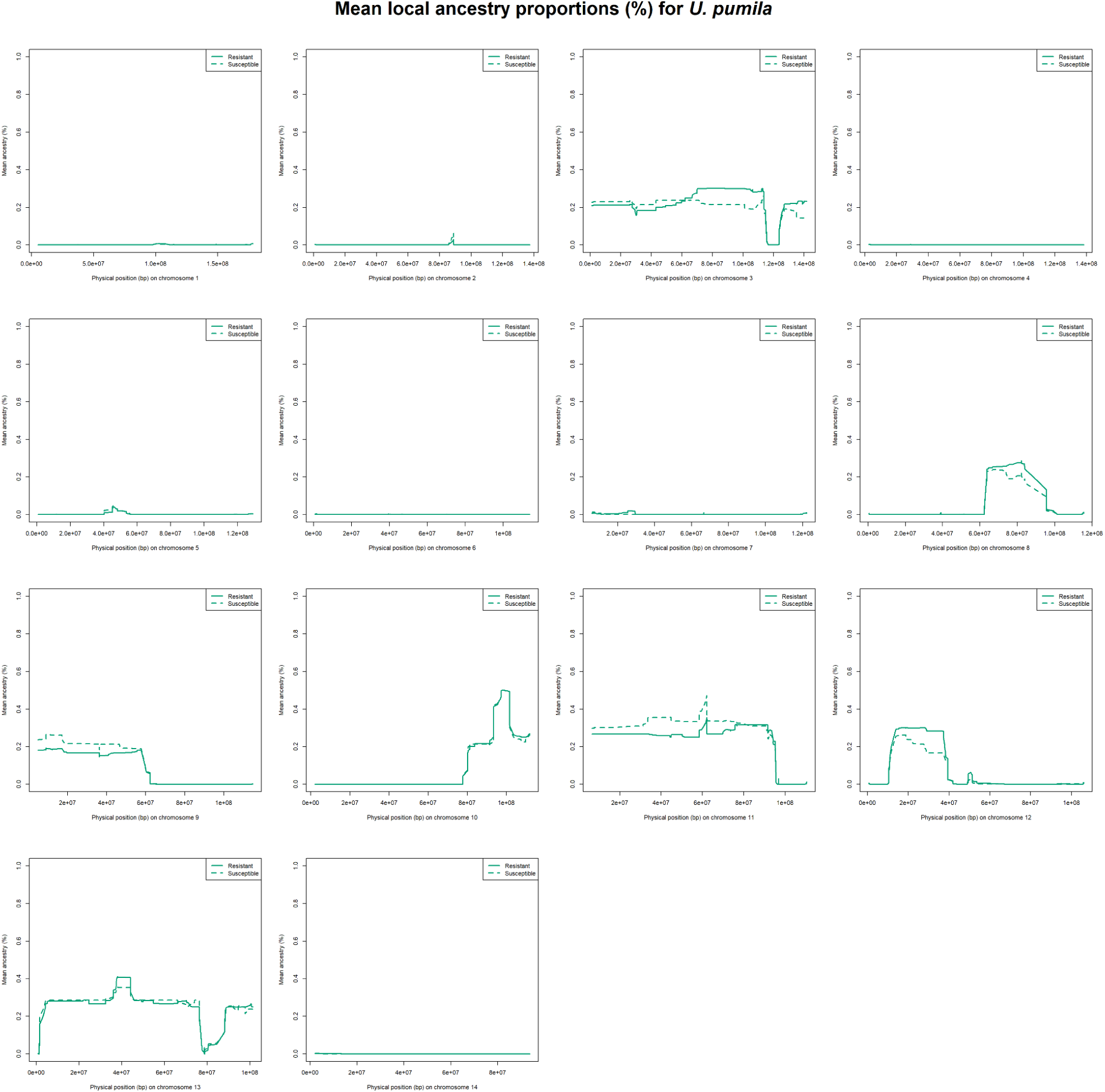
Mean local ancestry proportions (%) for *U. pumila*, comparing resistant WxTM progeny (solid line) with susceptible WxTM progeny (dashed line), based on a threshold of <70% defoliation, 8 weeks post-inoculation. Defoliation measurements were only available for 51/60 of the WxTM progeny during this timeframe.

**Figure S17.**
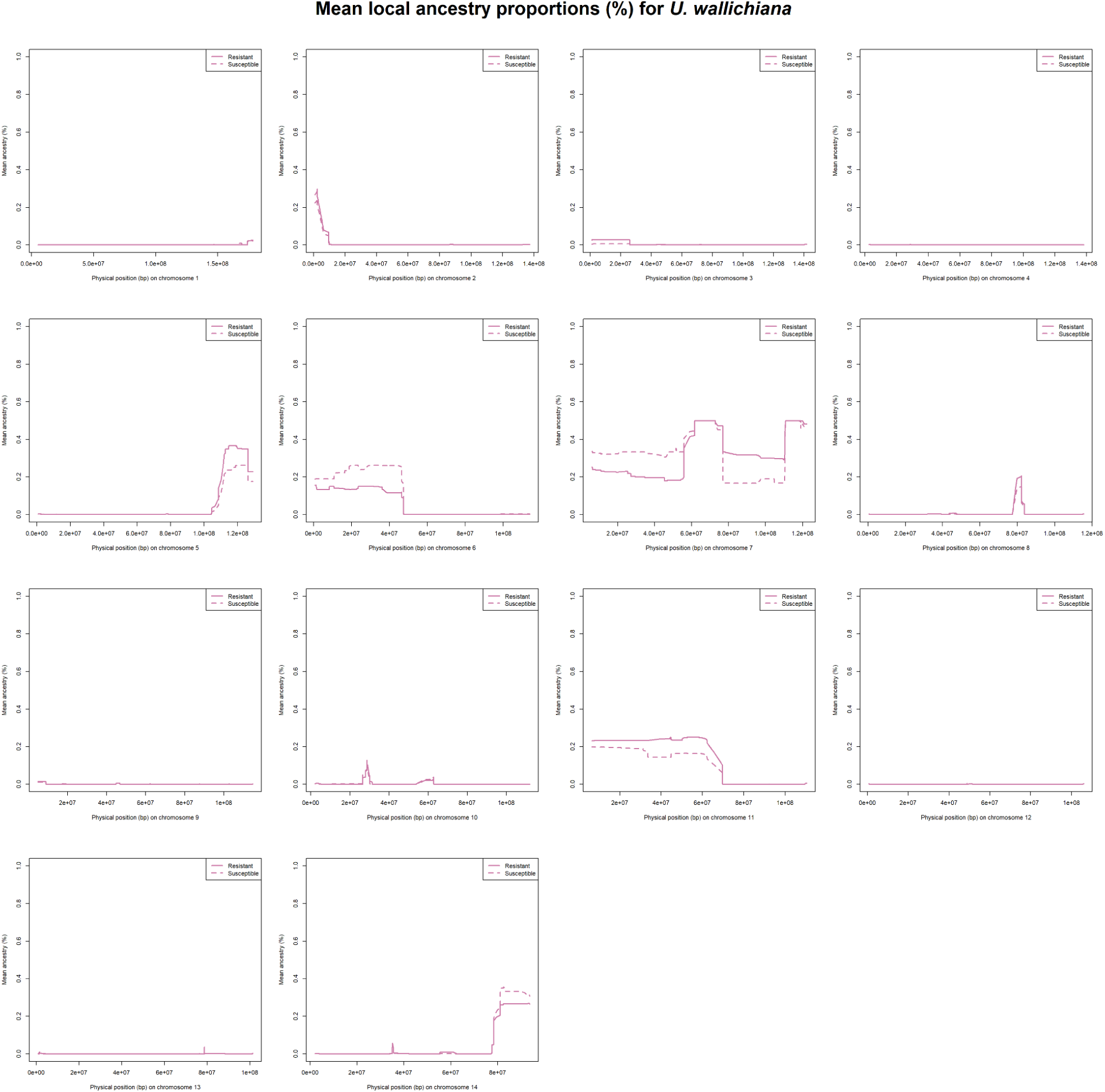
Mean local ancestry proportions (%) for *U. wallichiana*, comparing resistant WxTM progeny (solid line) with susceptible WxTM progeny (dashed line), based on a threshold of <70% defoliation, 8 weeks post-inoculation. Defoliation measurements were only available for 51/60 of the WxTM progeny during this timeframe.

**Figure S18.**
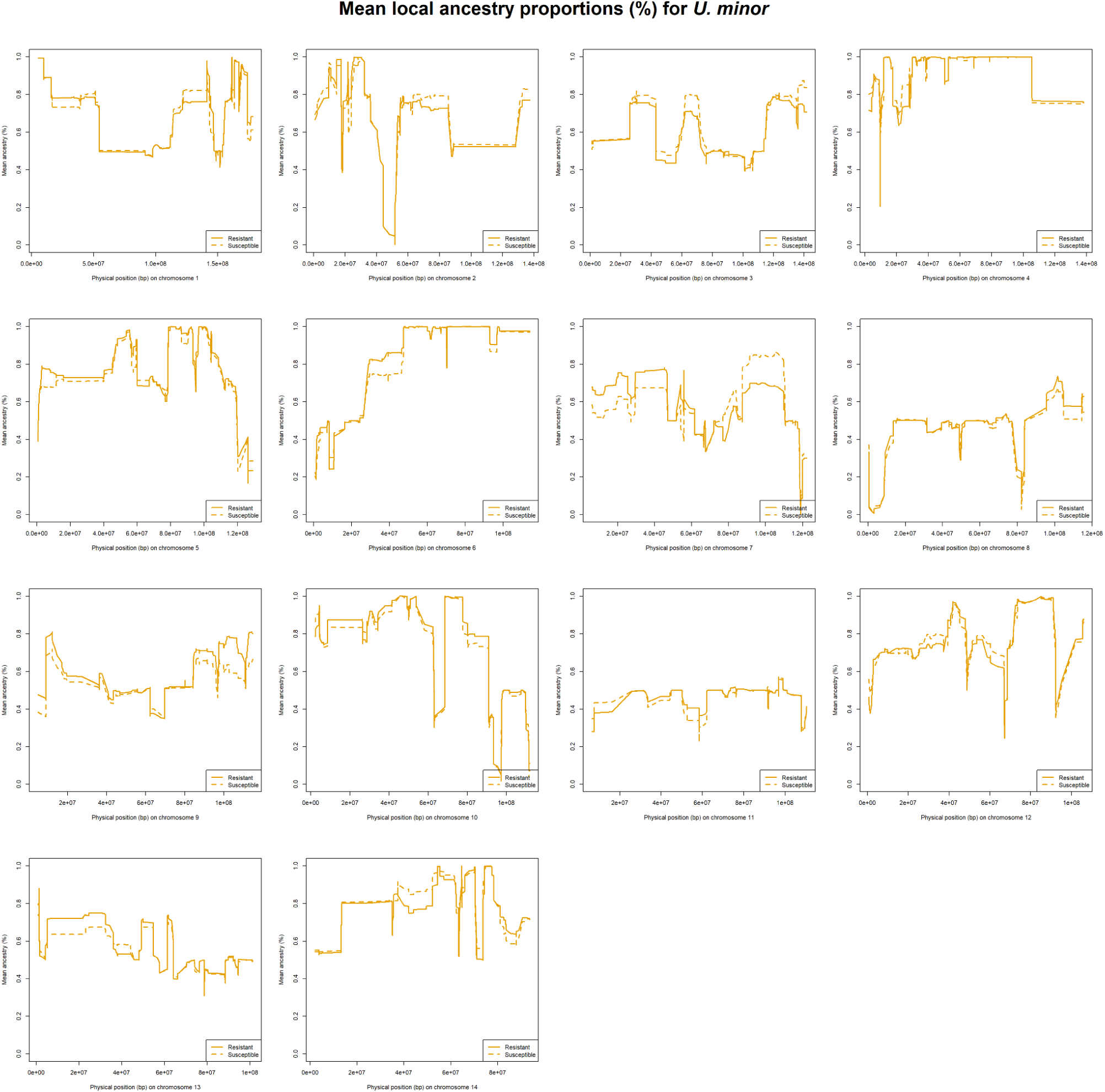
Mean local ancestry proportions (%) for *U. minor*, comparing resistant WxTM progeny (solid line) with susceptible WxTM progeny (dashed line), based on a threshold of <70% defoliation, 12 weeks post-inoculation.

**Figure S19.**
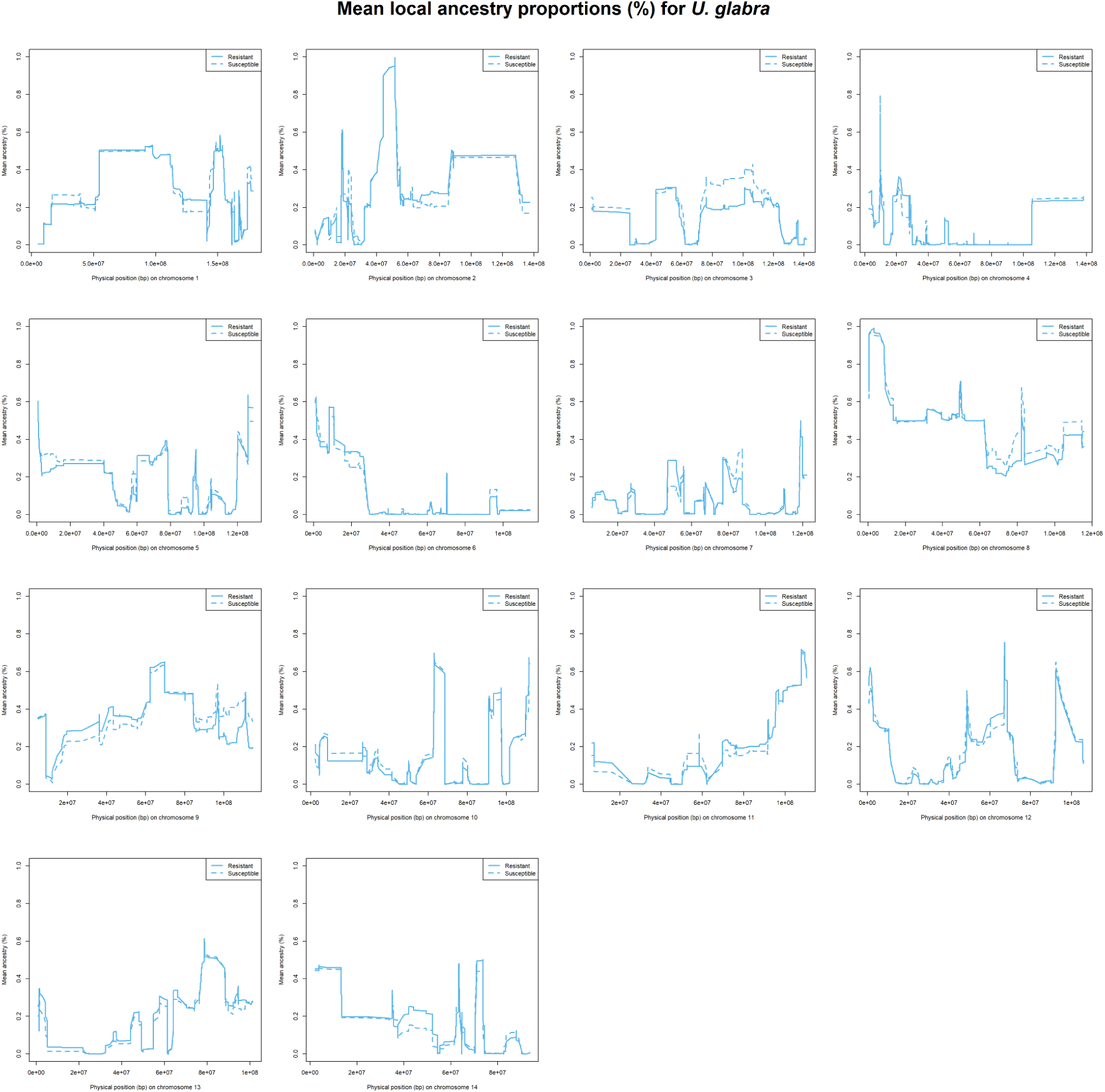
Mean local ancestry proportions (%) for *U. glabra*, comparing resistant WxTM progeny (solid line) with susceptible WxTM progeny (dashed line), based on a threshold of <70% defoliation, 12 weeks post-inoculation.

**Figure S20.**
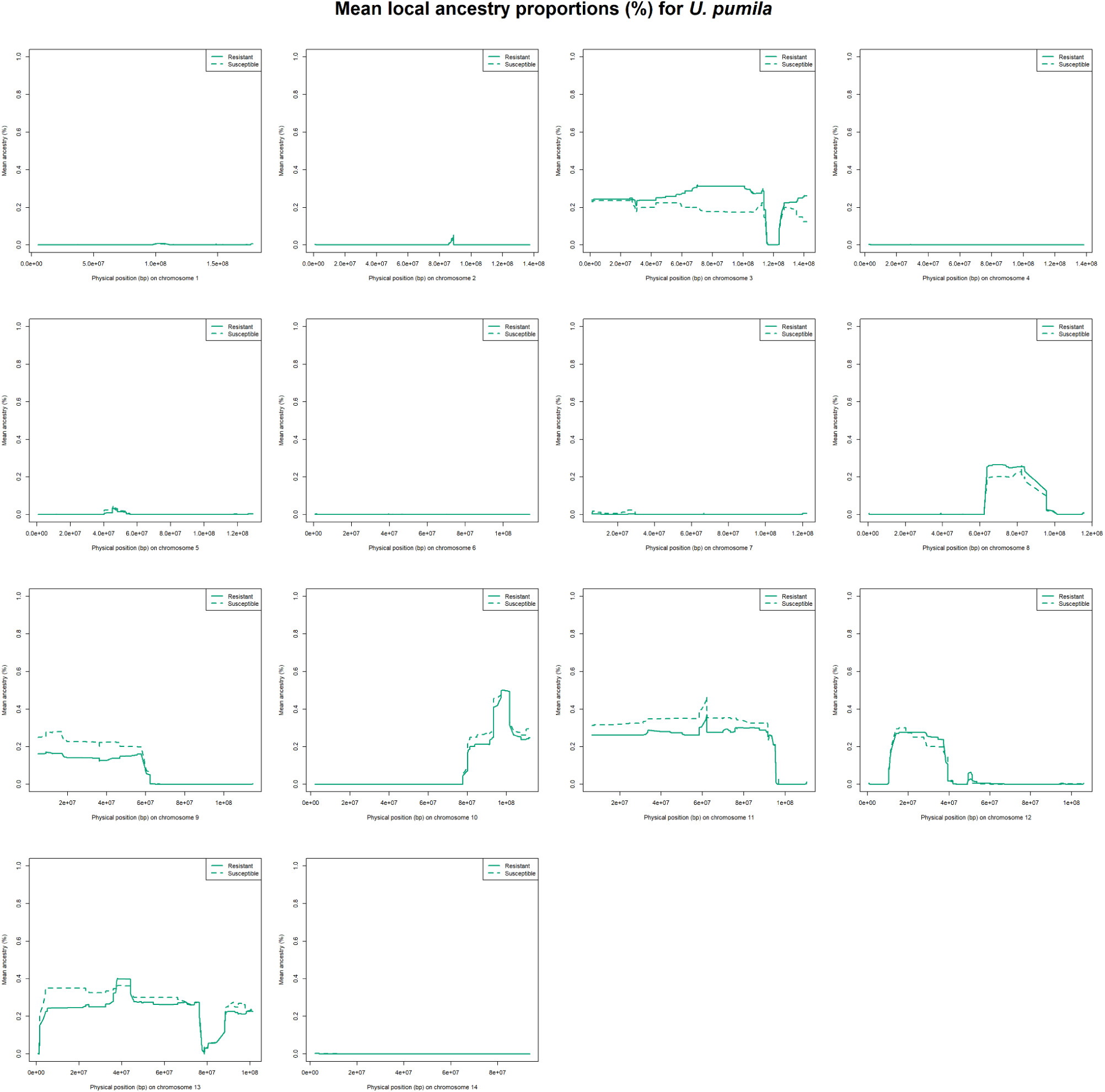
Mean local ancestry proportions (%) for *U. pumila*, comparing resistant WxTM progeny (solid line) with susceptible WxTM progeny (dashed line), based on a threshold of <70% defoliation, 12 weeks post-inoculation.

**Figure S21.**
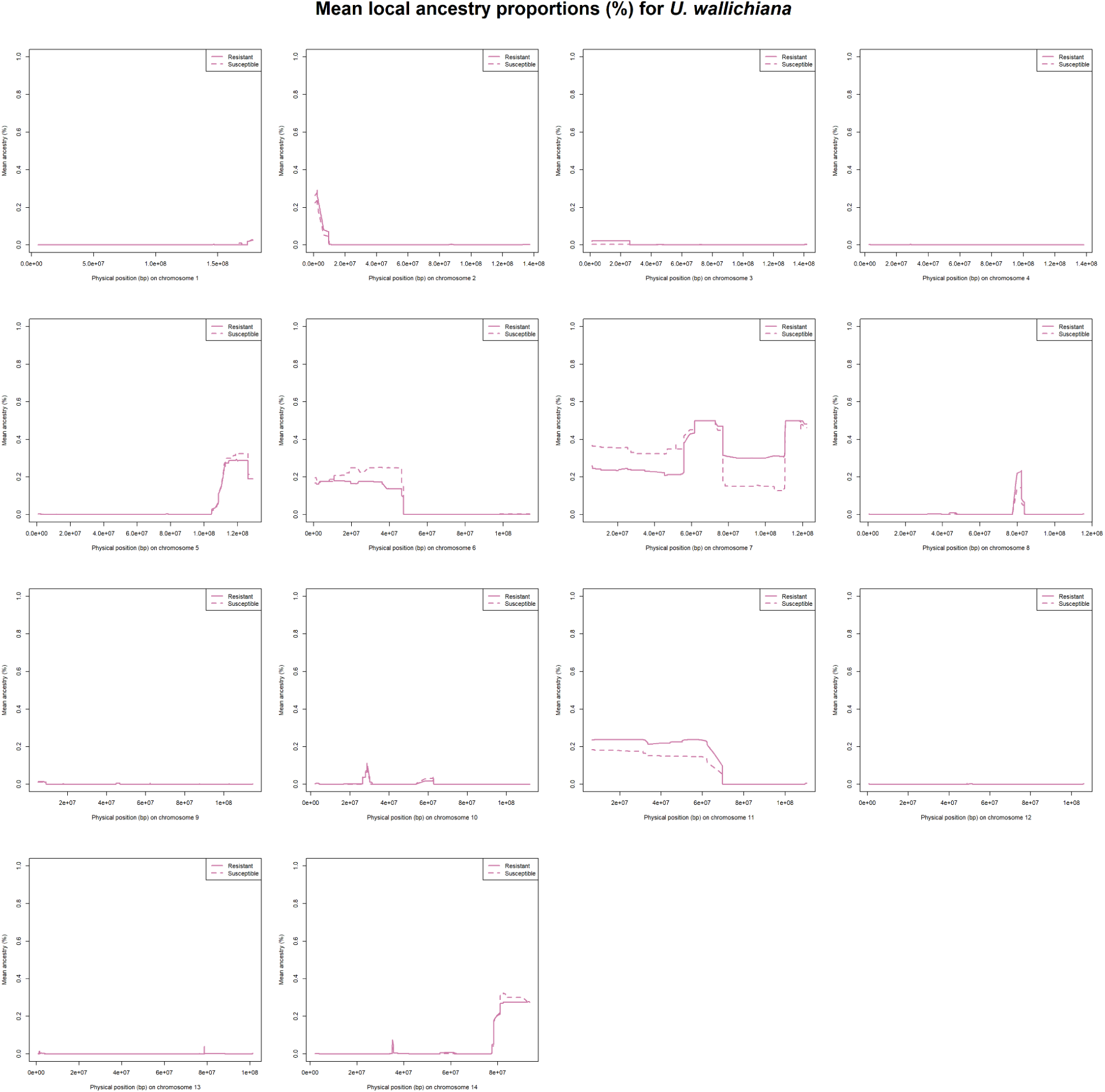
Mean local ancestry proportions (%) for *U. wallichiana*, comparing resistant WxTM progeny (solid line) with susceptible WxTM progeny (dashed line), based on a threshold of <70% defoliation, 12 weeks post-inoculation.

**Figure S22.**
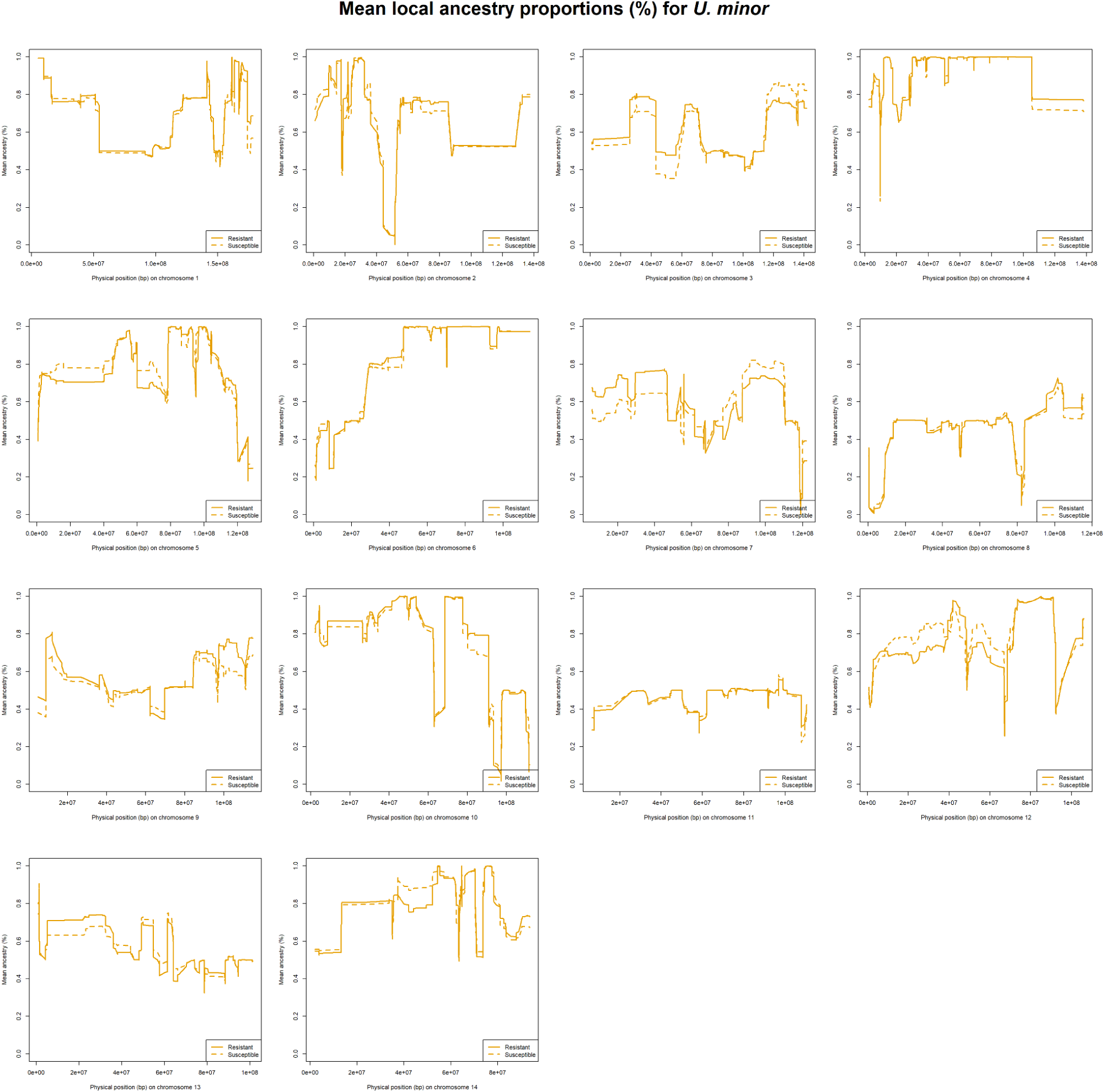
Mean local ancestry proportions (%) for *U. minor*, comparing resistant WxTM progeny (solid line) with susceptible WxTM progeny (dashed line), based on a threshold of <70% defoliation, 1 year post-inoculation.

**Figure S23.**
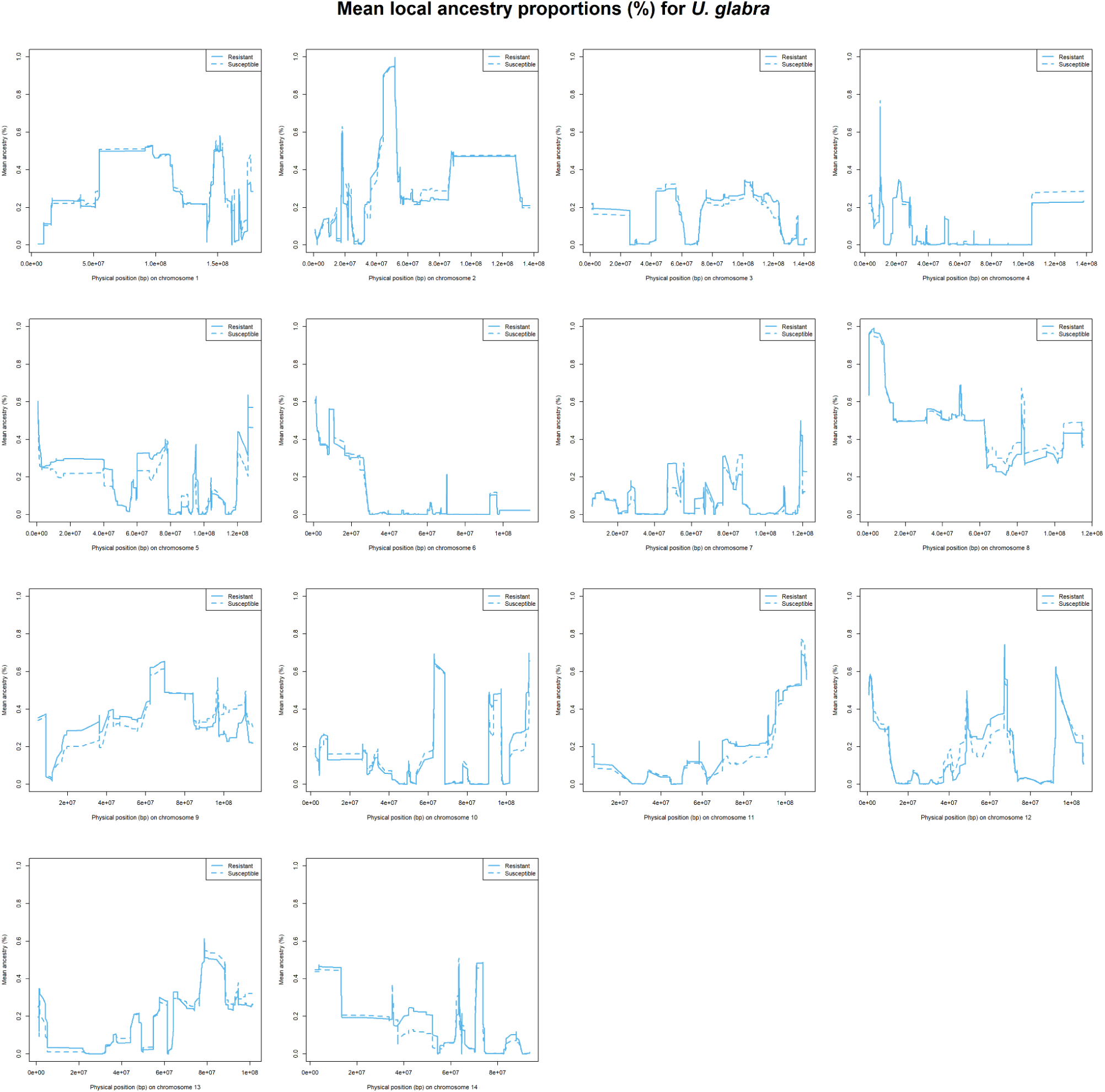
Mean local ancestry proportions (%) for *U. glabra*, comparing resistant WxTM progeny (solid line) with susceptible WxTM progeny (dashed line), based on a threshold of <70% defoliation, 1 year post-inoculation.

**Figure S24.**
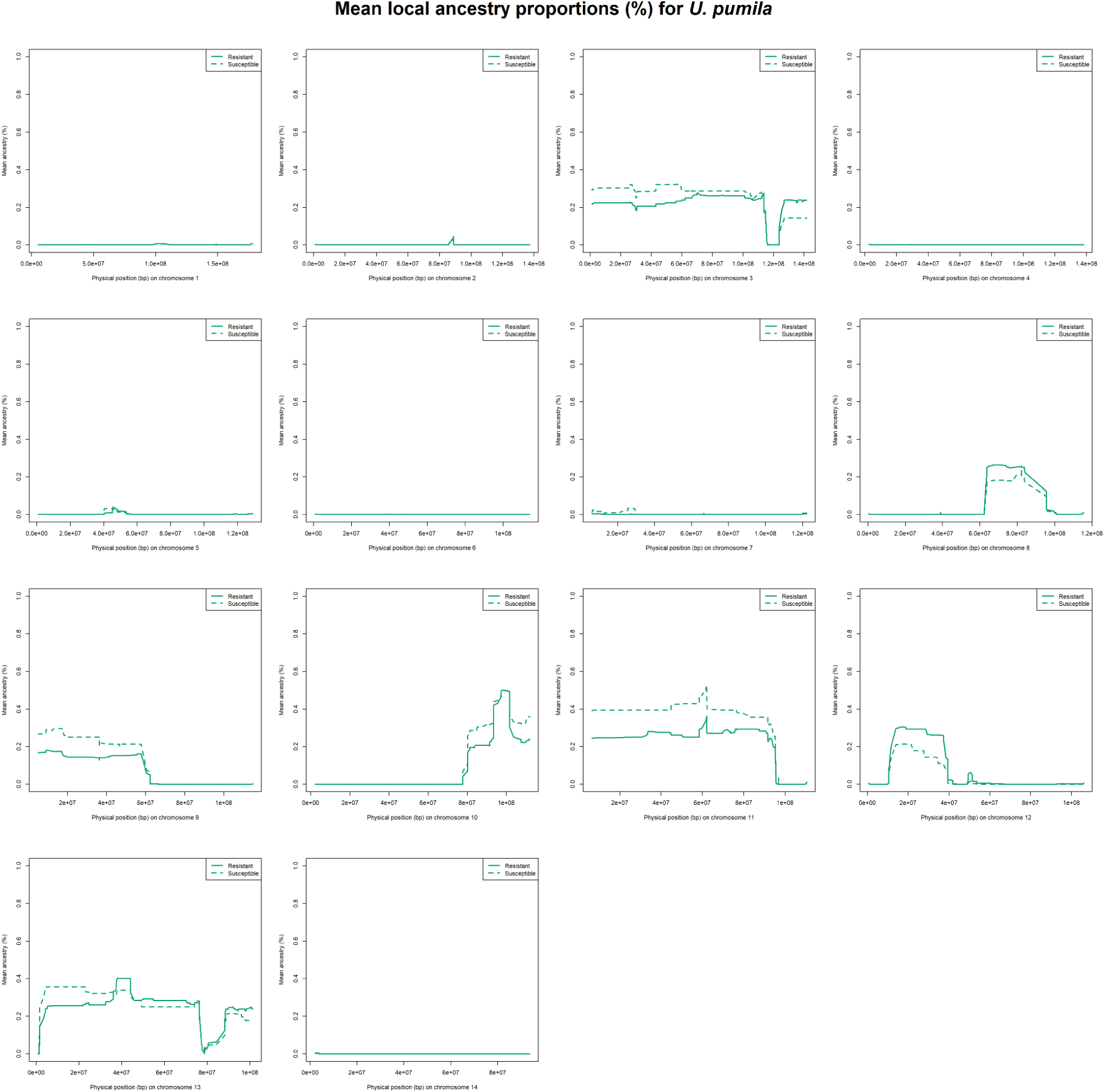
Mean local ancestry proportions (%) for *U. pumila*, comparing resistant WxTM progeny (solid line) with susceptible WxTM progeny (dashed line), based on a threshold of <70% defoliation, 1 year post-inoculation.

**Figure S25.**
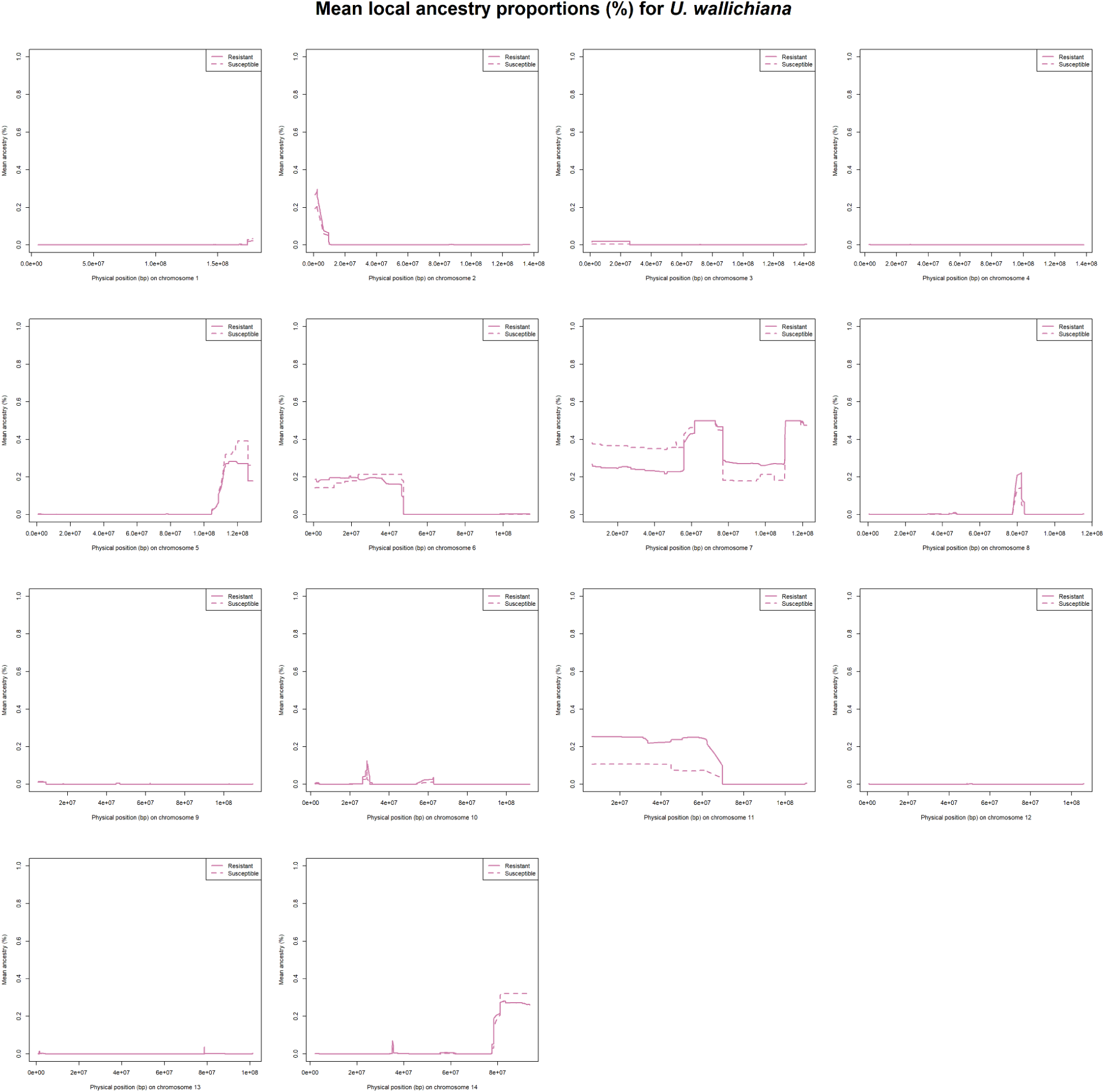
Mean local ancestry proportions (%) for *U. wallichiana*, comparing resistant WxTM progeny (solid line) with susceptible WxTM progeny (dashed line), based on a threshold of <70% defoliation, 1 year post-inoculation.

**Figure S26.**
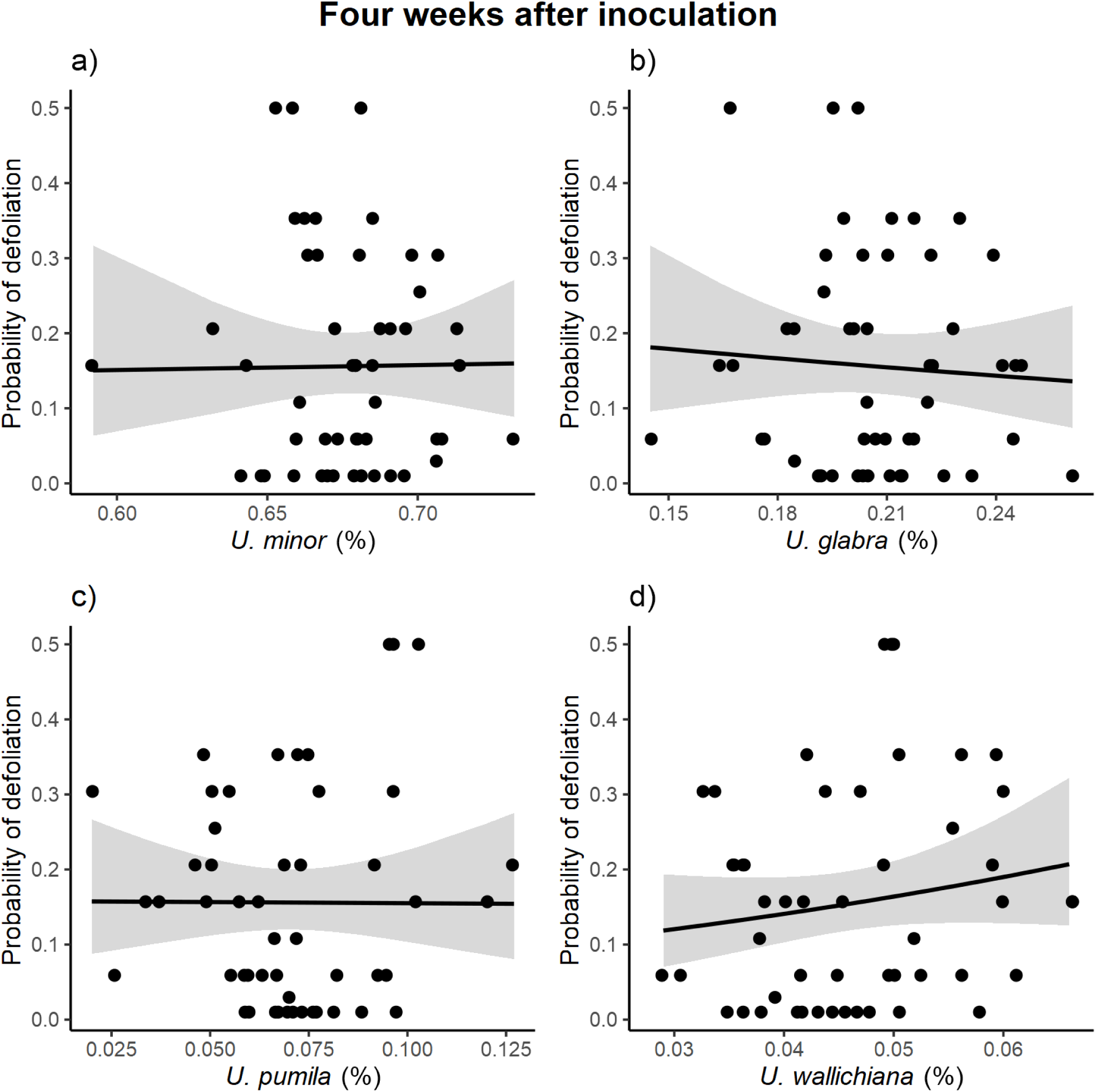
Relationship between the probability of defoliation at week 4 following inoculation and the % of global ancestry for all 4 parental species in the WxTM progeny (n = 50). Lines are the slopes of beta regressions, and shades the confidence intervals. Global ancestry refers to the % of the genome ascribed to a parental species following admixture analysis (see Results).

**Figure S27.**
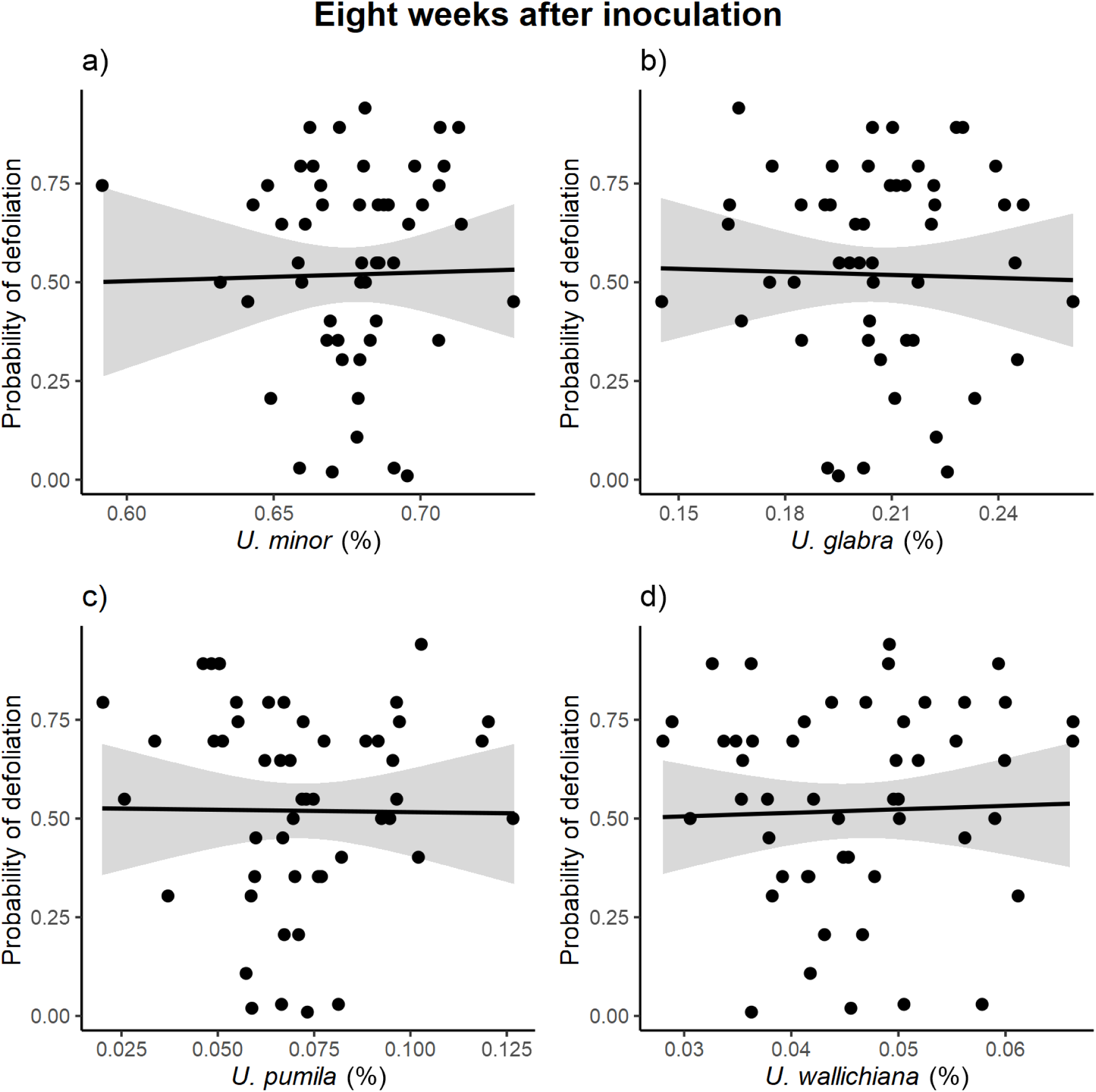
Relationship between the probability of defoliation at week 8 following inoculation and the % of global ancestry for all 4 parental species in the WxTM progeny (n = 51). Lines are the slopes of beta regressions, and shades the confidence intervals. Global ancestry refers to the % of the genome ascribed to a parental species following admixture analysis (see Results).

**Figure S28.**
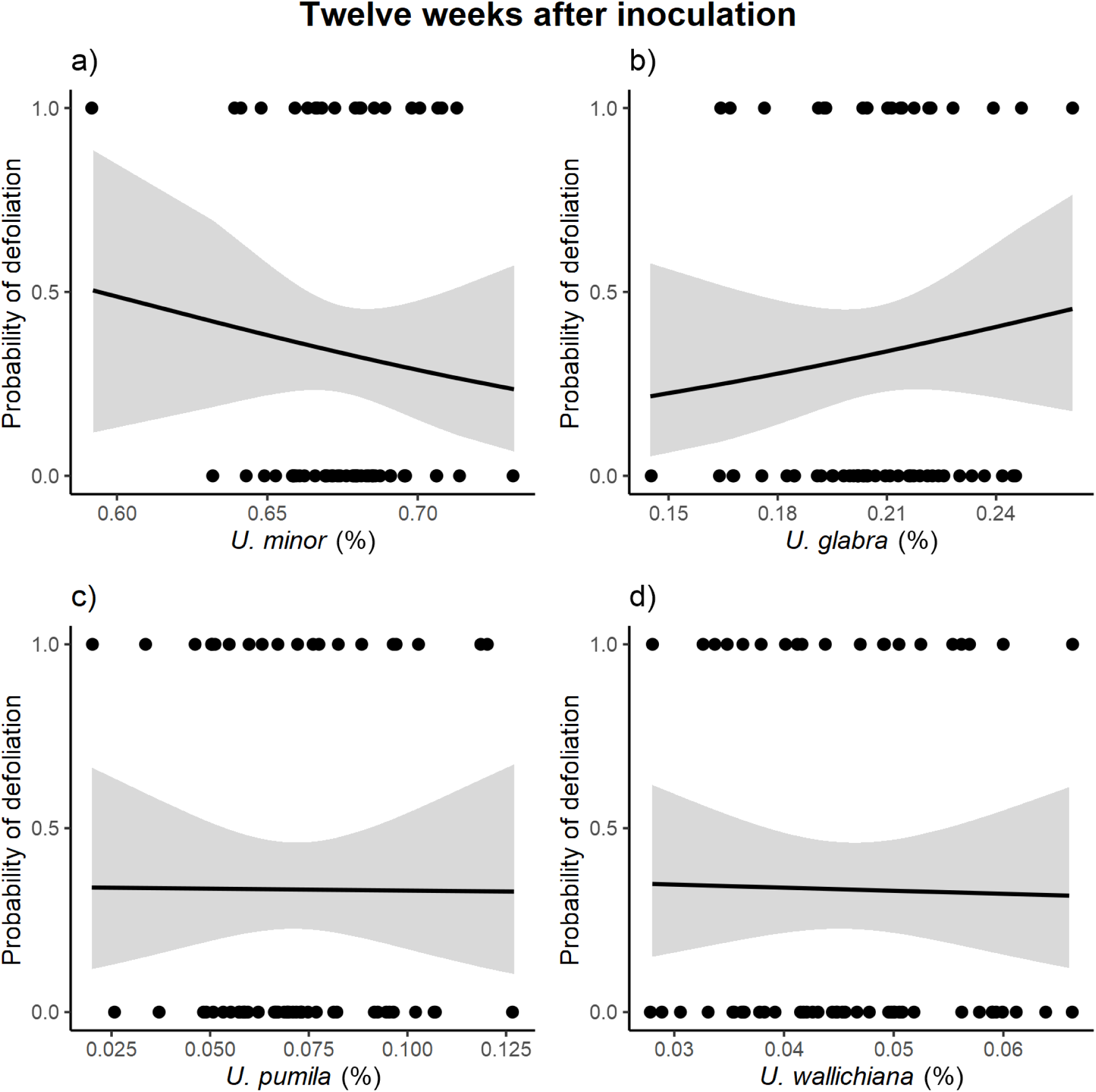
Relationship between the probability of defoliation at week 12 following inoculation and the % of global ancestry for all 4 parental species in the WxTM progeny (n = 60). Lines are the slopes of logistic regressions, and shades the confidence intervals. Note: probability of 1 refers to defoliation >= 70%, whereas 0 = <70%. Global ancestry refers to the % of the genome ascribed to a parental species following admixture analysis (see Results).

**Figure S29.**
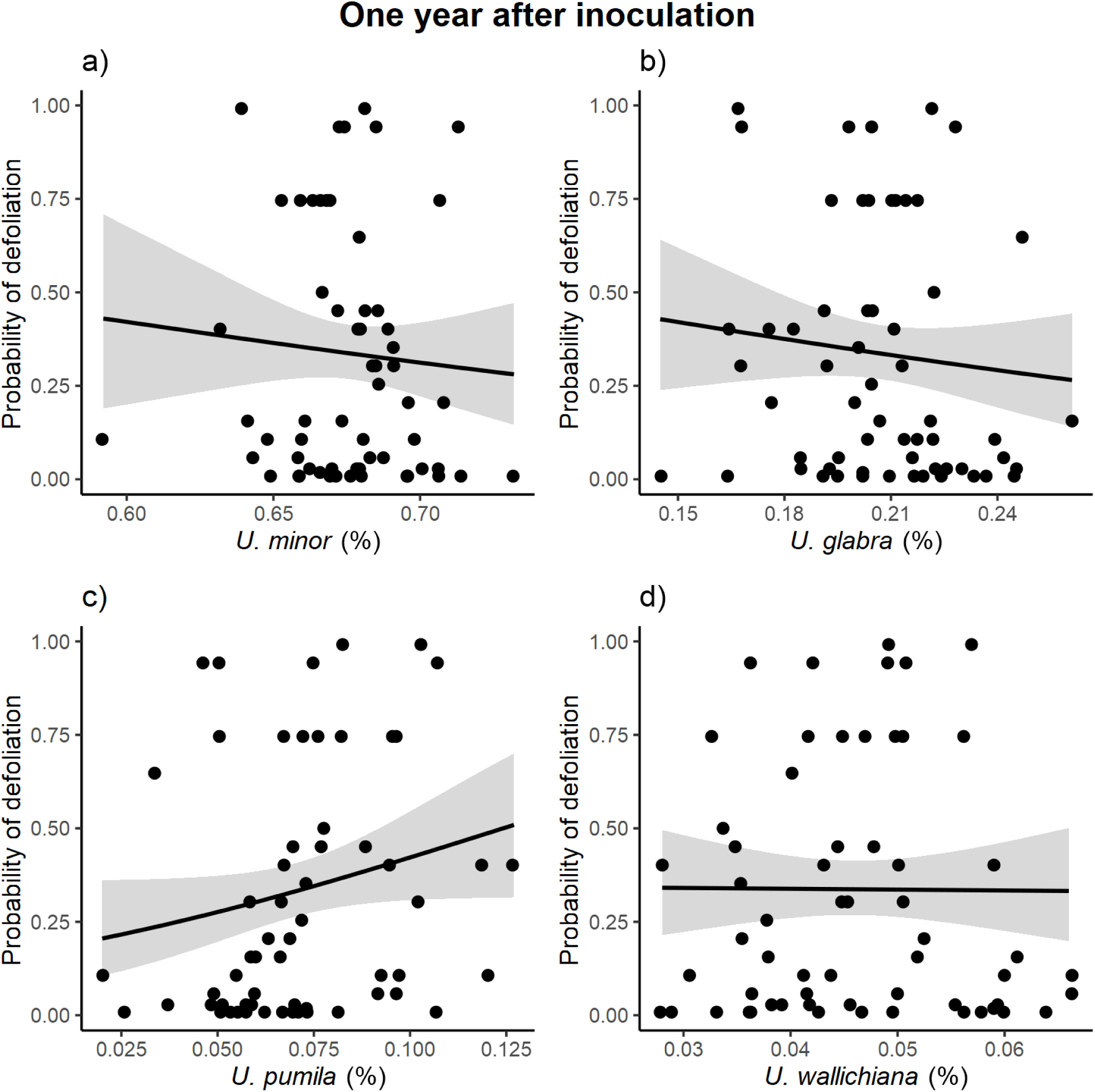
Relationship between the probability of defoliation at year 1 following inoculation and the % of global ancestry for all 4 parental species in the WxTM progeny (n = 60). Lines are the slopes of beta regressions, and shades the confidence intervals. Global ancestry refers to the % of the genome ascribed to a parental species following admixture analysis (see Results).

